# Elicitation of potent neutralizing antibody responses by designed protein nanoparticle vaccines for SARS-CoV-2

**DOI:** 10.1101/2020.08.11.247395

**Authors:** Alexandra C. Walls, Brooke Fiala, Alexandra Schäfer, Samuel Wrenn, Minh N. Pham, Michael Murphy, Longping V. Tse, Laila Shehata, Megan A. O'Connor, Chengbo Chen, Mary Jane Navarro, Marcos C. Miranda, Deleah Pettie, Rashmi Ravichandran, John C. Kraft, Cassandra Ogohara, Anne Palser, Sara Chalk, E-Chiang Lee, Elizabeth Kepl, Cameron M. Chow, Claire Sydeman, Edgar A. Hodge, Brieann Brown, Jim T. Fuller, Kenneth H. Dinnon, Lisa E. Gralinski, Sarah R. Leist, Kendra L. Gully, Thomas B. Lewis, Miklos Guttman, Helen Y. Chu, Kelly K. Lee, Deborah H. Fuller, Ralph S. Baric, Paul Kellam, Lauren Carter, Marion Pepper, Timothy P. Sheahan, David Veesler, Neil P. King

## Abstract

A safe, effective, and scalable vaccine is urgently needed to halt the ongoing SARS-CoV-2 pandemic. Here, we describe the structure-based design of self-assembling protein nanoparticle immunogens that elicit potent and protective antibody responses against SARS-CoV-2 in mice. The nanoparticle vaccines display 60 copies of the SARS-CoV-2 spike (S) glycoprotein receptor-binding domain (RBD) in a highly immunogenic array and induce neutralizing antibody titers roughly ten-fold higher than the prefusion-stabilized S ectodomain trimer despite a more than five-fold lower dose. Antibodies elicited by the nanoparticle immunogens target multiple distinct epitopes on the RBD, suggesting that they may not be easily susceptible to escape mutations, and exhibit a significantly lower binding:neutralizing ratio than convalescent human sera, which may minimize the risk of vaccine-associated enhanced respiratory disease. The high yield and stability of the protein components and assembled nanoparticles, especially compared to the SARS-CoV-2 prefusion-stabilized S trimer, suggest that manufacture of the nanoparticle vaccines will be highly scalable. These results highlight the utility of robust antigen display platforms for inducing potent neutralizing antibody responses and have launched cGMP manufacturing efforts to advance the lead RBD nanoparticle vaccine into the clinic.

## INTRODUCTION

The recent emergence of a previously unknown virus in Wuhan, China has resulted in the ongoing COVID-19 pandemic that has caused more than 18,700,000 infections and 700,000 fatalities as of August 6, 2020 (WHO). Rapid viral isolation and sequencing revealed by January 2020 that the newly emerged zoonotic pathogen was a coronavirus closely related to SARS-CoV and was therefore named SARS-CoV-2 (Zhou et al., 2020c; Zhu et al., 2020b). SARS-CoV-2 is believed to have originated in bats based on the isolation of the closely related RaTG13 virus from *Rhinolophus affinis* (Zhou et al., 2020c) and the identification of the RmYN02 genome sequence in metagenomics analyses of *Rhinolophus malayanus* (Zhou et al., 2020b), both from Yunnan, China.

Similar to other coronaviruses, SARS-CoV-2 entry into host cells is mediated by the transmembrane spike (S) glycoprotein, which forms prominent homotrimers protruding from the viral surface (Tortorici and Veesler, 2019; Walls et al., 2016a; Walls et al., 2017). Cryo-electron microscopy structures of SARS-CoV-2 S revealed its shared architecture with SARS-CoV S and provided a blueprint for the design of vaccines and antivirals (Walls et al., 2020; Wrapp et al., 2020). Both SARS-CoV-2 S and SARS-CoV S bind to angiotensin-converting enzyme 2 (ACE2), which serves as entry receptor (Hoffmann et al., 2020; Letko et al., 2020; Li et al., 2003; Walls et al., 2020; Wrapp et al., 2020; Zhou et al., 2020c). Structures of the SARS-CoV-2 S receptor-binding domain (RBD) in complex with ACE2 defined key residues involved in recognition and guide surveillance studies aiming to detect the emergence of mutants with altered binding affinity for ACE2 or distinct antigenicity (Lan et al., 2020; Shang et al., 2020; Wang et al., 2020b; Yan et al., 2020).

As the coronavirus S glycoprotein is surface-exposed and initiates infection, it is the main target of neutralizing antibodies (Abs) upon infection and the focus of vaccine design (Tortorici and Veesler, 2019). S trimers are extensively decorated with N-linked glycans that are important for proper folding (Rossen et al., 1998) and for modulating accessibility to host proteases and neutralizing Abs (Walls et al., 2016b; Walls et al., 2019; Watanabe et al., 2020; Xiong et al., 2018; Yang et al., 2015). We previously characterized potent human neutralizing Abs from rare memory B cells of individuals infected with SARS-CoV (Rockx et al., 2008; Traggiai et al., 2004) or MERS-CoV (Corti et al., 2015) in complex with their respective S glycoproteins to provide molecular-level information on the mechanism of competitive inhibition of RBD attachment to host receptors (Walls et al., 2019). Passive administration of these Abs protected mice from lethal challenges with MERS-CoV, SARS-CoV, and closely related viruses, indicating that they represent a promising therapeutic strategy against coronaviruses (Corti et al., 2015; Menachery et al., 2015; Menachery et al., 2016; Rockx et al., 2008). More recently, we identified a human monoclonal Ab that neutralizes SARS-CoV-2 and SARS-CoV through recognition of the RBD from the memory B cells of a SARS survivor obtained 10 years after recovery (Pinto et al., 2020). These findings showed that the RBD is a prime target of neutralizing Abs upon natural CoV infection, in agreement with reports of the isolation of RBD-targeted neutralizing Abs from COVID-19 convalescent patients (Barnes et al., 2020; Brouwer et al., 2020; Robbiani et al., 2020; Seydoux et al., 2020; Wang et al., 2020a; Wu et al., 2020) and the demonstration that they provide *in vivo* protection against SARS-CoV-2 challenge in small animals and non-human primates (Alsoussi et al., 2020; Wu et al., 2020; Zost et al., 2020). Collectively, these observations, along with a correlation between the presence of RBD-directed Abs and neutralization potency of COVID-19 patient plasma (Robbiani et al., 2020), motivate the use of the SARS-CoV-2 RBD as a vaccine immunogen.

Vaccine development efforts responding to the COVID-19 pandemic have made extensive use of platform technologies for antigen design, antigen display, and vaccine delivery (Kanekiyo et al., 2019a). For example, existing nucleic acid and vectored vaccine platforms enabled rapid entry into the clinic with vaccines encoding SARS-CoV-2 S antigens (Folegatti et al., 2020; Jackson et al., 2020; Mulligan et al., 2020; Sahin et al., 2020; Yu et al., 2020; Zhu et al., 2020a). However, the safety, efficacy, and scalability of these vaccine modalities are not fully understood, as there are currently no DNA or mRNA vaccines licensed for human use, and the first viral vector vaccine was approved only within the last few months (Anywaine et al., 2019). In contrast, self-assembling or particulate protein immunogens are a clinically validated vaccine modality with a proven track record of safety and efficacy in humans (Lopez-Sagaseta et al., 2016). For example, virus-like particle (VLP) vaccines for human papillomavirus (HPV) and hepatitis B virus (HBV) are among the most effective subunit vaccines known, with data suggesting that the HPV vaccines in particular provide potent, durable immunity even after a single vaccination (Kreimer et al., 2018). Recently, self-assembling protein platforms for heterologous antigen display have matured significantly, and several protein nanoparticle vaccines displaying viral glycoprotein antigens are currently being evaluated in clinical trials (NCT03547245, NCT03186781, NCT03814720). A new development in this area has been the emergence of computationally designed protein nanoparticles as a robust and versatile platform for multivalent antigen presentation (Bale et al., 2016; Hsia et al., 2016; King et al., 2012; Ueda et al., 2020). In preclinical studies, vaccine candidates based on designed protein nanoparticles have significantly improved the potency or breadth of antibody responses against numerous antigens, including prefusion RSV F (Marcandalli et al., 2019), HIV envelope (Brouwer et al., 2019), influenza hemagglutinin (Boyoglu-Barnum et al., 2020), and *P. falciparum* CyRPA (Bruun et al., 2018), relative to either soluble antigen or commercial vaccine comparators.

Here we report designed protein nanoparticle vaccines multivalently displaying the SARS-CoV-2 RBD that elicit potent and protective Ab responses in mice, with neutralizing titers an order of magnitude higher at a ~five-fold lower dose than soluble prefusion-stabilized S while also exhibiting a significantly lower binding:neutralizing ratio than convalescent human sera. We further show that nanoparticle vaccine-elicited Abs recognize multiple distinct RBD epitopes targeted by known neutralizing Abs, suggesting that they may not be easily susceptible to escape mutations.

## RESULTS

### Design, *In Vitro* Assembly, and Characterization of SARS-CoV-2 RBD Nanoparticle Immunogens

To design vaccine candidates that induce potent neutralizing Ab responses, we focused on the RBD of the SARS-CoV-2 S glycoprotein (Figure 1A–B). To overcome the limited immunogenicity of this small, monomeric antigen, we multivalently displayed the RBD on the exterior surface of the two-component protein nanoparticle I53-50 (Bale et al., 2016). I53-50 is a computationally designed, 28 nm, 120-subunit complex with icosahedral symmetry constructed from trimeric (I53-50A) and pentameric (I53-50B) components (all amino acid sequences provided in Supplementary Item 1). The nanoparticle can be assembled *in vitro* by simply mixing independently expressed and purified I53-50A and I53-50B, a feature that has facilitated its use as a platform for multivalent antigen presentation (Brouwer et al., 2019; Marcandalli et al., 2019). The RBD (residues 328–531) was genetically fused to I53-50A using linkers comprising 8, 12, or 16 glycine and serine residues (hereafter referred to as RBD-8GS-, RBD-12GS-, or RBD-16GS-I53-50A) to enable flexible presentation of the antigen extending from the nanoparticle surface (Figure 1C). All RBD-I53-50A constructs were recombinantly expressed using mammalian (Expi293F) cells to ensure proper folding and glycosylation of the viral antigen. Initial yields of purified RBD-I53-50A proteins (~30 mg purified protein per liter Expi293F cells) were ~20-fold higher than for the prefusion-stabilized S-2P trimer (Kirchdoerfer et al., 2018; Pallesen et al., 2017; Walls et al., 2020; Wrapp et al., 2020) (~1.5 mg/L), and increased to ~60 mg/L following promoter optimization. The RBD-I53-50A proteins were mixed with pentameric I53-50B purified from *E. coli* in a ~1:1 molar ratio (subunit:subunit) to initiate nanoparticle assembly (Figure 1D).

**Figure 1.**
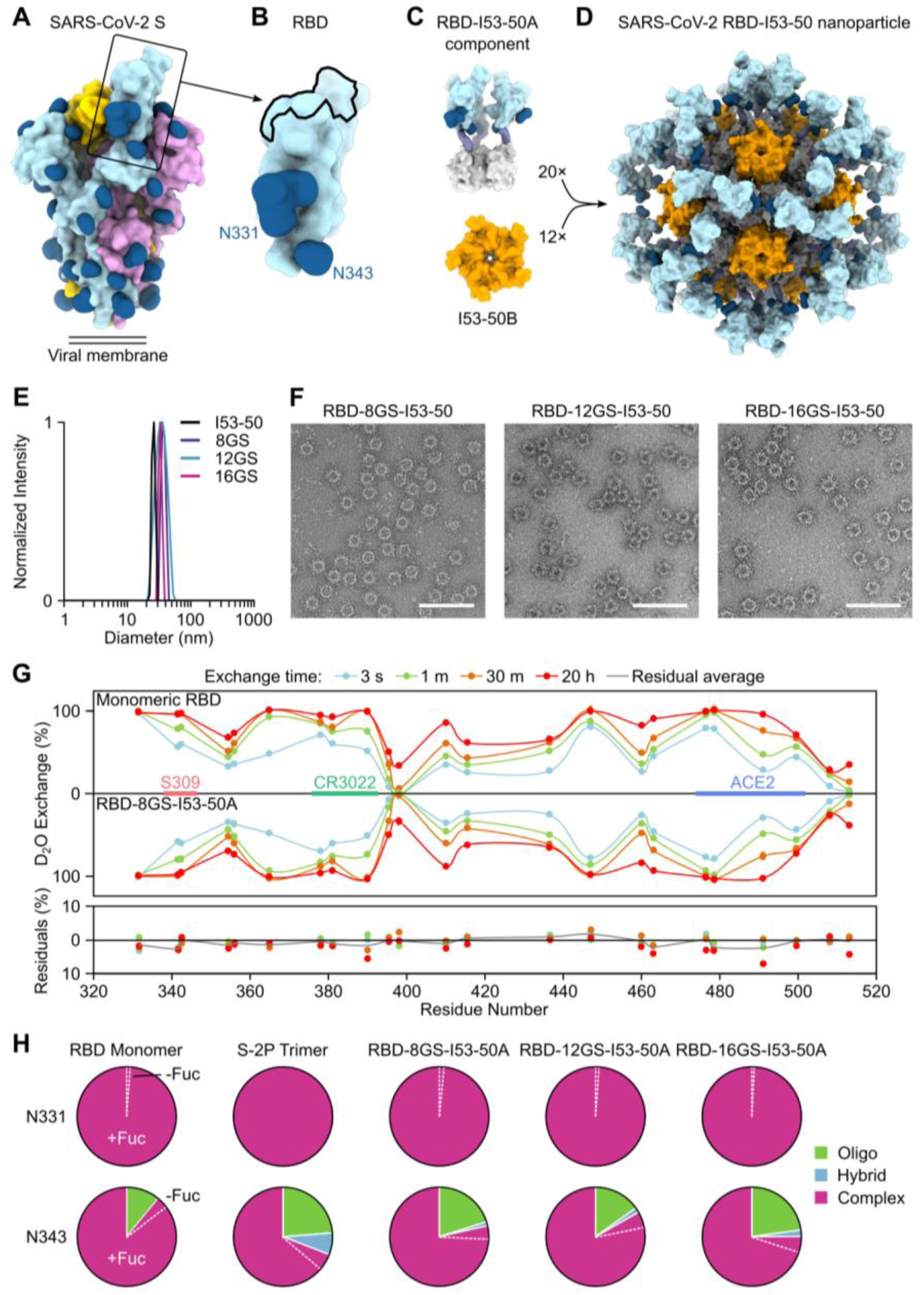
Design, *In Vitro* Assembly, and Characterization of SARS-CoV-2 RBD Nanoparticle Immunogens. (A) Molecular surface representation of the SARS-CoV-2 S-2P trimer in the prefusion conformation (PDB 6VYB). Each protomer is colored distinctly, and N-linked glycans are rendered dark blue (the glycan at position N343 was modeled based on PDB 6WPS and the receptor-binding motif (RBM) was modeled from PDB 6M0J). The single open RBD is boxed. (B) Molecular surface representation of the SARS-CoV-2 S RBD, including the N-linked glycans at positions 331 and 343. The ACE2 receptor-binding site or RBM is indicated with a black outline. (C) Structural models of the trimeric RBD-I53-50A (RBD in light blue and I53-50A in light gray) and pentameric I53-50B (orange) components. Upon mixing *in vitro*, 20 trimeric and 12 pentameric components assemble to form nanoparticle immunogens with icosahedral symmetry. Each nanoparticle displays 60 copies of the RBD. (D) Structural model of the RBD-12GS-I53-50 nanoparticle immunogen. Although a single orientation of the displayed RBD antigen and 12-residue linker are shown for simplicity, these regions are expected to be flexible relative to the I53-50 nanoparticle scaffold. (E) Dynamic light scattering (DLS) of the RBD-8GS-, RBD-12GS-, and RBD-16GS-I53-50 nanoparticles compared to unmodified I53-50 nanoparticles. (F) Representative electron micrographs of negatively stained RBD-8GS-, RBD-12GS-, and RBD-16GS-I53-50 nanoparticles. The samples were imaged after one freeze/thaw cycle. Scale bars, 100 nm. (G) Hydrogen/Deuterium-exchange mass spectrometry of monomeric RBD versus trimeric RBD-8GS-I53-50A component, represented here as a butterfly plot, confirms preservation of the RBD conformation, including at epitopes recognized by known neutralizing Abs. In the plot, each point along the horizontal sequence axis represents a peptide where deuterium uptake was monitored from 3 seconds to 20 hours. Error bars shown on the butterfly plot indicate standard deviations from two experimental replicates. The difference plot below demonstrates that monomeric RBD and RBD-8GS-I53-50A are virtually identical in local structural ordering across the RBD. (H) Pie charts summarizing the glycan populations present at the N-linked glycosylation sites N331 and N343 in five protein samples: monomeric RBD, S-2P trimer, and RBD-8GS-, RBD-12GS-, and RBD-16GS-I53-50A trimeric components. The majority of the complex glycans at both sites were fucosylated; minor populations of afucosylated glycans are set off by dashed white lines. Oligo, oligomannose.

Size-exclusion chromatography (SEC) of the SARS-CoV-2 RBD-I53-50 nanoparticles revealed predominant peaks corresponding to the target icosahedral assemblies and smaller peaks comprising residual unassembled RBD-I53-50A components (Figures S1A and S1B). Dynamic light scattering (DLS) and negative stain electron microscopy (nsEM) confirmed the homogeneity and monodispersity of the various RBD-I53-50 nanoparticles, both before and after freeze/thaw (Figures 1E, 1F, and S1C). The average hydrodynamic diameter and percent polydispersity measured by DLS for RBD-8GS-, RBD-12GS-, and RBD-16GS-I53-50 before freeze/thaw were 38.5 (27%), 37 (21%), and 41 (27%) nm, respectively, compared to 30 (22%) nm for unmodified I53-50 nanoparticles. Hydrogen/Deuterium-exchange mass spectrometry confirmed that display of the RBD on the trimeric RBD-8GS-I53-50A component preserved the conformation of the antigen and structural order of several distinct antibody epitopes (Figures 1G and S1D). Finally, we used glycoproteomics to show that all three RBD-I53-50A components were N-glycosylated at positions N331 and N343 similarly to the SARS-CoV-2 S-2P ectodomain trimer (Watanabe et al., 2020), again suggesting that the displayed antigen retained its native antigenic properties (Figure 1H and S1E).

### Antigenic Characterization of SARS-CoV-2 RBD-I53-50 Nanoparticle Components and Immunogens

We used recombinant human ACE2 ectodomain and two S-specific mAbs (CR3022 and S309) to characterize the antigenicity of the RBD when fused to I53-50A as well as the accessibility of multiple RBD epitopes in the context of the assembled nanoparticle immunogens. CR3022 and S309 were both isolated from individuals infected with SARS-CoV and cross-react with the SARS-CoV-2 RBD. CR3022 is a weakly neutralizing Ab that binds to a conserved, cryptic epitope in the RBD that becomes accessible upon RBD opening but is distinct from the receptor binding motif (RBM), the surface of the RBD that interacts with ACE2 (Huo et al., 2020; ter Meulen et al., 2006; Yuan et al., 2020). S309 neutralizes both SARS CoV and SARS-CoV-2 by binding to a glycan-containing epitope that is conserved amongst sarbecoviruses and accessible in both the open and closed prefusion S conformational states (Pinto et al., 2020).

We used bio-layer interferometry (BLI) to confirm the binding affinities of the monomeric human ACE2 (hACE2) ectodomain and the CR3022 Fab for the monomeric RBD. Equilibrium dissociation constants (K_D_) of these reagents for immobilized RBD-I53-50A fusion proteins closely matched those obtained for the monomeric RBD (Table 1 and Figure S2). These data further confirm that the RBD-I53-50A fusion proteins display the RBD in its native conformation.

**Table 1.** Antigenic Characterization of SARS-CoV-2 RBD-I53-50A Components. Each experiment was performed at least twice, and the values and fitting errors presented are derived from a representative experiment. The corresponding binding curves and fits are presented in Fig. S2.

| Antigen | Binder | $k_{on}$ ($M^{-1} s^{-1}$) | $k_{off}$ ($s^{-1}$) | $K_D$ (nM) |
| --- | --- | --- | --- | --- |
| SARS-CoV-2 RBD | hACE2 | $7 \times 10^4 \pm 5 \times 10^2$ | $5 \times 10^{-3} \pm 1 \times 10^{-5}$ | $69 \pm 0.5$ |
| | CR3022 Fab | $2 \times 10^5 \pm 2 \times 10^3$ | $9 \times 10^{-3} \pm 3 \times 10^{-5}$ | $45 \pm 0.5$ |
| RBD-8GS-I53-50A | hACE2 | $6 \times 10^4 \pm 4 \times 10^2$ | $4 \times 10^{-3} \pm 1 \times 10^{-5}$ | $70 \pm 0.5$ |
| | CR3022 Fab | $2 \times 10^5 \pm 1 \times 10^3$ | $1 \times 10^{-2} \pm 3 \times 10^{-5}$ | $57 \pm 0.4$ |
| RBD-12GS-I53-50A | hACE2 | $6 \times 10^4 \pm 4 \times 10^2$ | $5 \times 10^{-3} \pm 1 \times 10^{-5}$ | $78 \pm 0.5$ |
| | CR3022 Fab | $2 \times 10^5 \pm 2 \times 10^3$ | $9 \times 10^{-3} \pm 2 \times 10^{-5}$ | $42 \pm 0.4$ |
| RBD-16GS-I53-50A | hACE2 | $6 \times 10^4 \pm 4 \times 10^2$ | $4 \times 10^{-2} \pm 1 \times 10^{-5}$ | $66 \pm 0.4$ |
| | CR3022 Fab | $2 \times 10^5 \pm 1 \times 10^3$ | $1 \times 10^{-2} \pm 2 \times 10^{-5}$ | $56 \pm 0.4$ |

We previously observed that the magnitude and quality of nanoparticle immunogen-elicited Ab responses can be modulated by the accessibility of specific epitopes in the context of a dense, multivalent antigen array, most likely through steric crowding (Brouwer et al., 2019). To evaluate this possibility, we measured the binding of the nanoparticle immunogens to immobilized dimeric macaque ACE2 (mACE2-Fc) and the CR3022 and S309 mAbs, the latter of which roughly mimics the B cell receptor (BCR)-antigen interaction that is central to B cell activation. This approach does not allow the calculation of K_D_ values due to the multivalent nature of the interactions, but does enable qualitative comparisons of epitope accessibility in different nanoparticle constructs. We compared the full-valency nanoparticles displaying 60 RBDs to a less dense antigen array by leveraging the versatility of *in vitro* assembly to prepare nanoparticle immunogens displaying the RBD antigen at 50% valency (~30 RBDs per nanoparticle) (Figure S3). This was achieved by adding pentameric I53-50B to an equimolar mixture of RBD-I53-50A and unmodified I53-50A lacking fused antigen. We found that all of the RBD nanoparticles bound well to the immobilized mACE2-Fc, CR3022, and S309 (Figure 2A). Although there were no consistent trends among the 50% and 100% valency RBD-8GS-and RBD-12GS-I53-50 nanoparticles, the 100% valency RBD-16GS-I53-50 nanoparticles resulted in the highest binding signals against all three binders (Figure 2B). It is possible that the longer linker in the RBD-16GS-I53-50 nanoparticle enables better access to the epitopes targeted by ACE2, CR3022, and S309, although our data cannot rule out other possible explanations. We conclude that multiple distinct epitopes targeted by neutralizing antibodies are exposed and accessible for binding in the context of the RBD antigen array presented on the nanoparticle exterior.

**Figure 2.**
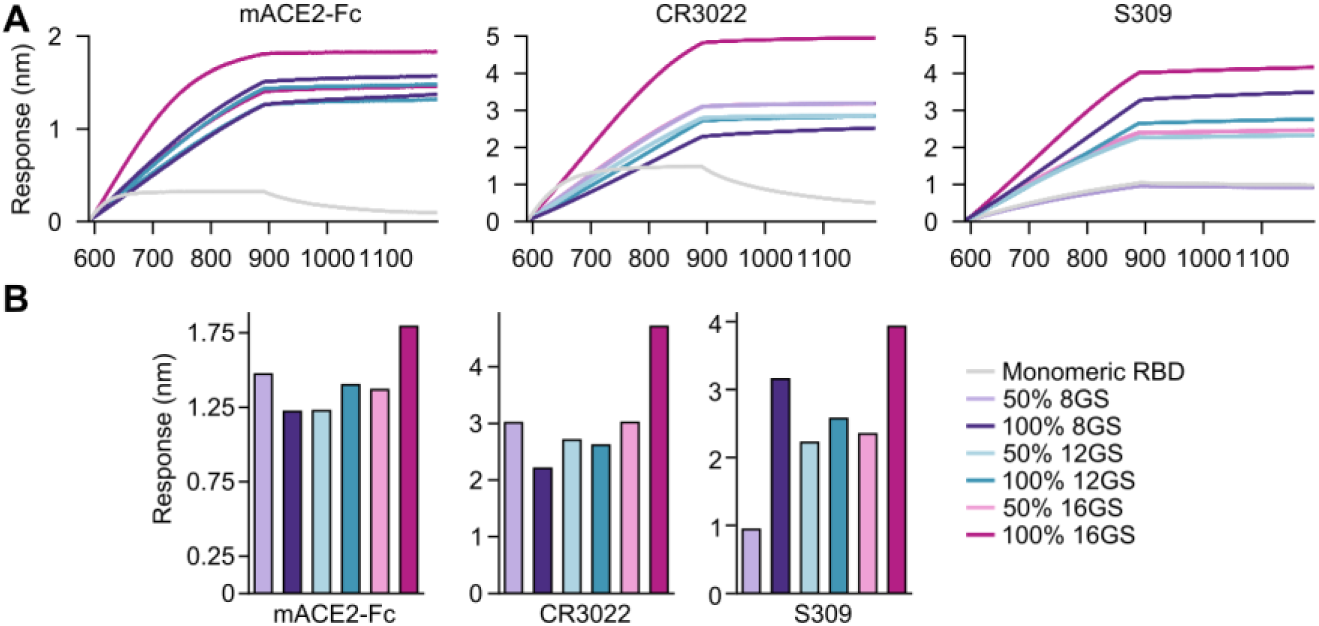
Antigenic Characterization of SARS-CoV-2 RBD-I53-50 Nanoparticle Immunogens. (A) Bio-layer interferometry of immobilized mACE2-Fc, CR3022 mAb, and S309 mAb binding to RBD-8GS-, RBD-12GS-, and RBD-16GS-I53-50 nanoparticles displaying the RBD antigen at 50% or 100% valency. The monomeric SARS-CoV-2 RBD was included in each experiment as a reference. (B) The binding signal at 880 s, near the end of the association phase, is plotted for each experiment in panel (A) to enable comparison of the binding signal obtained from each nanoparticle.

### Physical and Antigenic Stability of RBD Nanoparticle Immunogens and S-2P Trimer

Although subunit vaccines based on recombinant protein antigens have several intrinsic advantages over other vaccine modalities, they must meet stringent requirements related to stability during manufacture, storage, and distribution (Kumru et al., 2014). We first used chemical denaturation in guanidine hydrochloride (GdnHCl) to compare the stability of the RBD-I53-50A fusion proteins and RBD-12GS-I53-50 nanoparticle immunogen to recombinant monomeric RBD and the S-2P ectodomain trimer (Figure 3A). Fluorescence emission spectra from samples incubated in 0–6.5 M GdnHCl revealed that all three RBD-I53-50A fusion proteins and the RBD-12GS-I53-50 nanoparticle undergo a transition between 4 and 5 M GdnHCl that indicates at least partial unfolding, whereas the S-2P trimer showed a transition at lower [GdnHCl], between 2 and 4 M. The monomeric RBD exhibited a less cooperative unfolding transition over 0–5 M GdnHCl. We then used a suite of analytical assays to monitor physical and antigenic stability over four weeks post-purification at three temperatures: <−70°C, 2-8°C, and 22-27°C (Figure 3B–E). Consistent with previous reports, the monomeric RBD proved quite stable, yielding little change in appearance by SDS-PAGE (Figure S4A), mACE2-Fc and CR3022 binding (Figure S4B), or the ratio of UV/vis absorption at 320/280 nm, a measure of particulate scattering (Figure S4C). As reported recently (Edwards et al., 2020; Hsieh et al., 2020), the S-2P trimer was unstable at 2-8°C, exhibiting clear signs of unfolding by nsEM even at early time points (Figure S3D and Supplementary Item 2). It maintained its structure considerably better at 22-27°C until the latest time point (28 days), when unfolding was apparent by nsEM and UV/vis indicated some aggregation (Figure S4C). All three RBD-I53-50A components were highly stable, exhibiting no substantial change in any readout at any time point (Supplementary Item 2). Finally, the RBD-12GS-I53-50 nanoparticle was also quite stable over the four-week study, showing changes only in UV/vis absorbance, where a peak near 320 nm appeared after 7 days at 22-27°C (Supplementary Item 2). Electron micrographs and DLS of the RBD-12GS-I53-50 nanoparticle samples consistently showed monodisperse, well-formed nanoparticles at all temperatures over the four-week period (Figures S4D, S4E, and Supplementary Item 2). Collectively, these data show that the RBD-I53-50A components and the RBD-12GS-I53-50 nanoparticle have high physical and antigenic stability, superior to the S-2P ectodomain trimer.

**Figure 3.**
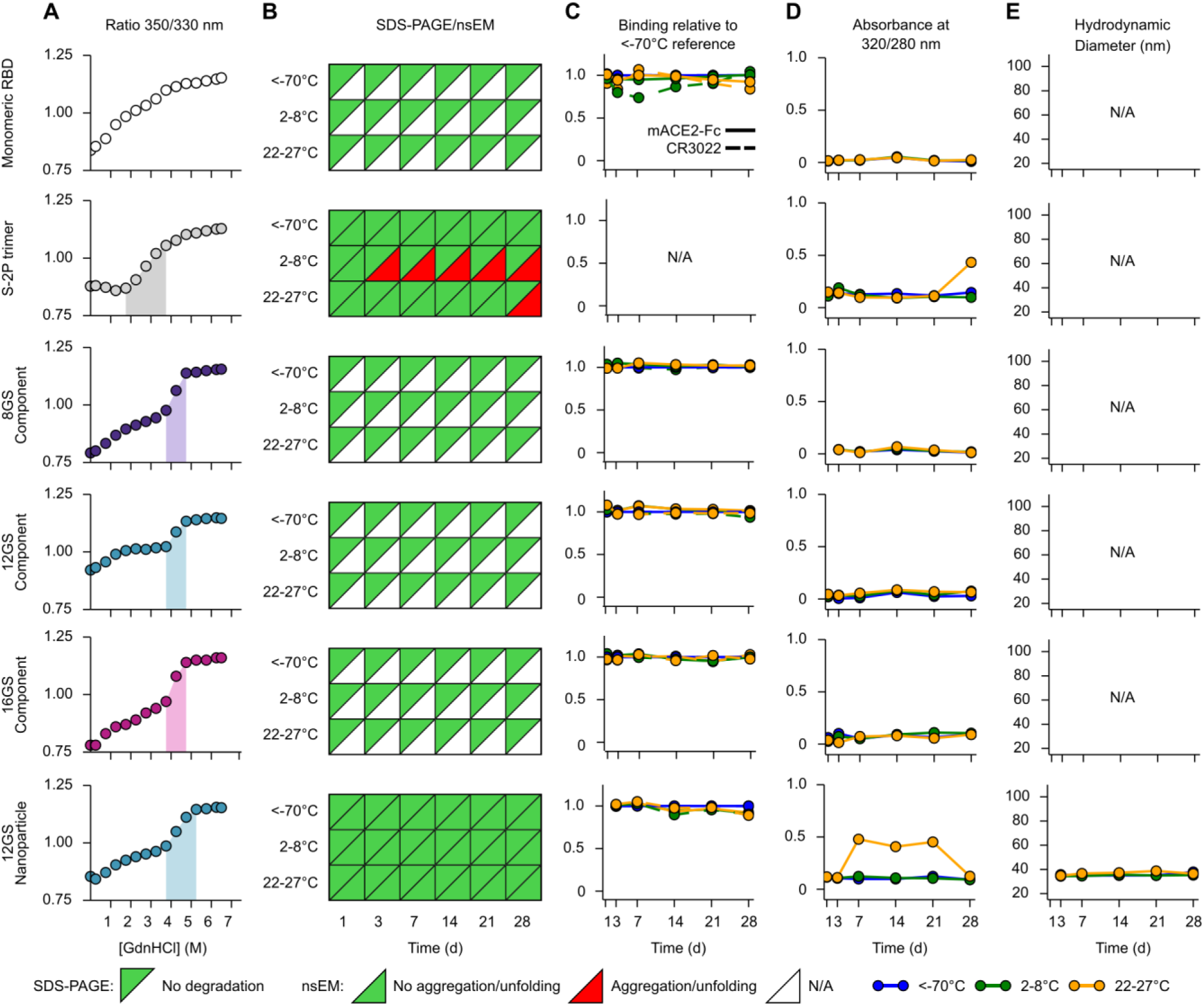
Physical and Antigenic Stability of RBD Nanoparticle Immunogens and S-2P Trimer. (A) Chemical denaturation by guanidine hydrochloride. The ratio of intrinsic tryptophan fluorescence emission at 350/320 nm was used to monitor protein tertiary structure. Major transitions are indicated by shaded regions. Representative data from one of three independent experiments are shown. (B) Summary of SDS-PAGE and nsEM stability data over four weeks. SDS-PAGE showed no detectable degradation in any sample. nsEM revealed substantial unfolding of the S-2P trimer at 2-8°C after three days incubation, and at 22-27°C after four weeks. N/A, not assessed. (C) Summary of antigenicity data over four weeks. The antigens were analyzed for mACE2-Fc (solid lines) and CR3022 mAb (dashed lines) binding by bio-layer interferometry after storage at the various temperatures. The plotted value represents the amplitude of the signal near the end of the association phase normalized to the corresponding <−70°C sample at each time point. (D) Summary of UV/vis stability data over four weeks. The ratio of absorbance at 320/280 nm is plotted as a measure of particulate scattering. Only the S-2P trimer and the RBD-12GS-I53-50 nanoparticle showed any increase in scattering, and only at ambient temperature. (E) DLS of the RBD-12GS-I53-50 nanoparticle indicated a monodisperse species with no detectable aggregate at all temperatures and time points. The data in panels B–E is from a four-week real-time stability study that was performed once.

### RBD-I53-50 Nanoparticle Immunogens Elicit Potent Neutralizing Antibody Responses in BALB/c and Human Immune Repertoire Mice

We compared the immunogenicity of the three RBD-I53-50 nanoparticles, each displaying the RBD at either 50% or 100% valency, to the S-2P ectodomain trimer and the monomeric RBD in BALB/c mice. Groups of ten mice were immunized intramuscularly at weeks 0 and 3 with AddaVax-adjuvanted formulations containing either 0.9 or 5 µg of SARS-CoV-2 antigen in either soluble or particulate form. Three weeks post-prime, all RBD nanoparticles elicited robust S-specific Ab responses with geometric mean reciprocal half-maximal effective concentrations ranging between 8×10^2^ and 1×10^4^ (Figure 4A, Supplementary Items 3 and 4). In contrast, the monomeric RBD and the low dose of S-2P trimer did not induce detectable levels of S-specific Abs, while the high dose of S-2P trimer elicited weak responses. Following a second immunization, we observed an enhancement of S-specific Ab titers for all RBD nanoparticle groups, with geometric mean titers (GMT) ranging from 1×10^5^ to 2×10^6^ (Figure 4B and Supplementary Items 3 and 4). These levels of S-specific Abs matched or exceeded most samples from a panel of 29 COVID-19 human convalescent sera (HCS) from Washington state and the benchmark 20/130 COVID-19 plasma from NIBSC (Figure 4A–B, Supplementary Item 5). Immunization with two 5 µg doses of S-2P trimer induced S-specific Ab responses ~1-2 orders of magnitude weaker than the RBD nanoparticles, and the monomeric RBD did not elicit detectable antigen-specific Abs after two immunizations. These data indicate that multivalent display of the RBD on a self-assembling nanoparticle scaffold dramatically improves its immunogenicity.

**Figure 4.**
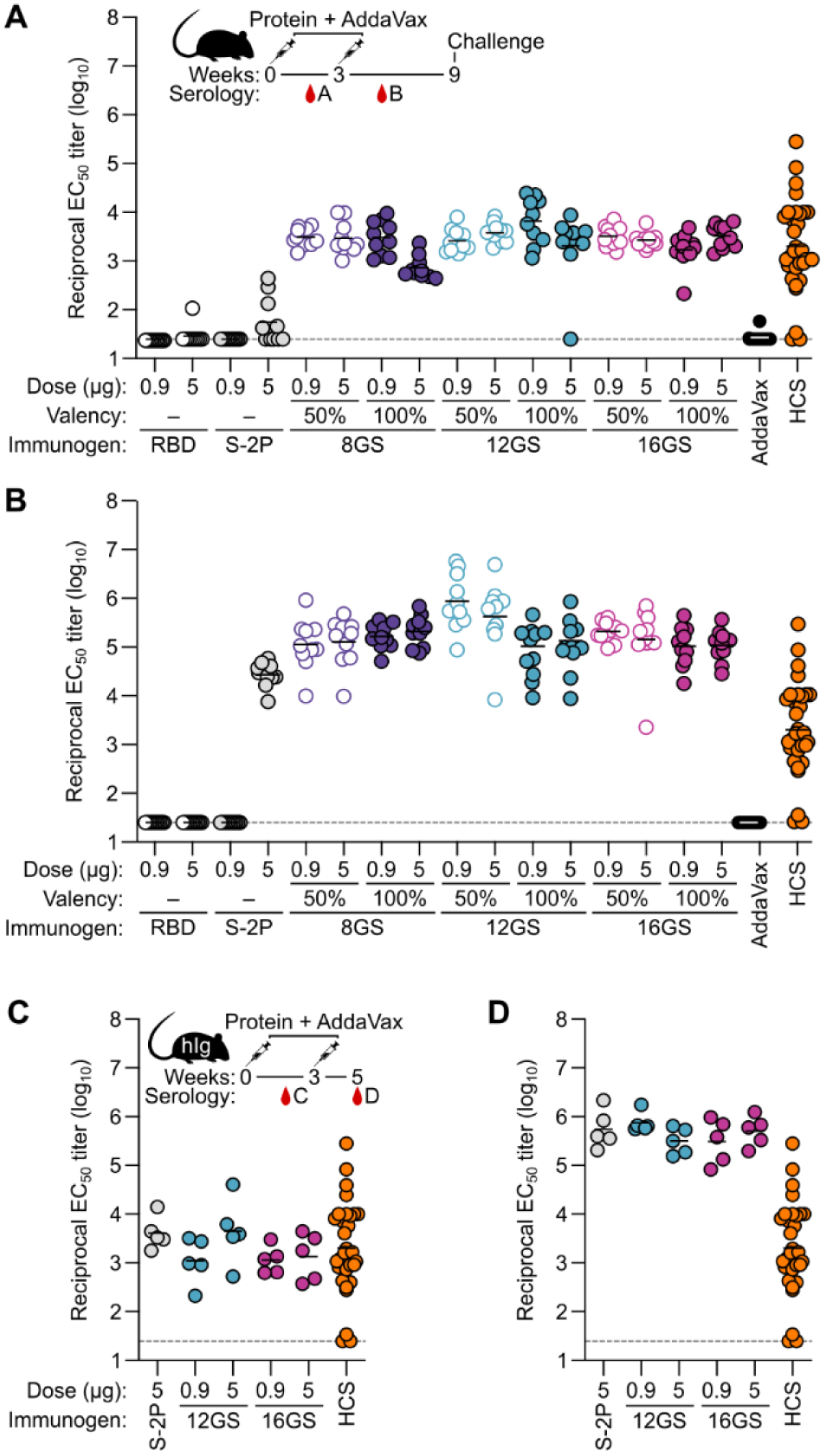
RBD-I53-50 Nanoparticle Immunogens Elicit Potent Antibody Responses in BALB/c and Human Immune Repertoire Mice. (A–B) Post-prime (week 2) (A) and post-boost (week 5) (B) anti-S binding titers in BALB/c mice, measured by ELISA. Each symbol represents an individual animal, and the geometric mean from each group is indicated by a horizontal line. The dotted line represents the lower limit of detection of the assay. 8GS, RBD-8GS-I53-50; 12GS, RBD-12GS-I53-50; 16GS, RBD-16GS-I53-50; HCS, human convalescent sera. The inset depicts the study timeline. The immunization experiment was repeated twice and representative data are shown. (C–D) Post-prime (week 2) (C) and post-boost (week 5) (D) anti-S binding titers in Kymab Darwin mice, which are transgenic for the non-rearranged human antibody variable and constant region germline repertoire, measured by ELISA and plotted as in (A). The inset depicts the study timeline. The immunization experiment was performed once. Statistical analyses are provided in Supplementary Item 4.

We prototyped potential human antibody responses to the RBD nanoparticle immunogens using the Kymab proprietary IntelliSelect™ Transgenic mouse platform (known as ‘Darwin’) that is transgenic for the non-rearranged human antibody variable and constant region germline repertoire. In contrast to previous mice with chimeric antibody loci that have been described (Lee et al., 2014), the mice in the present study differed in that they were engineered to express fully human kappa light chain Abs. Groups of five Darwin mice were immunized intramuscularly with S-2P trimer, 100% RBD-12GS-, or 100% RBD-16GS-I53-50 nanoparticles at antigen doses of 0.9 µg (nanoparticles only) or 5 µg (Figure 4C). All groups immunized with RBD nanoparticles elicited S-directed Ab responses post-prime (EC_50_ 2×10^3^–1×10^4^) that were substantially boosted by a second immunization at week 3 (EC_50_ ranging from 4×10^5^ to 8×10^5^) (Figures 4C and 4D, Supplementary Items 3 and 4). In this animal model, the S-2P trimer elicited levels of S-specific Abs comparable to the RBD nanoparticles after each immunization.

We then evaluated the neutralizing activity elicited by each immunogen using both pseudovirus and live virus neutralization assays. In BALB/c mice, all RBD nanoparticle immunogens elicited serum neutralizing Abs after a single immunization, with reciprocal half-maximal inhibition dilutions (IC_50_) ranging from 1×10^2^ to 5×10^2^ (GMT) in pseudovirus and 3×10^3^ to 7×10^3^ in live virus neutralization assays (Figure 5A and 5C, Supplementary Items 3 and 4). No significant differences in pseudovirus or live virus neutralization were observed between low or high doses of RBD-8GS-, RBD-12GS-, or RBD-16GS-I53-50 nanoparticles at 50% (pseudovirus neutralization only) or 100% valency, in agreement with the S-specific Ab data. The GMT of all three 100% valency RBD nanoparticle groups matched or exceeded that of the panel of 29 HCS tested in the pseudovirus neutralization assay (Figure 5A). Immunization with monomeric RBD or S-2P trimer did not elicit neutralizing Abs after a single immunization, in line with the observed lack of S-directed Ab responses and a previous study (Mandolesi et al., 2020) (Figures 5A and 5C). As in BALB/c mice, both high and low doses of the RBD-I53-50 nanoparticles in Darwin mice elicited pseudovirus neutralizing Ab titers (ic_50_ 8×10^1^ to 2.5×10^2^) comparable to HCS (IC_50_ 1×10^2^) after a single immunization, whereas 5 μg of the S-2P trimer did not elicit detectable levels of neutralizing Abs (Figure 5E and Supplementary Item 3) despite eliciting similar levels of total S-specific Abs.

**Figure 5.**
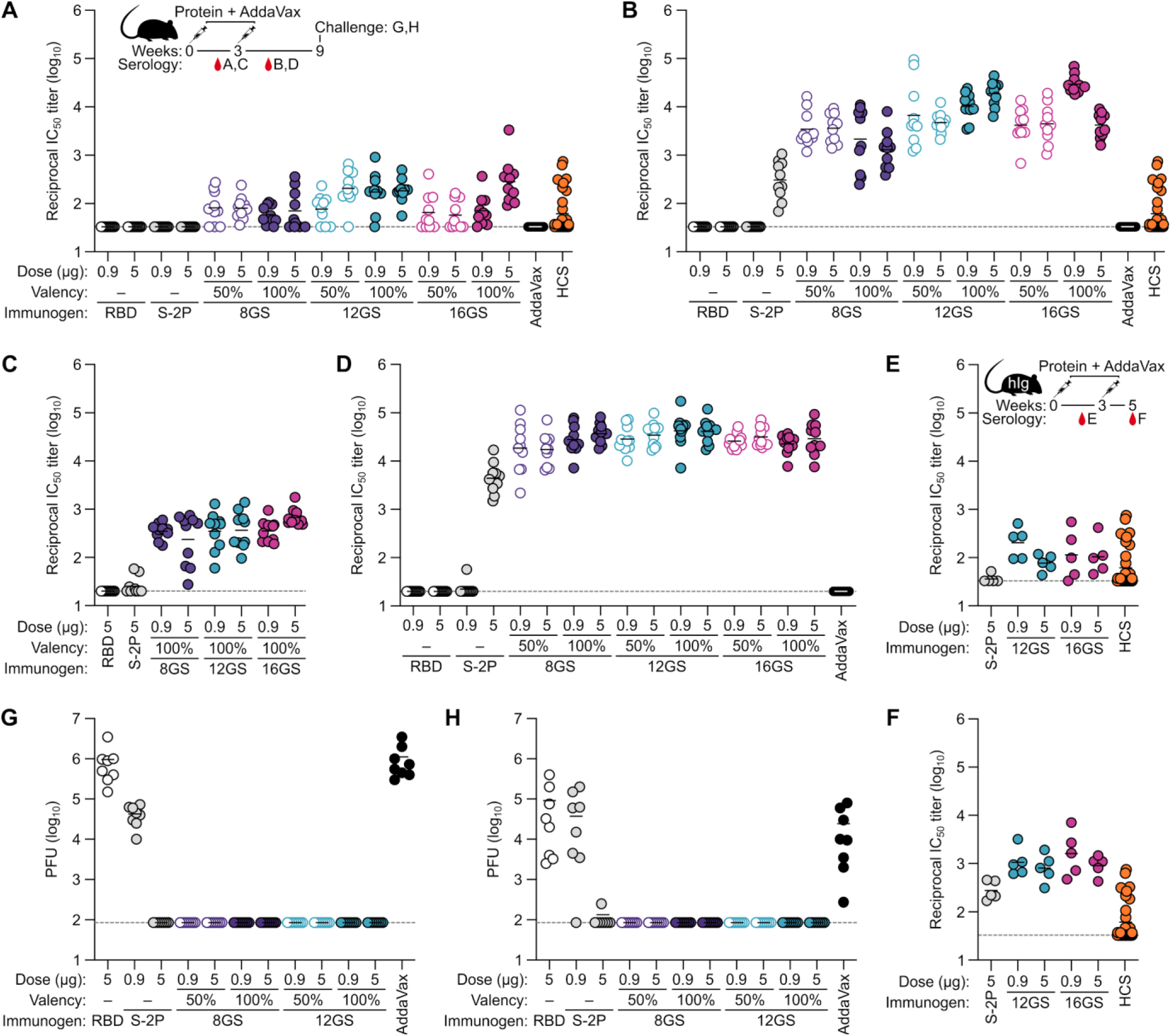
RBD-I53-50 Nanoparticle Immunogens Elicit Potent and Protective Neutralizing Antibody Responses. (A–B) Serum pseudovirus neutralizing titers post-prime (A) or post-boost (B) from mice immunized with monomeric RBD, S-2P trimer, or RBD-I53-50 nanoparticles. Each circle represents the reciprocal IC_50_ of an individual animal. The geometric mean from each group is indicated by a horizontal line. Limit of detection shown as a gray dotted line. The animal experiment was performed twice, and representative data from duplicate measurements are shown. (C–D) Serum live virus neutralizing titers post-prime (C) or post-boost (D) from mice immunized as described in (A). (E–F) Serum pseudovirus neutralizing titers from Kymab Darwin mice post-prime (E) and post-boost (F), immunized as described in (A). The animal experiment was performed once, and the neutralization assays were performed at least in duplicate. (G–H) Seven weeks post-boost, eight BALB/c mice per group were challenged with SARS-CoV-2 MA. Two days post-challenge, viral titers in lung tissue (G) and nasal turbinates (H) were assessed. Limit of detection depicted as a gray dotted line. Statistical analyses are provided in Supplementary Item 4.

In both mouse models, a second immunization with the RBD-I53-50 nanoparticles led to a large increase in neutralizing Ab titers. In BALB/c mice, pseudovirus neutralization GMT reached 2×10^3^ to 3×10^4^, exceeding that of the HCS by 1–2 orders of magnitude, and live virus neutralization titers reached 2×10^4^ to 3×10^4^ (Figures 5B and 5D). A second immunization with 5 μg of the S-2P trimer also strongly boosted neutralizing activity, although pseudovirus and live virus neutralization (GMTs of 3×10^2^ and 6×10^3^, respectively) were still lower than in sera from animals immunized with the RBD nanoparticles. The increases between the S-2P trimer and the RBD nanoparticles ranged from 7–90-fold and 4–9-fold in the pseudovirus and live virus neutralization assays, respectively. The 0.9 μg dose of the S-2P trimer and both doses of the monomeric RBD failed to elicit detectable neutralization after two immunizations. Similar increases in pseudovirus neutralization were observed after the second immunization in the Darwin mice, although the titers were lower overall than in BALB/c mice (Figure 5F and Supplementary Item 3).

Several conclusions can be drawn from these data. First, the RBD nanoparticles elicit potent neutralizing Ab responses in two mouse models that exceed those elicited by the prefusion-stabilized S-2P trimer and, after two doses, by infection in humans. Second, linker length and antigen valency did not substantially impact the overall immunogenicity of the RBD nanoparticles, although there is a trend suggesting that RBD-16GS-I53-50 may be more immunogenic than the nanoparticles with shorter linkers. These observations are consistent with the antigenicity and accessibility data presented in Table 1 and Figure 2 showing that multiple epitopes are intact and accessible in all RBD nanoparticle immunogens. Finally, the elicitation of comparable neutralizing Ab titers by both the 0.9 and 5 µg doses of each nanoparticle immunogen suggests that RBD presentation on the I53-50 nanoparticle enables dose sparing, which is a key consideration for vaccine manufacturing and distribution.

Eight mice immunized with AddaVax only, monomeric RBD, S-2P trimer, or RBD-8GS- or RBD-12GS-I53-50 nanoparticles were challenged seven weeks post-boost with a mouse-adapted SARS-CoV-2 virus (SARS-CoV-2 MA) to determine whether these immunogens confer protection from viral replication (Dinnon et al., 2020). The RBD-8GS- and RBD-12GS-I53-50 nanoparticles provided complete protection from detectable SARS-CoV-2 MA replication in mouse lung and nasal turbinates (Figure 5G–H). Immunization with the monomeric RBD, 0.9 µg S-2P trimer, and adjuvant control did not protect from SARS-CoV-2 MA replication. These results mirrored our pseudovirus and live virus neutralization data showing that the RBD nanoparticles induce potent anti-SARS-CoV-2 Ab responses at either dose or valency.

### RBD Nanoparticle Vaccines Elicit Robust B Cell Responses and Antibodies Targeting Multiple Epitopes in Mice and a Nonhuman Primate

Germinal center (GC) responses are a key process in the formation of durable B cell memory, resulting in the formation of affinity-matured, class-switched memory B cells and long-lived plasma cells. We therefore evaluated the antigen-specific GC B cell responses in mice immunized with the monomeric RBD, S-2P trimer, and RBD-8GS-, RBD-12GS-, or RBD-16GS-I53-50 nanoparticles. The quantity and phenotype of RBD-specific B cells were assessed 11 days after immunization to determine levels of GC precursors and B cells (B220^+^CD3^−^CD138^−^CD38^−^GL7^+^) (Figure S5). Immunization with RBD nanoparticles resulted in an expansion of RBD-specific B cells and GC precursors and B cells (Figure 6A–C). The S-2P trimer resulted in a detectable but lower number and frequency of RBD-specific B cells and GC precursors and B cells compared to the RBD nanoparticles, whereas the monomeric RBD construct did not elicit an appreciable B cell response. Consistent with these findings, immunization with the three RBD nanoparticles and trimeric S-2P led to the emergence of CD38^+/−^GL7^+^ IgM^+^ and class-switched (swIg^+^) RBD-specific B cells, indicative of functional GC precursors and GC B cells (Figure 6D). The robust GC B cell responses and increased proportions of IgM^+^ and swIg^+^ RBD-specific B cells in the mice immunized with the RBD-nanoparticles and, to a lesser extent, S-2P constructs is consistent with an ongoing GC reaction, which in time should result in the formation of memory B cells and long-lived plasma cells.

**Figure 6.**
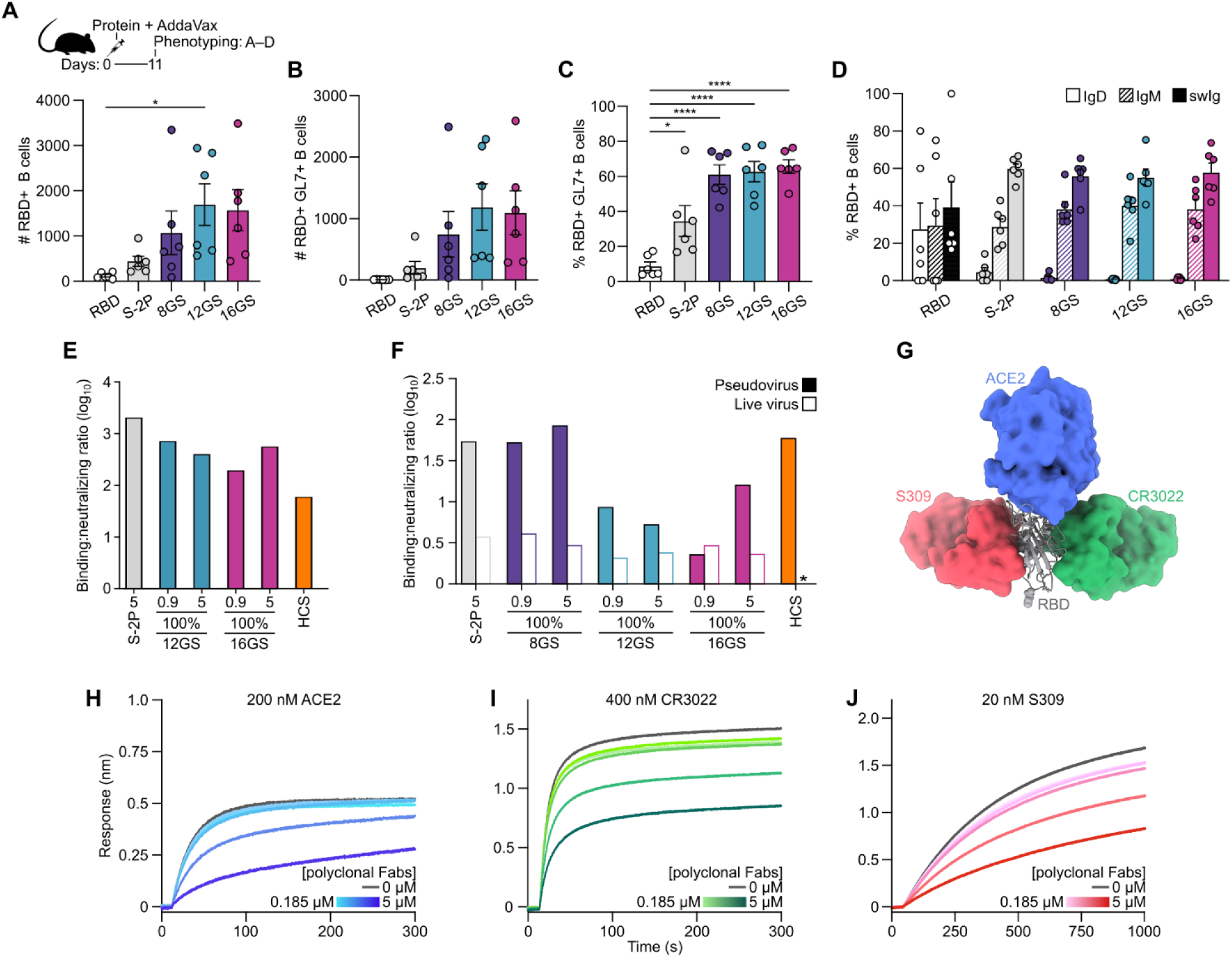
RBD Nanoparticle Vaccines Elicit Robust B Cell Responses and Antibodies Targeting Multiple Epitopes in Mice and a Nonhuman Primate. (A–B) Number of (A) RBD^+^ B cells (B220^+^CD3^−^CD138^−^) and (B) RBD^+^ GC precursors and B cells (CD38^+/−^GL7^+^) detected across each immunization group. (C–D) Frequency of (C) RBD^+^ GC precursors and B cells (CD38^+/−^GL7^+^) and (D) IgD^+^, IgM^+^, or class-switched (IgM^−^ IgD^−^; swIg^+^) RBD^+^ GC precursors and B cells. (A–D) N=6 across two experiments for each group. Statistical significance was determined by one-way ANOVA, and Tukey’s multiple comparisons tests were performed for any group with a p-value less than 0.05. Significance is indicated with stars: * p < 0.05, **** p < 0.0001. (E) Ratio post-boost (week 5) of S-2P ELISA binding titer (Figure 4D) to pseudovirus neutralization titers (Figure 5F) in Kymab Darwin mice. The ratio is the [GMT (EC_50_) of five mice]:[the GMT (IC_50_) of five mice] or the EC_50_:IC_50_ of all HCS tested. A lower value signifies a higher quality response. (F) Ratio post-boost (week 5) of S-2P ELISA binding titer (Figure 4B) to either pseudovirus (Figure 5B) or live virus (Figure 5D) neutralization titers in BALB/c mice. The ratio is the [GMT (EC_50_) of ten mice]:[the GMT (IC_50_) of ten mice] or the EC_50_:IC_50_ of all HCS tested. (G) SARS-CoV-2 RBD (gray ribbon) with monomeric ACE2 (blue surface), CR3022 Fab (green surface), and S309 Fab (red surface) bound. (H–J) Determination of vaccine-elicited Ab epitope specificity by competition BLI. A dilution series of polyclonal NHP Fabs was pre-incubated with RBD on the BLI tip. The polyclonal Fab concentration was maintained with the addition of competitor to each dilution point. The 1:3 dilution series of polyclonal Fabs is represented from dark to light, with a dark gray line representing competitor loaded to apo-RBD (no competition). Competition with (H) 200 nM ACE2, (I) 400 nM CR3022, or (J) 20 nM S309.

We compared the ratio of binding to neutralizing antibodies elicited by the S-2P and the RBD-8GS-, RBD-12GS-, and RBD-16GS-I53-50 nanoparticles and HCS as a measure of the quality of the Ab responses elicited by the nanoparticle immunogens (Graham, 2020). In Kymab Darwin mice, the nanoparticle vaccines had lower (better) ratios than S-2P–immunized mice, but higher than HCS (Figure 6E). In BALB/c mice, the ratio of binding to pseudovirus neutralizing titers elicited by RBD-12GS- and RBD-16GS-I53-50 was clearly decreased compared to S-2P and HCS (Figure 6F). This pattern was consistent when ratios were calculated using live virus neutralizing titers, although the magnitude of the differences between groups were smaller due to the high values obtained in the live virus neutralization assay. These results suggest the Ab responses elicited by the RBD-12GS- and RBD-16GS-I53-50 nanoparticle immunogens are of higher quality than that obtained from immunization with the S-2P trimer or acquired during natural infection, perhaps because it is focused on epitopes in the RBD that are the target of most neutralizing Abs.

We set out to identify the epitopes recognized by Abs elicited upon immunization with the nanoparticle immunogens in a nonhuman primate model that more closely resembles humans in their immune response to vaccination. We immunized a pigtail macaque with 250 µg of RBD-12GS-I53-50 (88 µg of RBD antigen) at weeks 0 and 4 and found that serum collected at week 8 had high levels of S-specific Abs (EC_50_ ~1×10^6^). Polyclonal Fabs were generated and purified for use in competition BLI with hACE2, CR3022, and S309, which recognize three distinct sites targeted by neutralizing Abs on the SARS-CoV-2 RBD (Figure 6G) (Barnes et al., 2020; Huo et al., 2020; Pinto et al., 2020; Yuan et al., 2020; Zhou et al., 2020a). The polyclonal sera inhibited binding of hACE2, CR3022 Fab, and S309 Fab at concentrations above their respective dissociation constants in a dose-dependent manner (Figure 6H–J). These data indicate that immunization with 12GS-RBD-I53-50 elicited Abs targeting several non-overlapping epitopes, which we expect to limit the potential for emergence and selection of escape mutants, especially since coronaviruses do not mutate quickly when compared to viruses such as influenza or human immunodeficiency virus (Li et al., 2020; Smith et al., 2014).

## DISCUSSION

A wide variety of SARS-CoV-2 vaccine candidates spanning diverse vaccine modalities are currently in preclinical or clinical development. Multivalent antigen presentation on self-assembling protein scaffolds is being increasingly explored in clinical vaccine development, largely due to the availability of robust platforms that enable the display of complex antigens, including oligomeric viral glycoproteins (Irvine and Read, 2020; Kanekiyo et al., 2019a; Lopez-Sagaseta et al., 2016). Such nanoparticle vaccines often significantly enhance neutralizing Ab responses compared to traditional subunit vaccines based on non-particulate antigens (Kanekiyo et al., 2015; Kanekiyo et al., 2013; Marcandalli et al., 2019). While preclinical and early clinical development of subunit vaccines is slower than nucleic acid and vector-based vaccines, multiple intrinsic advantages of nanoparticle vaccines strongly motivate their prioritization in SARS-CoV-2 vaccine development efforts. These advantages include the potential to induce potent neutralizing Ab responses, the ability to use existing worldwide capacity for manufacturing recombinant proteins, and an established regulatory track record.

Here we showed that two-component self-assembling SARS-CoV-2 RBD nanoparticle vaccine candidates elicit potent neutralizing Ab responses targeting multiple distinct RBD epitopes. Although comparing vaccine candidates across studies from different groups is complicated by variations in serological assays, experimental design, and many other factors, the greater neutralizing Ab responses elicited by the RBD nanoparticles compared to the prefusion-stabilized ectodomain trimer are very promising. One of the few other studies that benchmarked vaccine immunogenicity against the S-2P trimer delivered as recombinant protein was a recent preclinical evaluation of mRNA-1273 in mice (Corbett et al., 2020). At the highest dose tested (1 μg), two immunizations with mRNA-1273 elicited comparable levels of S-specific Abs to two immunizations with 1 μg of S-2P trimer formulated in an adjuvant containing a TLR4 agonist. Additional reports of potent neutralizing activity obtained by immunization with prefusion-stabilized (S-2P) ectodomain (Mandolesi et al., 2020) and full-length trimers (Keech et al., 2020) formulated with powerful adjuvants provide promising data on the immunogenicity of vaccine candidates based on recombinant S proteins. Our data indicate that RBD-12GS-I53-50 and RBD-16GS-I53-50 elicit nearly ten-fold higher levels of S-specific Abs and, more importantly, roughly ten-fold higher levels of neutralizing activity compared to the S-2P ectodomain trimer. This enhancement in potency is maintained at a more than five-fold lower antigen dose by mass, suggesting that presentation on the nanoparticle also has a dose-sparing effect. Both enhanced potency and dose-sparing could be critical for addressing the need to manufacture an unprecedented number of doses of vaccine to respond to the SARS-CoV-2 pandemic.

Several recent studies have indicated that the RBD is the target of most neutralizing activity in COVID-19 HCS (Barnes et al., 2020; Brouwer et al., 2020; Pinto et al., 2020; Robbiani et al., 2020; Seydoux et al., 2020; Wang et al., 2020a; Wu et al., 2020). Although the RBD is poorly immunogenic as a monomer, our data establish that it can form the basis of a highly immunogenic vaccine when presented multivalently. This conclusion is consistent with a recent report of dimerization increasing the immunogenicity of the RBD from several coronavirus S proteins, including SARS-CoV-2 S (Dai et al., 2020), as well as a nanoparticle vaccine candidate for Epstein-Barr virus conceptually similar to our RBD nanoparticle immunogens that displayed the CR2-binding domain of gp350 (Kanekiyo et al., 2015). The exceptionally low binding:neutralizing ratio elicited upon immunization with the RBD nanoparticles suggests that presentation of the RBD on I53-50 focuses the humoral response on epitopes recognized by neutralizing Abs. This metric has been identified as a potentially important indicator of vaccine safety, as high levels of binding yet non-neutralizing or weakly neutralizing Abs may contribute to vaccine-associated enhancement of respiratory disease (Graham, 2020; Kim et al., 1969; Polack et al., 2002). Our data further show that RBD-12GS-I53-50 elicited Ab responses targeting several of the non-overlapping epitopes recognized by neutralizing Abs that have been identified in the RBD. Such polyclonal responses targeting multiple distinct epitopes might explain the magnitude of neutralization observed and should minimize the risk of selection or emergence of escape mutations (Davis et al., 2018; Lee et al., 2019). Finally, the high production yield of RBD-I53-50A components and the robust stability of the antigen-bearing RBD nanoparticles suggests that these will likely be more amenable to large-scale manufacturing than the SARS-CoV-2 S-2P trimer, which expresses poorly and is unstable (Edwards et al., 2020; Hsieh et al., 2020; McCallum et al., 2020).

The emergence of three highly pathogenic zoonotic coronaviruses in the past two decades showcases that vaccines capable of providing broad protection against coronaviruses are urgently needed for future pandemic preparedness. As viruses similar to SARS-CoV (Menachery et al., 2015), MERS-CoV (Anthony et al., 2017; Woo et al., 2006) and SARS-CoV-2 (Zhou et al., 2020b; Zhou et al., 2020c) have been found in animal reservoirs, the potential for the emergence of similar viruses in the future poses a significant threat to global public health. The RBD nanoparticle vaccines described here are not expected to provide protection against distantly related coronaviruses (e.g., MERS-CoV) due to substantial sequence variation among the RBDs of coronavirus S glycoproteins. However, the potent neutralizing Ab responses elicited by the nanoparticle immunogens combined with recent work demonstrating that co-displaying multiple antigens on the same nanoparticle can improve the breadth of vaccine-elicited immune responses (Boyoglu-Barnum et al., 2020; Kanekiyo et al., 2019b) suggests a potential route to broader coronavirus vaccines. Alternatively, optimizing the expression, stability, and multivalent display of prefusion S ectodomain trimers may lead to elicitation of even broader Ab responses based on the greater sequence and structural conservation of the S2 subunit (i.e., the fusion machinery) among coronaviruses and the fact that it contains conserved epitopes that are targeted by neutralizing Abs such as the fusion peptide (Poh et al., 2020). Several reports of stabilized prefusion SARS-CoV-2 S variants provide promising antigens that can be used to test this hypothesis (Henderson et al., 2020; Hsieh et al., 2020; McCallum et al., 2020; Xiong et al., 2020). Although a single approach may be enough for generating protective responses against multiple closely related coronaviruses (e.g., sarbecoviruses), the genetic diversity across lineages and genera will likely necessitate a combination of several vaccine design approaches.

Here we leveraged the robustness and versatility of computationally designed two-component nanoparticles to rapidly generate promising SARS-CoV-2 vaccine candidates that are highly differentiated from many other candidates under development. Our results add another class I fusion protein to the growing list of antigens whose immunogenicity is enhanced through multivalent presentation on two-component nanoparticles. Continued development of such technology platforms could lead to vaccines that prevent the next pandemic rather than respond to it (Kanekiyo et al., 2019a; Kanekiyo and Graham, 2020).

## Supporting information

Supplementary Item 1

Supplementary Item 2

Supplementary Item 3

Supplementary Item 4

Supplementary Item 5

## ACKNOWLEDGMENTS

The authors gratefully acknowledge Caitlin Wolf, Naomi Wilcox, and Denise McCulloch for providing human convalescent serum samples; Adam Dingens and Jesse Bloom for facilitating acquisition of convalescent human serum samples and for comments on the manuscript; Maggie Ahlrichs for assistance with protein production; Karla-Luise Herpoldt for assistance and guidance with data analysis; and Ratika Krishnamurty for project management. This study was supported by the Bill & Melinda Gates Foundation (OPP1156262 to D.V. and N.P.K.; OPP1126258 to K.K.L.; H.Y.C.; and OPP1159947 to Kymab Ltd), a generous gift from the Audacious Project, a generous gift from Jodi Green and Mike Halperin, a generous gift from the Hanauer family, the Defense Threat Reduction Agency (HDTRA1-18-1-0001 to N.P.K.), the National Institute of General Medical Sciences (R01GM120553 to D.V.; R01GM099989 to K.K.L.), the National Institute of Allergy and Infectious Diseases (HHSN272201700059C to D.V.; 3U01AI42001-02S1 to M.P.), a Pew Biomedical Scholars Award (D.V.), Investigators in the Pathogenesis of Infectious Disease Awards from the Burroughs Wellcome Fund (D.V., M.P.), Fast Grants (D.V., M.P.) an Animal Models Contract HHSN272201700036I-75N93020F00001 (R.S.B), the University of Washington’s Proteomics Resource (UWPR95794), and the North Carolina Policy Collaboratory at the University of North Carolina at Chapel Hill with funding from the North Carolina Coronavirus Relief Fund established and appropriated by the North Carolina General Assembly (R.S.B).

## AUTHOR CONTRIBUTIONS

Conceptualization: A.C.W., B.F., D.V., N.P.K.; Modeling and design: A.C.W., D.V., N.P.K.; Formal Analysis: A.C.W., B.F., S.W., A.S., L.V.T., L.S., C.C., M.J.N., M.C.M, E.K. E.A.H., M.G., K.K.L., L.C., M.P., T.P.S., D.V., N.P.K.; Investigation: A.C.W., B.F., A.S., S.W., M.N.P., M.M., L.V.T., L.S., M.A.O., C.C., M.J.N., M.C.M., D.P., R.R., J.C.K., C.O., A.P., S.C., E.C.L., E.K., C.M.C., C.S., E.A.H., B.B., J.T.F., K.H.D., L.E.G., S.R.L., K.L.G., T.B.L; Resources: H.Y.C., P.K., M.P.; Writing – Original Draft: A.C.W., B.F., L.S., C.C., K.K.L., D.V., and N.P.K.; Writing – Review & Editing: All authors; Visualization: A.C.W., B.F., A.S., L.S., C.C., K.K.L., D.V., and N.P.K.; Supervision: D.H.F., R.S.B., P.K., K.K.L., L.C., M.P., T.P.S., D.V., and N.P.K.; Funding Acquisition: R.S.B., D.V., and N.P.K.

## STAR★METHODS

### KEY RESOURCES TABLE

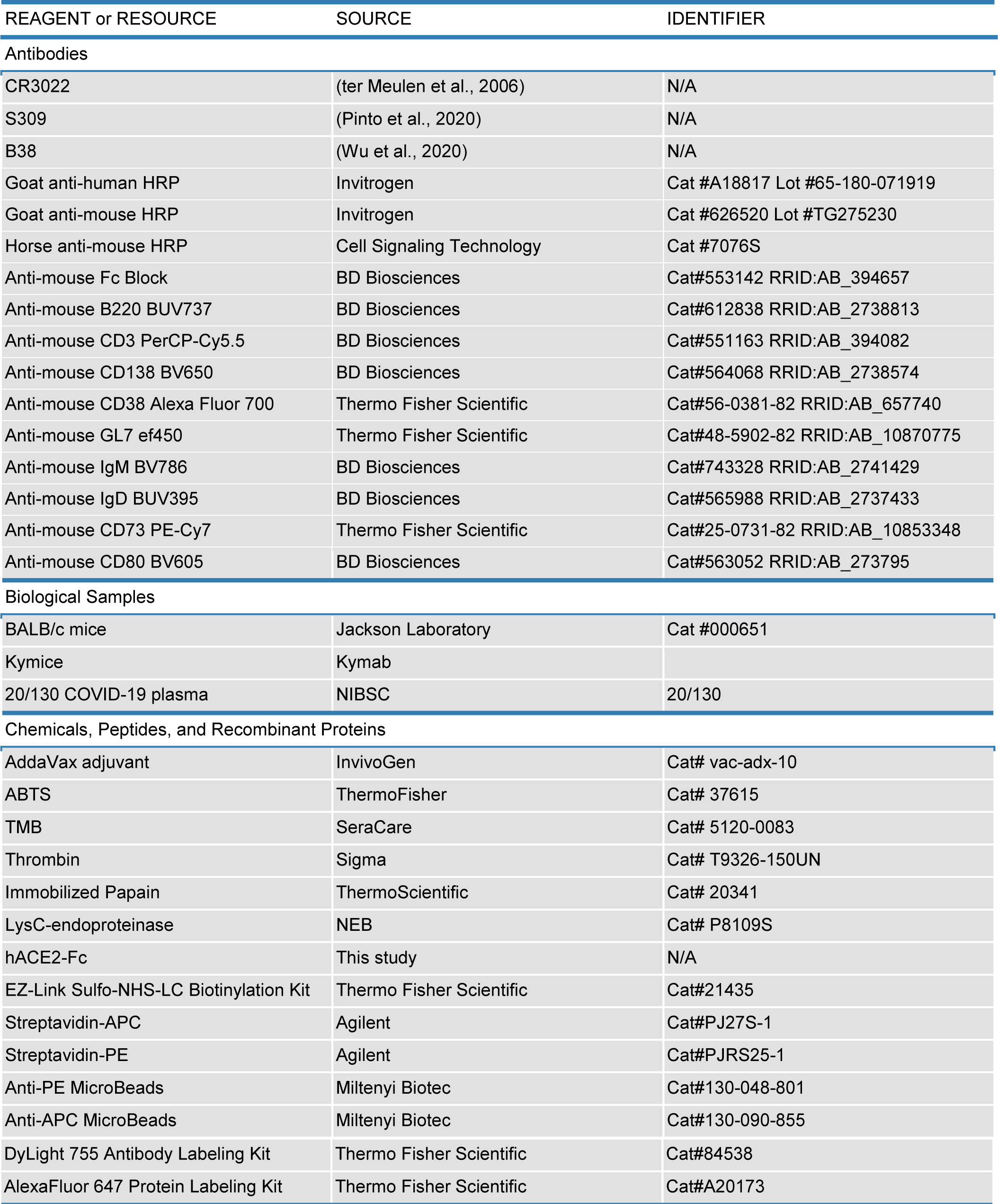
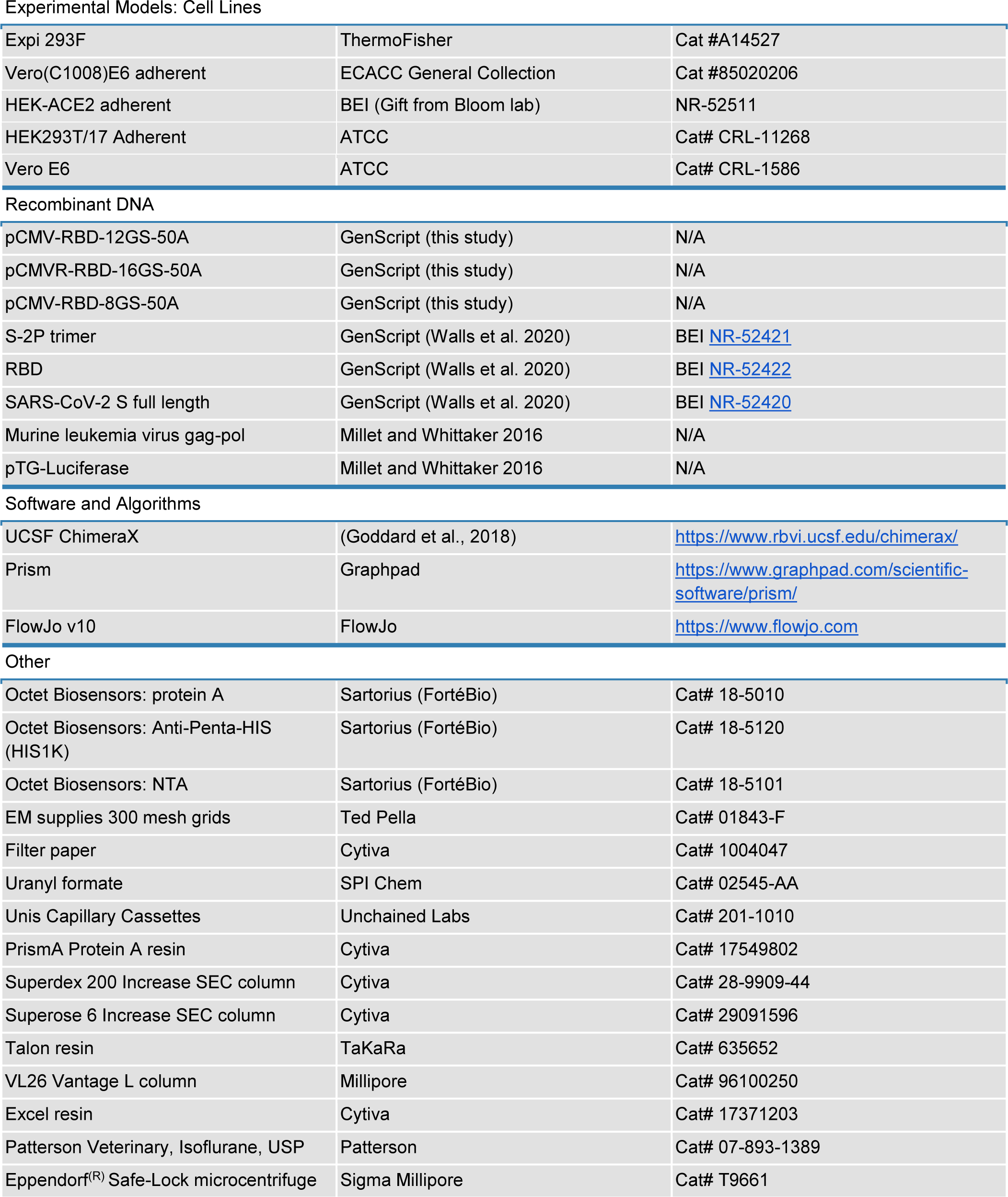
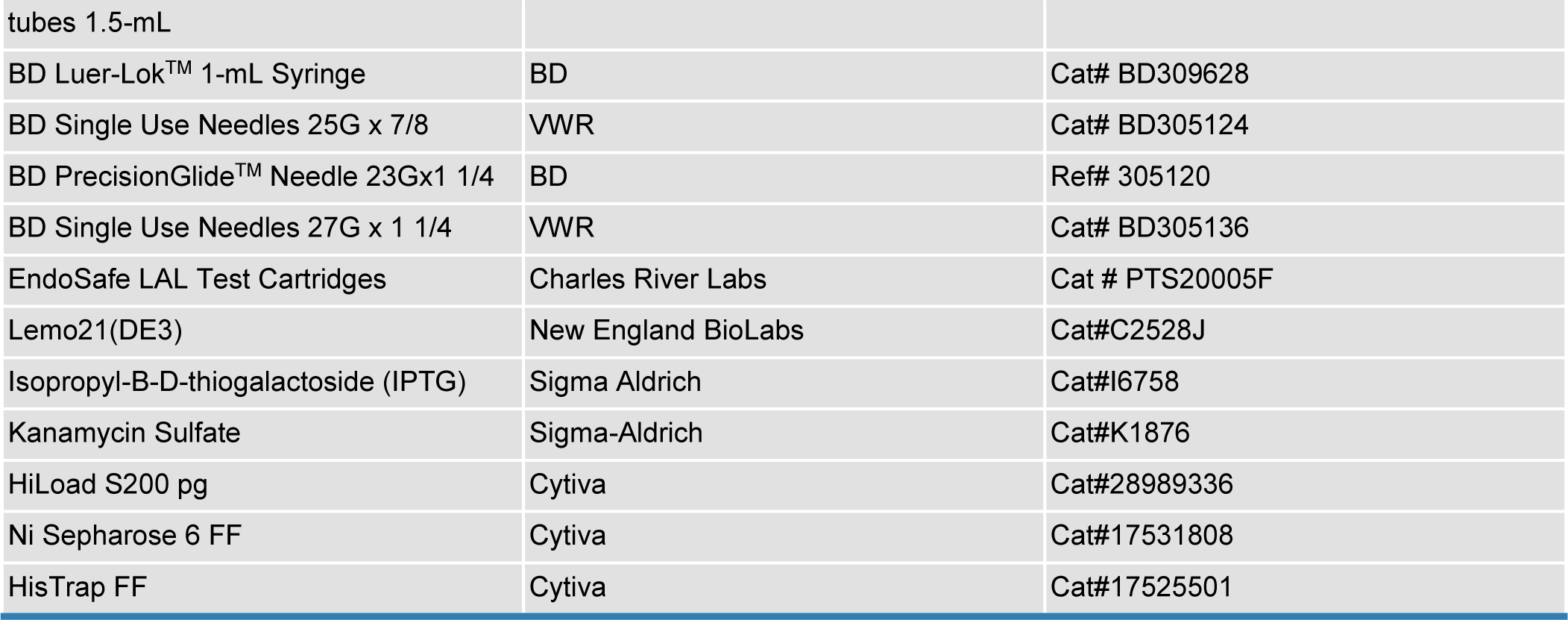

| REAGENT or RESOURCE | SOURCE | IDENTIFIER |
| --- | --- | --- |
| Antibodies |  |  |
| CR3022 | (ter Meulen et al., 2006) | N/A |
| S309 | (Pinto et al., 2020) | N/A |
| B38 | (Wu et al., 2020) | N/A |
| Goat anti-human HRP | Invitrogen | Cat #A18817 Lot #65-180-071919 |
| Goat anti-mouse HRP | Invitrogen | Cat #626520 Lot #TG275230 |
| Horse anti-mouse HRP | Cell Signaling Technology | Cat #7076S |
| Anti-mouse Fc Block | BD Biosciences | Cat#553142 RRID:AB_394657 |
| Anti-mouse B220 BUV737 | BD Biosciences | Cat#612838 RRID:AB_2738813 |
| Anti-mouse CD3 PerCP-Cy5.5 | BD Biosciences | Cat#551163 RRID:AB_394082 |
| Anti-mouse CD138 BV650 | BD Biosciences | Cat#564068 RRID:AB_2738574 |
| Anti-mouse CD38 Alexa Fluor 700 | Thermo Fisher Scientific | Cat#56-0381-82 RRID:AB_657740 |
| Anti-mouse GL7 ef450 | Thermo Fisher Scientific | Cat#48-5902-82 RRID:AB_10870775 |
| Anti-mouse IgM BV786 | BD Biosciences | Cat#743328 RRID:AB_2741429 |
| Anti-mouse IgD BUV395 | BD Biosciences | Cat#565988 RRID:AB_2737433 |
| Anti-mouse CD73 PE-Cy7 | Thermo Fisher Scientific | Cat#25-0731-82 RRID:AB_10853348 |
| Anti-mouse CD80 BV605 | BD Biosciences | Cat#563052 RRID:AB_273795 |
| Biological Samples |  |  |
| BALB/c mice | Jackson Laboratory | Cat #000651 |
| Kymice | Kymab |  |
| 20/130 COVID-19 plasma | NIBSC | 20/130 |
| Chemicals, Peptides, and Recombinant Proteins |  |  |
| AddaVax adjuvant | InvivoGen | Cat# vac-adx-10 |
| ABTS | ThermoFisher | Cat# 37615 |
| TMB | SeraCare | Cat# 5120-0083 |
| Thrombin | Sigma | Cat# T9326-150UN |
| Immobilized Papain | ThermoScientific | Cat# 20341 |
| LysC-endoproteinase | NEB | Cat# P8109S |
| hACE2-Fc | This study | N/A |
| EZ-Link Sulfo-NHS-LC Biotinylation Kit | Thermo Fisher Scientific | Cat#21435 |
| Streptavidin-APC | Agilent | Cat#PJ27S-1 |
| Streptavidin-PE | Agilent | Cat#PJRS25-1 |
| Anti-PE MicroBeads | Miltenyi Biotec | Cat#130-048-801 |
| Anti-APC MicroBeads | Miltenyi Biotec | Cat#130-090-855 |
| DyLight 755 Antibody Labeling Kit | Thermo Fisher Scientific | Cat#84538 |
| AlexaFluor 647 Protein Labeling Kit | Thermo Fisher Scientific | Cat#A20173 |
| Experimental Models: Cell Lines |  |  |
| Expi 293F | ThermoFisher | Cat #A14527 |
| Vero(C1008)E6 adherent | ECACC General Collection | Cat #85020206 |
| HEK-ACE2 adherent | BEI (Gift from Bloom lab) | NR-52511 |
| HEK293T/17 Adherent | ATCC | Cat# CRL-11268 |
| Vero E6 | ATCC | Cat# CRL-1586 |
| Recombinant DNA |  |  |
| pCMV-RBD-12GS-50A | GenScript (this study) | N/A |
| pCMVR-RBD-16GS-50A | GenScript (this study) | N/A |
| pCMV-RBD-8GS-50A | GenScript (this study) | N/A |
| S-2P trimer | GenScript (Walls et al. 2020) | BEI NR-52421 |
| RBD | GenScript (Walls et al. 2020) | BEI NR-52422 |
| SARS-CoV-2 S full length | GenScript (Walls et al. 2020) | BEI NR-52420 |
| Murine leukemia virus gag-pol | Millet and Whittaker 2016 | N/A |
| pTG-Luciferase | Millet and Whittaker 2016 | N/A |
| Software and Algorithms |  |  |
| UCSF ChimeraX | (Goddard et al., 2018) | https://www.rbvi.ucsf.edu/chimerax/ |
| Prism | Graphpad | https://www.graphpad.com/scientific-software/prism/ |
| FlowJo v10 | FlowJo | https://www.flowjo.com |
| Other |  |  |
| Octet Biosensors: protein A | Sartorius (FortéBio) | Cat# 18-5010 |
| Octet Biosensors: Anti-Penta-HIS (HIS1K) | Sartorius (FortéBio) | Cat# 18-5120 |
| Octet Biosensors: NTA | Sartorius (FortéBio) | Cat# 18-5101 |
| EM supplies 300 mesh grids | Ted Pella | Cat# 01843-F |
| Filter paper | Cytiva | Cat# 1004047 |
| Uranyl formate | SPI Chem | Cat# 02545-AA |
| Unis Capillary Cassettes | Unchained Labs | Cat# 201-1010 |
| Prisma Protein A resin | Cytiva | Cat# 17549802 |
| Superdex 200 Increase SEC column | Cytiva | Cat# 28-9909-44 |
| Superose 6 Increase SEC column | Cytiva | Cat# 29091596 |
| Talon resin | TaKaRa | Cat# 635652 |
| VL26 Vantage L column | Millipore | Cat# 96100250 |
| Excel resin | Cytiva | Cat# 17371203 |
| Patterson Veterinary, Isoflurane, USP | Patterson | Cat# 07-893-1389 |
| Eppendorf<sup>(R)</sup> Safe-Lock microcentrifuge | Sigma Millipore | Cat# T9661 |
| tubes 1.5-mL |  |  |
| BD Luer-Lok™ 1-mL Syringe | BD | Cat# BD309628 |
| BD Single Use Needles 25G x 7/8 | VWR | Cat# BD305124 |
| BD PrecisionGlide™ Needle 23Gx1 1/4 | BD | Ref# 305120 |
| BD Single Use Needles 27G x 1 1/4 | VWR | Cat# BD305136 |
| EndoSafe LAL Test Cartridges | Charles River Labs | Cat # PTS20005F |
| Lemo21(DE3) | New England BioLabs | Cat#C2528J |
| Isopropyl-B-D-thiogalactoside (IPTG) | Sigma Aldrich | Cat#I6758 |
| Kanamycin Sulfate | Sigma-Aldrich | Cat#K1876 |
| HiLoad S200 pg | Cytiva | Cat#28989336 |
| Ni Sepharose 6 FF | Cytiva | Cat#17531808 |
| HisTrap FF | Cytiva | Cat#17525501 |

## RESOURCE AVAILABILITY

### Lead Contact

Further information and requests for resources and reagents should be directed to and will be fulfilled by the Lead Contact, Neil P. King.

### Materials Availability

All reagents will be made available on request after completion of a Materials Transfer Agreement.

### Data and Code Availability

All data supporting the findings of this study are found within the paper and its Supplementary Information, and are available from the Lead Contact author upon request.

## EXPERIMENTAL MODEL AND SUBJECT DETAILS

### Cell lines

HEK293F is a female human embryonic kidney cell line transformed and adapted to grow in suspension (Life Technologies). HEK293F cells were grown in 293FreeStyle expression medium (Life Technologies), cultured at 37°C with 8% CO_2_ and shaking at 130 rpm. Expi293F™ cells are derived from the HEK293F cell line (Life Technologies). Expi293F™ cells were grown in Expi293™ Expression Medium (Life Technologies), cultured at 36.5°C with 8% CO_2_ and shaking at 150 rpm. VeroE6 is a female kidney epithelial cell from African green monkey. HEK293T/17 is a female human embryonic kidney cell line (ATCC). The HEK-ACE2 adherent cell line was obtained through BEI Resources, NIAID, NIH: Human Embryonic Kidney Cells (HEK-293T) Expressing Human Angiotensin-Converting Enzyme 2, HEK-293T-hACE2 Cell Line, NR-52511. All adherent cells were cultured at 37°C with 8% CO_2_ in flasks with DMEM + 10% FBS (Hyclone) + 1% penicillin-streptomycin. Cell lines other than Expi293F were not tested for mycoplasma contamination nor authenticated.

### Mice

Female BALB/c mice four weeks old were obtained from Jackson Laboratory, Bar Harbor, Maine. Animal procedures were performed under the approvals of the Institutional Animal Care and Use Committee of University of Washington, Seattle, WA, and University of North Carolina, Chapel Hill, NC. Kymab’s proprietary IntelliSelect™ Transgenic mouse platform, known as Darwin, has complete human antibody loci with a non-rearranged human antibody variable and constant germline repertoire. Consequently, the antibodies produced by these mice are fully human.

### Pigtail macaques

Two adult male Pigtail macaques (*Macaca nemestrina*) were immunized in this study. All animals were housed at the Washington National Primate Research Center (WaNPRC), an American Association for the Accreditation of Laboratory Animal Care International (AAALAC)-accredited institution, as previously described (Erasmus et al., 2020). All procedures performed on the animals were with the approval of the University of Washington's Institutional Animal Care and Use Committee (IACUC).

### Convalescent human sera

Samples collected between 1–60 days post infection from 31 individuals who tested positive for SARS-CoV-2 by PCR were profiled for anti-SARS-CoV-2 S antibody responses and the 29 with anti-S Ab responses were maintained in the cohort (Figures 4 and 5). Individuals were enrolled as part of the HAARVI study at the University of Washington in Seattle, WA. Baseline sociodemographic and clinical data for these individuals are summarized in Supplementary Item 5. This study was approved by the University of Washington Human Subjects Division Institutional Review Board (STUDY00000959 and STUDY00003376). All experiments were performed in at least two technical and two biological replicates (for ELISA and pseudovirus neutralization assays). One sample is the 20/130 COVID-19 plasma from NIBSC (https://nibsc.org/documents/ifu/20-130.pdf).

## METHOD DETAILS

### Plasmid construction

The SARS-CoV-2 RBD (BEI NR-52422) construct was synthesized by GenScript into pcDNA3.1-with an N-terminal mu-phosphatase signal peptide and a C-terminal octa-histidine tag (GHHHHHHHH). The boundaries of the construct are N-_328_RFPN_331_ and _528_KKST_531_-C (Walls et al., 2020). The SARS-CoV-2 S-2P ectodomain trimer (GenBank: YP_009724390.1, BEI NR-52420) was synthesized by GenScript into pCMV with an N-terminal mu-phosphatase signal peptide and a C-terminal TEV cleavage site (GSGRENLYPQG), T4 fibritin foldon (GGGSGYIPEAPRDGQAYVRKDGEWVLLSTPL), and octa-histidine tag (GHHHHHHHH) (Walls et al., 2020). The construct contains the 2P mutations (proline substitutions at residues 986 and 987; (Pallesen et al., 2017)) and an _682_SGAG_685_ substitution at the furin cleavage site. The SARS-CoV-2 RBD was genetically fused to the N terminus of the trimeric I53-50A nanoparticle component using linkers of 8, 12, or 16 glycine and serine residues. RBD-8GS- and RBD-12GS-I53-50A fusions were synthesized and cloned by Genscript into pCMV. The RBD-16GS-I53-50A fusion was cloned into pCMV/R using the Xba1 and AvrII restriction sites and Gibson assembly (Gibson et al., 2009). All RBD-bearing components contained an N-terminal mu-phosphatase signal peptide and a C-terminal octa-histidine tag. The macaque or human ACE2 ectodomain was genetically fused to a sequence encoding a thrombin cleavage site and a human Fc fragment at the C-terminal end. hACE2-Fc was synthesized and cloned by GenScript with a BM40 signal peptide. Plasmids were transformed into the NEB 5-alpha strain of *E. coli* (New England Biolabs) for subsequent DNA extraction from bacterial culture (NucleoBond Xtra Midi kit) to obtain plasmid for transient transfection into Expi293F cells. The amino acid sequences of all novel proteins used in this study can be found in Supplementary Item 1.

### Transient transfection

SARS-CoV-2 S and ACE2-Fc proteins were produced in Expi293F cells grown in suspension using Expi293F expression medium (Life Technologies) at 33°C, 70% humidity, 8% CO_2_ rotating at 150 rpm. The cultures were transfected using PEI-MAX (Polyscience) with cells grown to a density of 3.0 million cells per mL and cultivated for 3 days. Supernatants were clarified by centrifugation (5 minutes at 4000 rcf), addition of PDADMAC solution to a final concentration of 0.0375% (Sigma Aldrich, #409014), and a second spin (5 minutes at 4000 rcf).

Genes encoding CR3022 heavy and light chains were ordered from GenScript and cloned into pCMV/R. Antibodies were expressed by transient co-transfection of both heavy and light chain plasmids in Expi293F cells using PEI MAX (Polyscience) transfection reagent. Cell supernatants were harvested and clarified after 3 or 6 days as described above.

### Protein purification

Proteins containing His tags were purified from clarified supernatants via a batch bind method where each clarified supernatant was supplemented with 1 M Tris-HCl pH 8.0 to a final concentration of 45 mM and 5 M NaCl to a final concentration of ~310 mM. Talon cobalt affinity resin (Takara) was added to the treated supernatants and allowed to incubate for 15 minutes with gentle shaking. Resin was collected using vacuum filtration with a 0.2 µm filter and transferred to a gravity column. The resin was washed with 20 mM Tris pH 8.0, 300 mM NaCl, and the protein was eluted with 3 column volumes of 20 mM Tris pH 8.0, 300 mM NaCl, 300 mM imidazole. The batch bind process was then repeated and the first and second elutions combined. SDS-PAGE was used to assess purity. RBD-I53-50A fusion protein IMAC elutions were concentrated to >1 mg/mL and subjected to three rounds of dialysis into 50 mM Tris pH 7, 185 mM NaCl, 100 mM Arginine, 4.5% glycerol, and 0.75% w/v 3-[(3-cholamidopropyl)dimethylammonio]-1-propanesulfonate (CHAPS) in a hydrated 10K molecular weight cutoff dialysis cassette (Thermo Scientific). S-2P IMAC elution fractions were concentrated to ~1 mg/mL and dialyzed three times into 50 mM Tris pH 8, 150 mM NaCl, 0.25% L-Histidine in a hydrated 10K molecular weight cutoff dialysis cassette (Thermo Scientific). Due to inherent instability, the S-2P trimer was immediately flash frozen and stored at −80°C.

Clarified supernatants of cells expressing monoclonal antibodies and human or macaque ACE2-Fc were purified using a MabSelect PrismA 2.6×5 cm column (Cytiva) on an AKTA Avant150 FPLC (Cytiva). Bound antibodies were washed with five column volumes of 20 mM NaPO4, 150 mM NaCl pH 7.2, then five column volumes of 20 mM NaPO_4_, 1 M NaCl pH 7.4 and eluted with three column volumes of 100 mM glycine at pH 3.0. The eluate was neutralized with 2 M Trizma base to 50 mM final concentration. SDS-PAGE was used to assess purity.

Recombinant S309 was expressed as a Fab in expiCHO cells transiently co-transfected with plasmids expressing the heavy and light chain, as described above (see Transient transfection) (Stettler et al., 2016). The protein was affinity-purified using a HiTrap Protein A Mab select Xtra column (Cytiva) followed by desalting against 20 mM NaPO_4_, 150 mM NaCl pH 7.2 using a HiTrap Fast desalting column (Cytiva). The protein was sterilized with a 0.22 µm filter and stored at 4°C until use.

### Microbial protein expression and purification

The I53-50A and I53-50B.4.PT1 proteins were expressed in Lemo21(DE3) (NEB) in LB (10 g Tryptone, 5 g Yeast Extract, 10 g NaCl) grown in 2 L baffled shake flasks or a 10 L BioFlo 320 Fermenter (Eppendorf). Cells were grown at 37°C to an OD600 ~0.8, and then induced with 1 mM IPTG. Expression temperature was reduced to 18°C and the cells shaken for ~16 h. The cells were harvested and lysed by microfluidization using a Microfluidics M110P at 18,000 psi in 50 mM Tris, 500 mM NaCl, 30 mM imidazole, 1 mM PMSF, 0.75% CHAPS. Lysates were clarified by centrifugation at 24,000 g for 30 min and applied to a 2.6×10 cm Ni Sepharose 6 FF column (Cytiva) for purification by IMAC on an AKTA Avant150 FPLC system (Cytiva). Protein of interest was eluted over a linear gradient of 30 mM to 500 mM imidazole in a background of 50 mM Tris pH 8, 500 mM NaCl, 0.75% CHAPS buffer. Peak fractions were pooled, concentrated in 10K MWCO centrifugal filters (Millipore), sterile filtered (0.22 μm) and applied to either a Superdex 200 Increase 10/300, or HiLoad S200 pg GL SEC column (Cytiva) using 50 mM Tris pH 8, 500 mM NaCl, 0.75% CHAPS buffer. I53-50A elutes at ∼0.6 column volume (CV). I53-50B.4PT1 elutes at ~0.45 CV. After sizing, bacterial-derived components were tested to confirm low levels of endotoxin before using for nanoparticle assembly.

### *In vitro* nanoparticle assembly

Total protein concentration of purified individual nanoparticle components was determined by measuring absorbance at 280 nm using a UV/vis spectrophotometer (Agilent Cary 8454) and calculated extinction coefficients (Gasteiger et al., 2005). The assembly steps were performed at room temperature with addition in the following order: RBD-I53-50A trimeric fusion protein, followed by additional buffer as needed to achieve desired final concentration, and finally I53-50B.4PT1 pentameric component (in 50 mM Tris pH 8, 500 mM NaCl, 0.75% w/v CHAPS), with a molar ratio of RBD-I53-50A:I53-B.4PT1 of 1.1:1. In order to produce partial valency RBD-I53-50 nanoparticles (50% RBD-I53-50), both RBD-I53-50A and unmodified I53-50A trimers (in 50 mM Tris pH 8, 500 mM NaCl, 0.75% w/v CHAPS) were added in a slight molar excess (1.1×) to I53-50B.4PT1. All RBD-I53-50 *in vitro* assemblies were incubated at 2-8°C with gentle rocking for at least 30 minutes before subsequent purification by SEC in order to remove residual unassembled component. Different columns were utilized depending on purpose: Superose 6 Increase 10/300 GL column was used analytically for nanoparticle size estimation, a Superdex 200 Increase 10/300 GL column used for small-scale pilot assemblies, and a HiLoad 26/600 Superdex 200 pg column used for nanoparticle production. Assembled particles elute at ~11 mL on the Superose 6 column and in the void volume of Superdex 200 columns. Assembled nanoparticles were sterile filtered (0.22 µm) immediately prior to column application and following pooling of fractions.

### hACE2-Fc and CR3022 digestion

hACE2-Fc was digested with thrombin protease (Sigma Aldrich) in the presence of 2.5 mM CaCl_2_ at a 1:300 w/w thrombin:protein ratio. The reaction was incubated at ambient temperature for 16–18 hours with gentle rocking. Following incubation, the reaction mixture was concentrated using Ultracel 10K centrifugal filters (Millipore Amicon Ultra) and sterile filtered (0.22 µM). Cleaved hACE2 monomer was separated from uncleaved hACE2-Fc and the cleaved Fc regions using Protein A purification (see Protein purification above) on a HiScreen MabSelect SuRe column (Cytiva) using an ӒKTA avant 25 FPLC (Cytiva). Cleaved hACE2 monomer was collected in the flow through, sterile filtered (0.22 µm), and quantified by UV/vis.

LysC (New England BioLabs) was diluted to 10 ng/µL in 10 mM Tris pH 8 and added to CR3022 IgG at 1:2000 w/w LysC:IgG and subsequently incubated for 18 hours at 37°C with orbital shaking at 230 rpm. The cleavage reaction was concentrated using Ultracel 10K centrifugal filters (Millipore Amicon Ultra) and sterile filtered (0.22 µM). Cleaved CR3022 mAb was separated from uncleaved CR3022 IgG and the Fc portion of cleaved IgG, using Protein A purification as described above. Cleaved CR3022 was collected in the flow through, sterile filtered (0.22 µm), and quantified by UV/vis.

### Bio-layer interferometry (antigenicity)

Antigenicity assays were performed and analyzed using BLI on an Octet Red 96 System (Pall Forté Bio/Sartorius) at ambient temperature with shaking at 1000 rpm. RBD-I53-50A trimeric components and monomeric RBD were diluted to 40 µg/mL in Kinetics buffer (1x HEPES-EP+ (Pall Forté Bio), 0.05% nonfat milk, and 0.02% sodium azide). Monomeric hACE2 and CR3022 Fab were diluted to 750 nM in Kinetics buffer and serially diluted three-fold for a final concentration of 3.1 nM. Reagents were applied to a black 96-well Greiner Bio-one microplate at 200 μL per well as described below. RBD-I53-50A components or monomeric RBD were immobilized onto Anti-Penta-HIS (HIS1K) biosensors per manufacturer instructions (Forté Bio) except using the following sensor incubation times. HIS1K biosensors were hydrated in water for 10 minutes, and were then equilibrated in Kinetics buffer for 60 seconds. The HIS1K tips were loaded with diluted trimeric RBD-I53-50A component or monomeric RBD for 150 seconds and washed with Kinetics buffer for 300 seconds. The association step was performed by dipping the HIS1K biosensors with immobilized immunogen into diluted hACE2 monomer or CR3022 Fab for 600 seconds, then dissociation was measured by inserting the biosensors back into Kinetics buffer for 600 seconds. The data were baseline subtracted and the plots fitted using the Pall FortéBio/Sartorius analysis software (version 12.0). Plots in Figure S2 show the association and dissociation steps.

### Bio-layer interferometry (accessibility)

Binding of mACE2-Fc, CR3022 IgG, and S309 IgG to monomeric RBD, RBD-I53-50A trimers, and RBD-I53-50 nanoparticles was analyzed for accessibility experiments and real-time stability studies using an Octet Red 96 System (Pall FortéBio/Sartorius) at ambient temperature with shaking at 1000 rpm. Protein samples were diluted to 100 nM in Kinetics buffer. Buffer, immunogen, and analyte were then applied to a black 96-well Greiner Bio-one microplate at 200 µL per well. Protein A biosensors (FortéBio/Sartorius) were first hydrated for 10 minutes in Kinetics buffer, then dipped into either mACE2-Fc, CR3022, or S309 IgG diluted to 10 µg/mL in Kinetics buffer in the immobilization step. After 500 seconds, the tips were transferred to Kinetics buffer for 60 seconds to reach a baseline. The association step was performed by dipping the loaded biosensors into the immunogens for 300 seconds, and subsequent dissociation was performed by dipping the biosensors back into Kinetics buffer for an additional 300 seconds. The data were baseline subtracted prior to plotting using the FortéBio analysis software (version 12.0). Plots in Figure 2 show the 600 seconds of association and dissociation.

### Negative stain electron microscopy

RBD-I53-50 nanoparticles were first diluted to 75 µg/mL in 50 mM Tris pH 7, 185 mM NaCl, 100 mM Arginine, 4.5% v/v Glycerol, 0.75% w/v CHAPS, and S-2P protein was diluted to 0.03 mg/mL in 50 mM Tris pH 8, 150 mM NaCl, 0.25% L-Histidine prior to application of 3 µL of sample onto freshly glow-discharged 300-mesh copper grids. Sample was incubated on the grid for 1 minute before the grid was dipped in a 50 µL droplet of water and excess liquid blotted away with filter paper (Whatman). The grids were then dipped into 6 µL of 0.75% w/v uranyl formate stain. Stain was blotted off with filter paper, then the grids were dipped into another 6 µL of stain and incubated for ~70 seconds. Finally, the stain was blotted away and the grids were allowed to dry for 1 minute. Prepared grids were imaged in a Talos model L120C electron microscope at 45,000x (nanoparticles) or 92,000x magnification (S-2P).

### Dynamic light scattering

Dynamic Light Scattering (DLS) was used to measure hydrodynamic diameter (Dh) and % Polydispersity (%Pd) of RBD-I53-50 nanoparticle samples on an UNcle Nano-DSF (UNchained Laboratories). Sample was applied to a 8.8 µL quartz capillary cassette (UNi, UNchained Laboratories) and measured with 10 acquisitions of 5 seconds each, using auto-attenuation of the laser. Increased viscosity due to 4.5% v/v glycerol in the RBD nanoparticle buffer was accounted for by the UNcle Client software in Dh measurements.

### Guanidine HCl denaturation

Monomeric RBD, RBD-I53-50A fusion proteins, and RBD-I53-50 nanoparticle immunogens were diluted to 2.5 µM in 50 mM Tris pH 7.0, 185 mM NaCl, 100 mM Arginine, 4.5% v/v glycerol, 0.75% w/v CHAPS, and guanidine chloride [GdnHCl] ranging from 0 M to 6.5 M, increasing in 0.25 M increments, and prepared in triplicate. S-2P trimer was also diluted to 2.5 µM using 50 mM Tris pH 8, 150 mM NaCl, 0.25% L-Histidine, and the same GuHCl concentration range. Dilutions were mixed 10x by pipetting. The samples were then incubated 18–19 hours at ambient temperature. Using a Nano-DSF (UNcle, UNchained Laboratories) and an 8.8 µL quartz capillary cassette (UNi, UNchained Laboratories), fluorescence spectra were collected in triplicate, exciting at 266 nm and measuring emission from 200 nm to 750 nm at 25°C.

### Endotoxin measurements

Endotoxin levels in protein samples were measured using the EndoSafe Nexgen-MCS System (Charles River). Samples were diluted 1:50 or 1:100 in Endotoxin-free LAL reagent water, and applied into wells of an EndoSafe LAL reagent cartridge. Charles River EndoScan-V software was used to analyze endotoxin content, automatically back-calculating for the dilution factor. Endotoxin values were reported as EU/mL which were then converted to EU/mg based on UV/vis measurements. Our threshold for samples suitable for immunization was <50 EU/mg.

### UV/vis

Ultraviolet-visible spectrophotometry (UV/vis) was measured using an Agilent Technologies Cary 8454. Samples were applied to a 10 mm, 50 µL quartz cell (Starna Cells, Inc.) and absorbance was measured from 180 to 1000 nm. Net absorbance at 280 nm, obtained from measurement and single reference wavelength baseline subtraction, was used with calculated extinction coefficients and molecular weights to obtain protein concentration. The ratio of absorbance at 320/280 nm was used to determine relative aggregation levels in real-time stability study samples. Samples were diluted with respective purification/instrument blanking buffers to obtain an absorbance between 0.1 and 1.0. All data produced from the UV/vis instrument was processed in the 845x UV/visible System software.

### Glycan profiling

To identify site-specific glycosylation profiles, including glycoform distribution and occupancy determination, a bottom up mass spectrometry (MS) approach was utilized. Aliquots of 1 mg/mL monomeric, 8GS, 12GS and 16GS RBD protein were prepared to evaluate the glycosylation profiles at N331 and N343 of the four RBD variants. Comprehensive glycoprofiling on the stabilized Spike ectodomain (S-2P) was performed in parallel using 1.5 mg/mL SARS-CoV-2 S-2P protein. All the samples were denatured in a solution containing 25 mM Tris (pH 8.0), 7 M guanidinium chloride (GdnHCl) and 50 mM dithiothreitol (DTT) at 90°C for 30 minutes. Reduced cysteines were alkylated by adding fresh iodoacetamide (IAA) to 100 mM and incubating at room temperature for 1 hour in the dark. 50 mM excess DTT was then added to quench the remaining IAA. The GndHCl concentration was reduced to 0.6 M by diluting the samples 11-fold with a 10 mM Tris (pH 8.0), 2 mM calcium chloride solution. Each sample was then split in half. One half (275 µL) was mixed with 10 units of recombinant Peptide N-glycanase F (GST-PNGase F) (Krenkova et al., 2013) and incubated at 37°C for 1 hour in order to convert glycosylated Asn into deglycosylated Asp.

Protease digestions were performed in the following manner: all RBD samples and one S-2P sample were digested with Lys-C at a ratio of 1:40 (w/w) for RBD and 1:30 (w/w) for S-2P for 4 hours at 37°C, followed by Glu-C digestion overnight at the same ratios and conditions. The other three S-2P samples were digested with trypsin, chymotrypsin and alpha lytic protease, respectively, at a ratio of 1:30 (w/w) overnight at 37°C. All the digestion proteases used were MS grade (Promega). The next day, the digestion reactions were quenched by 0.02% formic acid (FA, Optima™, Fisher).

The glycoform determination of four S-2P samples was performed by nano LC-MS using an Orbitrap Fusion™ mass spectrometer (Thermo Fisher). The digested samples were desalted by Sep-Pak C18 cartridges (Waters) following the manufacturer’s suggested protocol. A 2 cm trapping column and a 35 cm analytical column were freshly prepared in fused silica (100 µm ID) with 5 µM ReproSil-Pur C18 AQ beads (Dr. Maisch). 8 µL sample was injected and run by a 60-minute linear gradient from 2% to 30% acetonitrile in 0.1% FA, followed by 10 minutes of 80% acetonitrile. An EThcD method was optimized as followed: ion source: 2.1 kV for positive mode; ion transfer tube temperature: 350 °C; resolution: MS^1^ = 120000, MS^2^ = 30000; AGC target: MS^1^ = 2e^5^, MS^2^ = 1e^5^; and injection time: MS^1^ = 50 ms, MS^2^ = 60 ms.

Glycopeptide data were visualized and processed by Byonic™ and Byologic™ (Version 3.8, Protein Metrics Inc.) using a 6 ppm precursor and 10 ppm fragment mass tolerance. Glycopeptides were searched using the N-glycan 309 mammalian database in Protein Metrics PMI-Suite and scored based on the assignment of correct c- and z-fragment ions. The true-positive entities were further validated by the presence of glycan oxonium ions m/z at 204 (HexNAc ions) and 366 (HexNAcHex ions) and the absence in its corresponding spectrum in the deglycosylated sample. The relative abundance of each glycoform was determined by the peak area analyzed in Byologic™. Glycoforms were categorized in Oligo (Oligomannose), Hybrid, and Complex as well as subtypes in Complex, described in the previous study (Watanabe et al., 2020). HexNAc(2)Hex(9-5) is M(annose)9 to M5; HexNAc(3)Hex(5-6) is classified as Hybrid; HexNAc(3)Hex(3-4)X is A1 subtype; HexNAc(4)X is A2/A1B; HexNAc(5)X is A3/A2B and HexNAc(6)X is A4/A3B subtype. Hybrid and Complex forms with fucosylation are separately listed as FHybrid and FComplex (eg. FA1), respectively.

Glycan occupancy analysis and glycoform determination of the four RBD variants were performed by LC-MS on the Synapt G2-Si™ TOF mass spectrometer coupled to an Acquity UPLC system (Waters). Samples were resolved over a Waters CSH C18 1 x 100 mm 1.7 µm column with a linear gradient from 3% to 40% B over 30 minutes (A: 98% water, 2% acetonitrile, 0.1% FA; B: 100% acetonitrile, 0.1% FA). Data dependent acquisition (DDA) method was utilized with precursor mass range 300-2000, MS/MS mass range 50-2000 and a collision energy ramped from 70 to 100 V. Chromatographic peaks for the most abundant and non-overlapped isotopic peaks were determined and integrated with MassLynx (Waters). All the water and organic solvents used, unless specifically stated, were MS grade (OptimaTM, Fisher). The peak area ratio of the non-glycosylated (Asn) to the deglycosylated (Asp) glycopeptide was used to measure the glycan occupancy at each site.

### Hydrogen/Deuterium-exchange mass spectrometry

3 µg of monomeric RBD and RBD-8GS-I53-50A were incubated and H/D exchanged (HDX) in the deuteration buffer (pH* 7.6, 85% D_2_O, Cambridge Isotope Laboratories, Inc.) for 3, 60, 1800, and 72000 seconds, respectively, at 23°C. Samples were subsequently mixed 1:1 with ice-cold quench buffer (200 mM tris(2-chlorethyl) phosphate (TCEP), 8 M Urea, 0.2% formic acid) for a final pH 2.5 and immediately flash frozen in liquid nitrogen. Samples were in-line pepsin digested and analyzed by LC-MS-IMS on Synapt G2-Si™ TOF mass spectrometer (Waters) as previously described (Verkerke et al., 2016) with an 18 minute gradient applied. A fully deuteration control was made by collecting the pepsin digest eluate from an undeuterated sample LC-MS run, drying by speedvac, incubating in deuteration buffer for 1 hour at 85°C, and quenching the same as all other HDX samples. Internal exchange standards (Pro-Pro-Pro-Ile [PPPI] and Pro-Pro-Pro-Phe [PPPF]) were added in each sample to ensure consistent labeling conditions for all samples (Zhang et al., 2012). Pepsin digests for undeuterated samples were also analyzed by nano LC-MS using an Orbitrap Fusion™ mass spectrometer (Thermo Fisher) with the settings as described above for glycoprofiling. The data was then processed by Byonic™ to obtain the peptide reference list. Peptides were manually validated using DriftScope™ (Waters) and identified with orthogonal retention time (rt) and drift time (dt) coordinates. Deuterium uptake analysis was performed with HX-Express v2 (Guttman et al., 2013; Weis et al., 2006). Peaks were identified from the peptide spectra with binomial fitting applied. The deuterium uptake level was normalized relative to fully deuterated standards.

### Mouse immunizations and challenge

Female BALB/c (Stock: 000651) mice were purchased at the age of four weeks from The Jackson Laboratory, Bar Harbor, Maine, and maintained at the Comparative Medicine Facility at the University of Washington, Seattle, WA, accredited by the American Association for the Accreditation of Laboratory Animal Care International (AAALAC). At six weeks of age, 10 mice per dosing group were vaccinated with a prime immunization, and three weeks later mice were boosted with a second vaccination. Prior to inoculation, immunogen suspensions were gently mixed 1:1 vol/vol with AddaVax adjuvant (Invivogen, San Diego, CA) to reach a final concentration of 0.009 or 0.05 mg/mL antigen. Mice were injected intramuscularly into the gastrocnemius muscle of each hind leg using a 27-gauge needle (BD, San Diego, CA) with 50 μL per injection site (100 μL total) of immunogen under isoflurane anesthesia. To obtain sera all mice were bled two weeks after prime and boost immunizations. Blood was collected via submental venous puncture and rested in 1.5 mL plastic Eppendorf tubes at room temperature for 30 minutes to allow for coagulation. Serum was separated from hematocrit via centrifugation at 2000 g for 10 minutes. Complement factors and pathogens in isolated serum were heat-inactivated via incubating serum at 56°C for 60 minutes. Serum was stored at 4°C or −80°C until use. Six weeks post-boost, mice were exported from Comparative Medicine Facility at the University of Washington, Seattle, WA to an AAALAC accredited Animal Biosafety Level 3 (ABSL3) Laboratory at the University of North Carolina, Chapel Hill. After a 7-day acclimation time, mice were anesthetized with a mixture of ketamine/xylazine and challenged intranasally with 10^5^ plaque-forming units (pfu) of mouse-adapted SARS-CoV-2 MA strain for the evaluation of vaccine efficacy (IACUC protocol 20-114.0). After infection, body weight was monitored daily until the termination of the study two days post-infection, when lung and nasal turbinate tissues were harvested to evaluate the viral load by plaque assay. All experiments were conducted at the University of Washington, Seattle, WA, and University of North Carolina, Chapel Hill, NC according to approved Institutional Animal Care and Use Committee protocols.

### Immunization (Kymab Darwin mice)

Kymab Darwin mice (a mix of males and females, 10 weeks of age), 5 mice per dosing group, were vaccinated with a prime immunization and three weeks later boosted with a second vaccination. Prior to inoculation, immunogen suspensions were gently mixed 1:1 vol/vol with AddaVax adjuvant (Invivogen) to reach a final concentration of 0.009 or 0.05 mg/mL antigen. Mice were injected intramuscularly into the tibialis muscle of each hind leg using a 30-gauge needle (BD) with 20 μL per injection site (40 μL total) of immunogen under isoflurane anesthesia. A final boost was administered intravenously (50 uL) with no adjuvant at week 7. Mice were sacrificed 5 days later under UK Home Office Schedule 1 (rising concentration of CO_2_) and spleen, lymph nodes, and bone marrow cryopreserved. Whole blood (0.1 ml) was collected 2 weeks after each dose (weeks 0, 2, 5, and week 8 terminal bleed). Serum was separated from hematocrit via centrifugation at 2000 g for 10 minutes. Serum was stored at −20°C and was used to monitor titers by ELISA. All mice were maintained and all procedures carried out under United Kingdom Home Office License 70/8718 and with the approval of the Wellcome Trust Sanger Institute Animal Welfare and Ethical Review Body.

### ELISA

For anti-S-2P ELISA, 25 μL of 2 μg/mL S-2P was plated onto 384-well Nunc Maxisorp (ThermoFisher) plates in PBS and sealed overnight at 4°C. The next day plates were washed 4x in Tris Buffered Saline Tween (TBST) using a plate washer (BioTek) and blocked with 2% BSA in TBST for 1 h at 37°C. Plates were washed 4x in TBST and 1:5 serial dilutions of mouse, NHP, or human sera were made in 25 μL TBST starting at 1:25 or 1:50 and incubated at 37°C for 1 h. Plates were washed 4x in TBST, then anti-mouse (Invitrogen) or anti-human (Invitrogen) horseradish peroxidase-conjugated antibodies were diluted 1:5,000 and 25 μL added to each well and incubated at 37°C for 1 h. Plates were washed 4x in TBST and 25 μL of TMB (SeraCare) was added to every well for 5 min at room temperature. The reaction was quenched with the addition of 25 μL of 1N HCl. Plates were immediately read at 450 nm on a VarioSkanLux plate reader (ThermoFisher) and data plotted and fit in Prism (GraphPad) using nonlinear regression sigmoidal, 4PL, X is log(concentration) to determine EC_50_ values from curve fits.

### Pseudovirus production

MLV-based SARS-CoV-2 S, SARS-CoV S, and WIV-1 pseudotypes were prepared as previously described (Millet and Whittaker, 2016; Walls et al., 2020). Briefly, HEK293T cells were co-transfected using Lipofectamine 2000 (Life Technologies) with an S-encoding plasmid, an MLV Gag-Pol packaging construct, and the MLV transfer vector encoding a luciferase reporter according to the manufacturer’s instructions. Cells were washed 3x with Opti-MEM and incubated for 5 h at 37°C with transfection medium. DMEM containing 10% FBS was added for 60 h. The supernatants were harvested by a 2,500 g spin, filtered through a 0.45 μm filter, concentrated with a 100 kDa membrane for 10 min at 2,500 g and then aliquoted and placed at −80°C.

### Pseudovirus entry and serum neutralization assays

HEK-hACE2 cells were cultured in DMEM with 10% FBS (Hyclone) and 1% PenStrep with 8% CO_2_ in a 37°C incubator (ThermoFisher). One day prior to infection, 40 μL of poly-lysine (Sigma) was placed into 96-well plates and incubated with rotation for 5 min. Poly-lysine was removed, plates were dried for 5 min then washed 1x with DMEM prior to plating cells. The following day, cells were checked to be at 80% confluence. In a half-area 96-well plate a 1:3 serial dilution of sera was made in DMEM starting between 1:3 and 1:66 initial dilution in 22 μL final volume. 22 μL of pseudovirus was then added to the serial dilution and incubated at room temperature for 30-60 min. HEK-hACE2 plate media was removed and 40 μL of the sera/virus mixture was added to the cells and incubated for 2 h at 37°C with 8% CO_2_. Following incubation, 40 μL 20% FBS and 2% PenStrep containing DMEM was added to the cells for 48 h. Following the 48-h infection, One-Glo-EX (Promega) was added to the cells in half culturing volume (40 μL added) and incubated in the dark for 5 min prior to reading on a Varioskan LUX plate reader (ThermoFisher). Measurements were done on all ten mouse sera samples from each group in at least duplicate. Relative luciferase units were plotted and normalized in Prism (GraphPad) using a zero value of cells alone and a 100% value of 1:2 virus alone. Nonlinear regression of log(inhibitor) vs. normalized response was used to determine IC_50_ values from curve fits. Mann-Whitney tests were used to compare two groups to determine whether they were statistically different.

### Live virus production

SARS-CoV-2-nanoLuc virus (WA1 strain) in which ORF7 was replaced by nanoluciferase gene (nanoLuc), and mouse-adapted SARS-CoV-2 (SARS-CoV-2 MA) (Dinnon et al., 2020) were generated by the coronavirus reverse genetics system described previously (Hou et al., 2020). Recombinant viruses were generated in Vero E6 cells (ATCC-CRL1586) grown in DMEM high glucose media (Gibco #11995065) supplemented with 10% Hyclone Fetal Clone II (GE #SH3006603HI), 1% non-essential amino acid, and 1% Pen/Strep in a 37°C +5% CO_2_ incubator. To generate recombinant SARS-CoV-2, seven DNA fragments which collectively encode the full-length genome of SARS-CoV-2 flanked by a 5′ T7 promoter and a 3′ polyA tail were ligated and transcribed *in vitro*. The transcribed RNA was electroporated into Vero E6 cells to generate a P0 virus stock. The seed virus was amplified twice in Vero E6 cells at low moi for 48 h to create a working stock which was titered by plaque assay (Hou et al., 2020). All the live virus experiments, including the ligation and electroporation steps, were performed under biosafety level 3 (BSL-3) conditions at negative pressure, by operators in Tyvek suits wearing personal powered-air purifying respirators.

### Luciferase-based serum neutralization assay, SARS-CoV-2-nanoLuc

Vero E6 cells were seeded at 2×10^4^ cells/well in a 96-well plate 24 h before the assay. One hundred pfu of SARS-CoV-2-nanoLuc virus (Hou et al., 2020) were mixed with serum at 1:1 ratio and incubated at 37°C for 1 h. An 8-point, 3-fold dilution curve was generated for each sample with starting concentration at 1:20 (standard) or 1:2000 (high neutralizer). Virus and serum mix was added to each well and incubated at 37°C + 5% CO_2_ for 48 h. Luciferase activities were measured by Nano-Glo Luciferase Assay System (Promega, WI) following manufacturer protocol using SpectraMax M3 luminometer (Molecular Device). Percent inhibition and 50% inhibition concentration (IC_50_) were calculated by the following equation: [1-(RLU with sample/ RLU with mock treatment)] x 100%. Fifty percent inhibition titer (IC_50_) was calculated in GraphPad Prism 8.3.0 by fitting the data points using a sigmoidal dose-response (variable slope) curve.

### Tetramer production

Recombinant SARS-CoV-2 S-2P trimer was biotinylated using the EZ-Link Sulfo-NHS-LC Biotinylation Kit (ThermoFisher) and tetramerized with streptavidin-APC (Agilent) as previously described (Krishnamurty et al., 2016; Taylor et al., 2012). The RBD domain of SARS-CoV-2 S was biotinylated and tetramerized with streptavidin-APC (Agilent). The APC decoy reagent was generated by conjugating SA-APC to Dylight 755 using a DyLight 755 antibody labeling kit (ThermoFisher), washing and removing unbound DyLight 755, and incubating with excess of an irrelevant biotinylated His-tagged protein. The PE decoy was generated in the same manner, by conjugating SA-PE to Alexa Fluor 647 with an AF647 antibody labeling kit (ThermoFisher).

### Mouse immunization, cell enrichment, and flow cytometry

For phenotyping of B cells, 6-week old female BALB/c mice, three per dosing group, were immunized intramuscularly with 50 μL per injection site of vaccine formulations containing 5 μg of SARS-CoV-2 antigen (either S-2P trimer or RBD, but not including mass from the I53-50 nanoparticle) mixed 1:1 vol/vol with AddaVax adjuvant on day 0. All experimental mice were euthanized for harvesting of inguinal and popliteal lymph nodes on day 11. The experiment was repeated two times. Popliteal and inguinal lymph nodes were collected and pooled for individual mice. Cell suspensions were prepared by mashing lymph nodes and filtering through 100 μM Nitex mesh. Cells were resuspended in PBS containing 2% FBS and Fc block (2.4G2), and were incubated with 10 nM Decoy tetramers at room temperature for 20 min. RBD-PE tetramer and Spike-APC tetramer were added at a concentration of 10 nM and incubated on ice for 20 min. Cells were washed, incubated with anti-PE and anti-APC magnetic beads on ice for 30 min, then passed over magnetized LS columns (Miltenyi Biotec). Bound B cells were stained with anti-mouse B220 (BUV737), CD3 (PerCP-Cy5.5), CD138 (BV650), CD38 (Alexa Fluor 700), GL7 (eFluor 450), IgM (BV786), IgD (BUV395), CD73 (PE-Cy7), and CD80 (BV605) on ice for 20 min. Cells were run on the Cytek Aurora and analyzed using FlowJo software (Treestar). Cell counts were determined using Accucheck cell counting beads.

### NHP immunization

A Pigtail macaque was immunized with 250 μg of RBD-12GS-I53-50 nanoparticle (88 μg RBD antigen) at day 0 and day 28. Blood was collected at days 0, 10, 14, 28, 42, and 56 days post-prime. Serum and plasma were collected as previously described (Erasmus et al., 2020). Prior to vaccination or blood collection, animals were sedated with an intramuscular injection (10 mg/kg) of ketamine (Ketaset®; Henry Schein). Prior to inoculation, immunogen suspensions were gently mixed 1:1 vol/vol with AddaVax adjuvant (Invivogen, San Diego, CA) to reach a final concentration of 0.250 mg/mL antigen. The vaccine was delivered intramuscularly into both quadriceps muscles with 1 mL per injection site on days 0 and 28. All injection sites were shaved prior to injection and monitored post-injection for any signs of local reactogenicity. At each study timepoint, full physical exams and evaluation of general health were performed on the animals, as previously described (Erasmus et al., 2020), and no adverse events were observed.

### Competition Bio-layer Interferometry

Purification of Fabs from NHP serum was adapted from (Boyoglu-Barnum et al., 2020). Briefly, 1mL of day 56 serum was diluted to 10 mL with PBS and incubated with 1 mL of 3x PBS washed protein A beads (GenScript) with agitation overnight at 37°C. The next day beads were thoroughly washed with PBS using a gravity flow column and bound antibodies were eluted with 0.1 M glycine pH 3.5 into 1M Tris-HCl (pH 8.0) to a final concentration of 100 mM. Serum and early washes that flowed through were re-bound to beads overnight again for a second, repeat elution. IgGs were concentrated (Amicon 30 kDa) and buffer exchanged into PBS. 2x digestion buffer (40 mM sodium phosphate pH 6.5, 20 mM EDTA, 40 mM cysteine) was added to concentrated and pooled IgGs. 500 μL of resuspended immobilized papain resin (ThermoFisher Scientific) freshly washed in 1x digestion buffer (20 mM sodium phosphate, 10 mM EDTA, 20 mM cysteine, pH 6.5) was added to purified IgGs in 2x digestion buffer and samples were agitated for 5 h at 37°C. The supernatant was separated from resin and resin washes were collected and pooled with the resin flow through. Pooled supernatants were sterile-filtered at 0.22 μm and applied 6x to PBS-washed protein A beads in a gravity flow column. The column was eluted as described above and the papain procedure repeated overnight with undigested IgGs to increase yield. The protein A flowthroughs were pooled, concentrated (Amicon 10 kDa), and buffer exchanged into PBS. Purity was checked by SDS-PAGE.

Epitope competition was performed and analyzed using BLI on an Octet Red 96 System (Pall Forté Bio/Sartorius) at 30°C with shaking at 1000 rpm. NTA biosensors (Pall Forté Bio/Sartorius) were hydrated in water for at least 10 minutes, and were then equilibrated in 10x Kinetics buffer (KB) (Pall Forté Bio/Sartorius) for 60 seconds. 10 ng/µL monomeric RBD in 10x KB was loaded for 100 seconds prior to baseline acquisition in 10xKB for 300 seconds. Tips were then dipped into diluted polyclonal Fab in 10x KB in a 1:3 serial dilution beginning with 5000 nM for 2000 seconds or maintained in 10xKB. Tips bound at varying levels depending on the polyclonal Fab concentration. Tips were then dipped into the same concentration of polyclonal Fab plus either 200 nM of hACE2, 400nM CR3022, or 20nM S309 and incubated for 300-2000 seconds. The data were baseline subtracted and aligned to pre-loading with polyclonal Fabs using the Pall Forté Bio/Sartorius analysis software (version 12.0) and plotted in PRISM.

---

**Figure S1.**
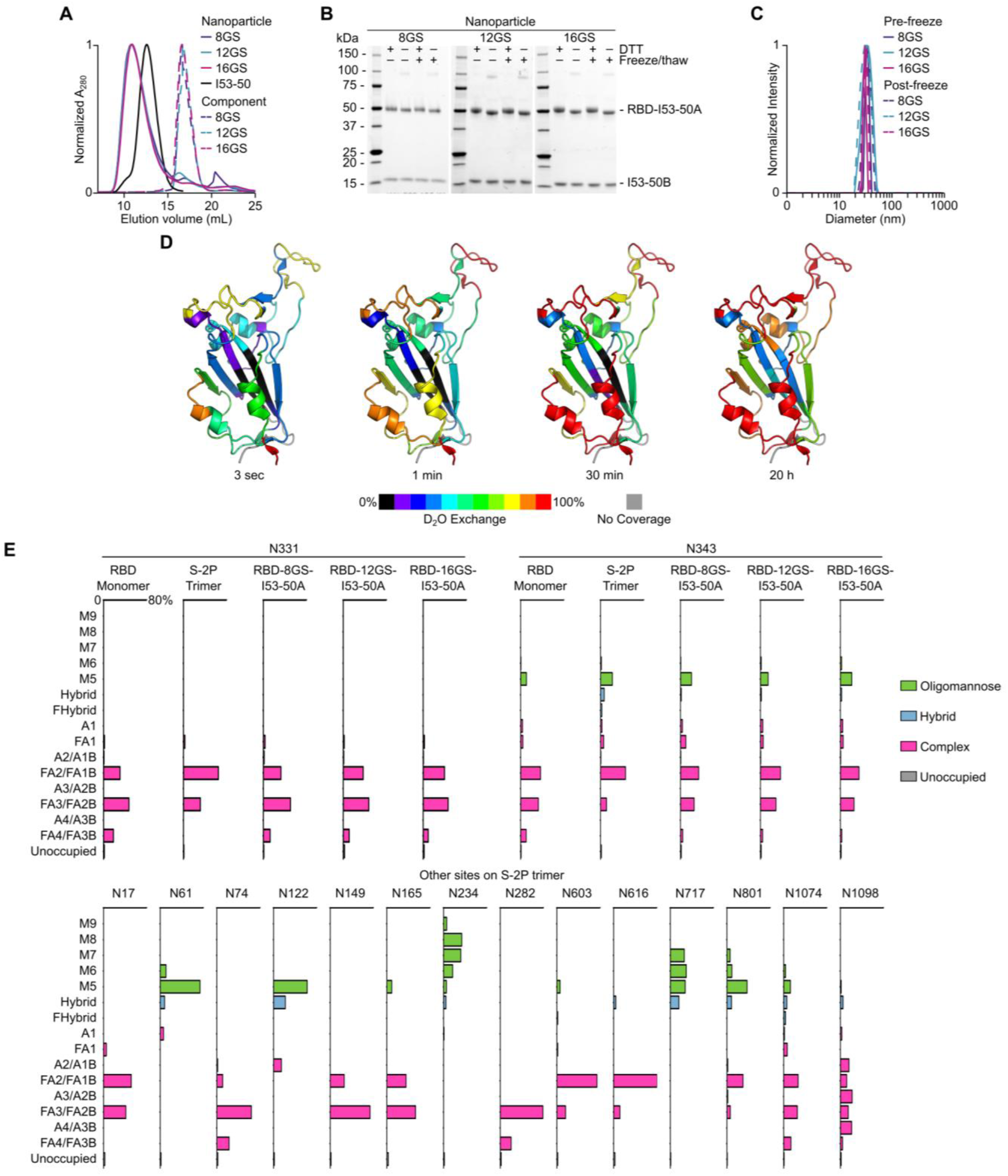
Additional characterization of RBD Nanoparticle Immunogens, Related to Figure 1. (A) Size exclusion chromatography of RBD-I53-50 nanoparticles, unmodified I53-50 nanoparticle, and trimeric RBD-I53-50A components on a Superose 6 Increase 10/300 GL. (B)SDS-PAGE of SEC-purified RBD-I53-50 nanoparticles under reducing and non-reducing conditions before and after one freeze/thaw cycle. (C) Dynamic light scattering of RBD-I53-50 nanoparticles before and after one freeze/thaw cycle indicates monodisperse nanoparticles with a lack of detectable aggregates in each sample. (D) Hydrogen/Deuterium-exchange mass spectrometry, represented here as heatmaps, reveals the structural accessibility and dynamics on RBD (PDB 6W41). Color codes indicate deuterium uptake levels. Monomeric RBD and RBD-8GS-I53-50A have indistinguishable uptake patterns, and are presented in a single heatmap at each time point. (E) Top, bar graphs reveal similar glycan profiles at the N-linked glycosylation sites N331 and N343 in five protein samples: monomeric RBD, S-2P trimer, and RBD-8GS-, RBD-12GS-, and RBD-16GS-I53-50A trimeric components. Bottom, comprehensive glycan profiling on other N-linked glycosylation sites besides N331 and N343 that are found in the S-2P trimer. The axis of each bar graph is scaled to 0-80%. M9 to M5, oligomannose with 9 to 5 mannose residues, are colored green. Hybrid and FHybrid, hybrid types with or without fucosylation, are blue. Subtypes in complex type, shown in pink, are classified based on antennae number and fucosylation.

**Figure S2.**
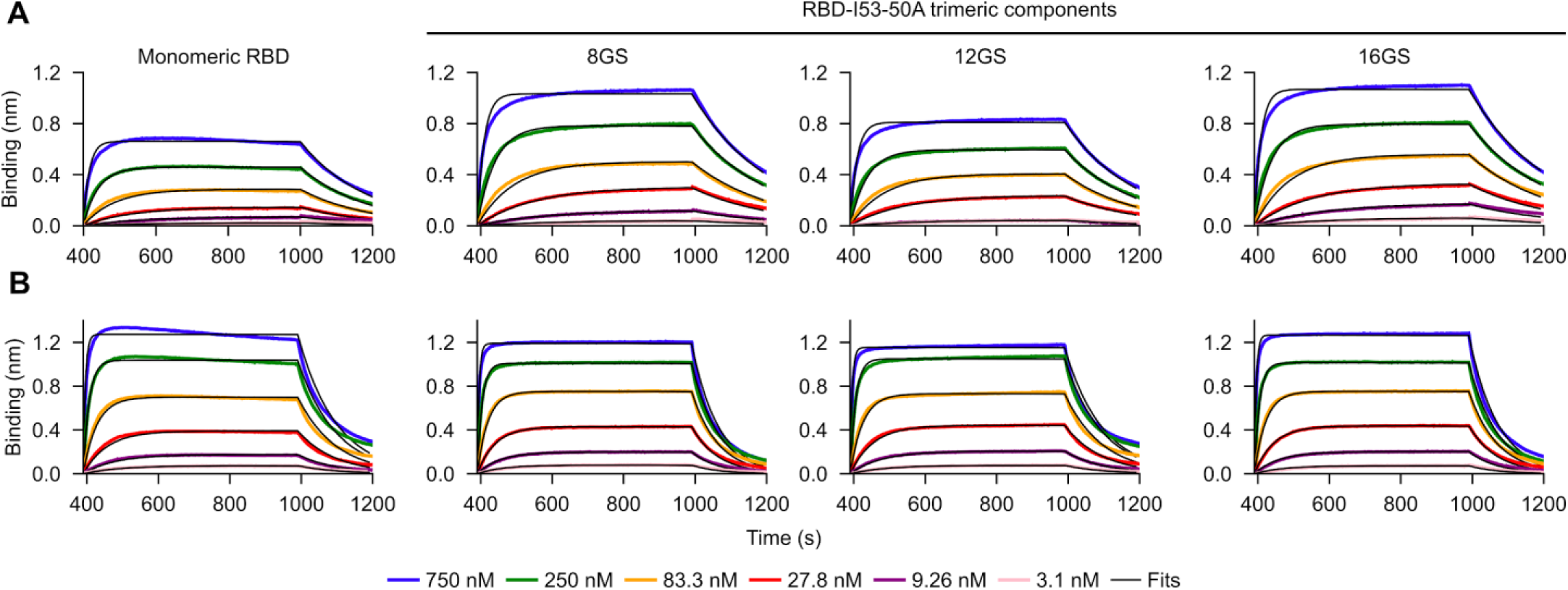
Determination of hACE2 and CR3022 Fab Affinities by Bio-layer Interferometry, Related to Table 1. (A) Analysis of monomeric hACE2 binding to immobilized monomeric RBD and trimeric RBD-8GS-, RBD-12GS-, and RBD-16GS-I53-50A components. (B) Analysis of CR3022 Fab binding to immobilized monomeric RBD and trimeric RBD-8GS-, RBD-12GS-, and RBD-16GS-I53-50A components. Affinity constants (Table 1) were determined by global fitting of the kinetic data from six analyte concentrations to a 1:1 binding model.

**Figure S3.**
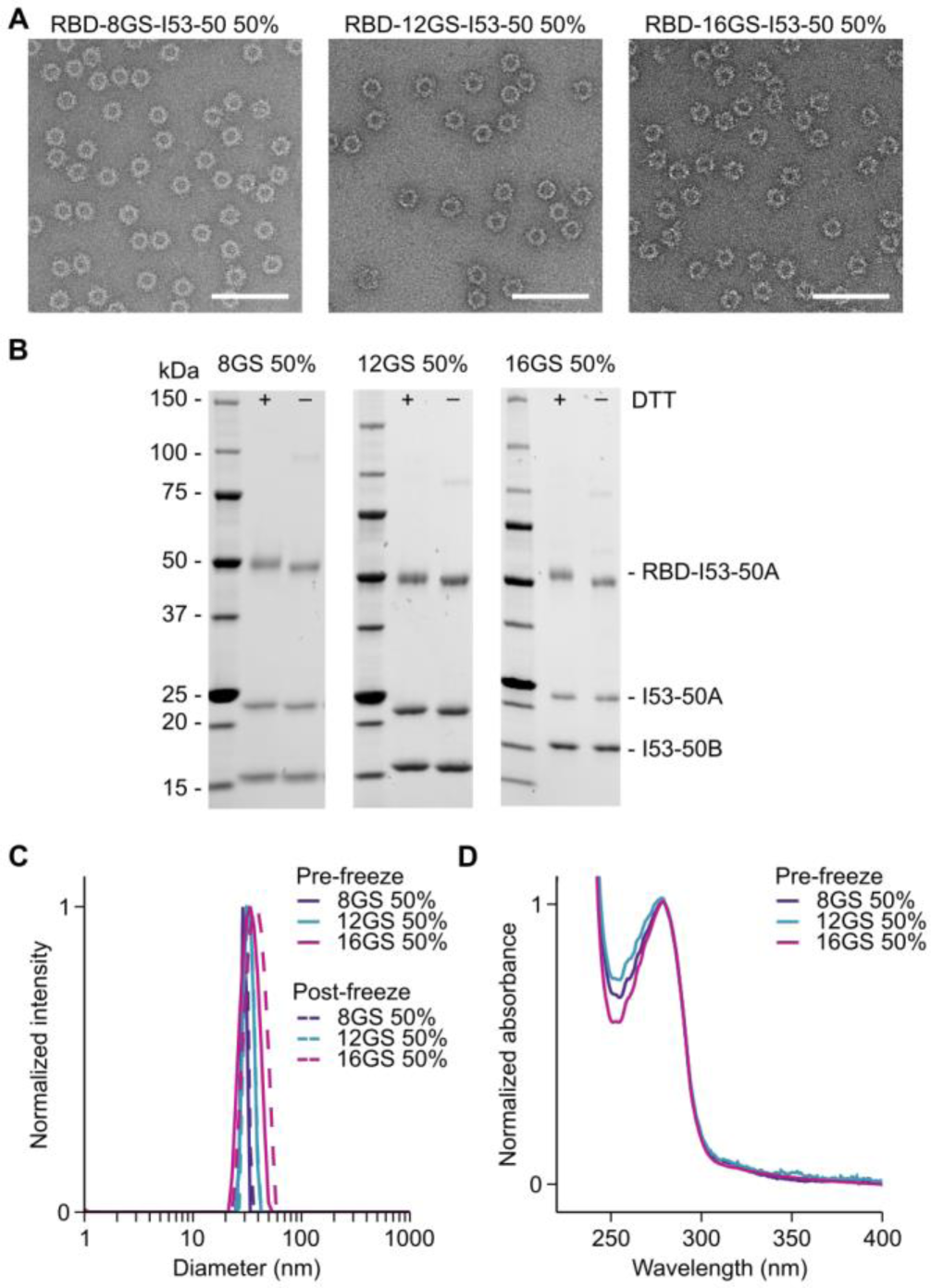
Characterization of Partial Valency RBD Nanoparticles, Related to Figure 2. (A) Representative electron micrographs of negatively stained RBD-8GS-, RBD-12GS-, and RBD-16GS-I53-50 nanoparticles displaying the RBD at 50% valency. The samples were imaged after one freeze/thaw cycle. Scale bars, 100 nm. (B) SDS-PAGE of purified RBD-8GS-, RBD-12GS-, and RBD-16GS-I53-50 nanoparticles displaying the RBD at 50% valency. Both RBD-bearing and unmodified I53-50A subunits are visible on the gels. (C) Dynamic light scattering (DLS) of 50% valency RBD-8GS-, RBD-12GS-, and RBD-16GS-I53-50 nanoparticles both before and after freeze/thaw. No aggregates or unassembled components were observed. (D) UV/vis absorption spectra of 50% valency RBD-8GS-, RBD-12GS-, and RBD-16GS-I53-50 nanoparticles. Turbidity in the samples is low, as indicated by the low absorbance at 320 nm.

**Figure S4.**
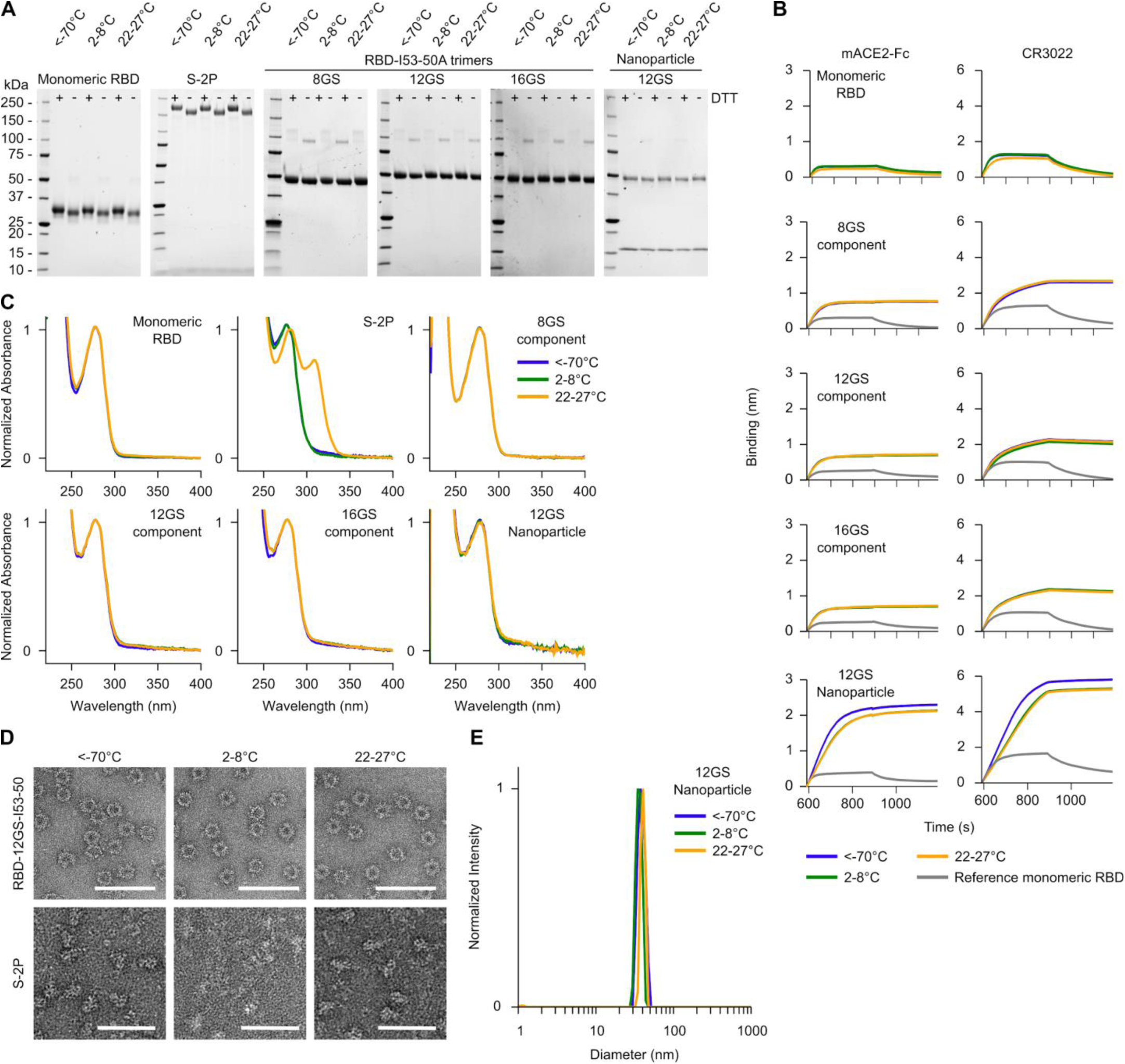
Day 28 Stability Data, Related to Figure 3. (A) SDS-PAGE of purified monomeric RBD, S-2P trimer, RBD-I53-50A components and RBD-12GS-I53-50 nanoparticle in reducing and non-reducing conditions. No degradation of any immunogen was observed after a four-week incubation at any temperature analyzed. (B) Analysis of mACE2-Fc and CR3022 IgG binding to monomeric RBD, RBD-I53-50A trimeric components, and RBD-12GS-I53-50 nanoparticle by BLI after a four-week incubation at three temperatures. Monomeric RBD was used as a reference standard in nanoparticle component and nanoparticle BLI experiments. The RBD-12GS-I53-50 nanoparticle lost minimal binding at the higher temperatures after four weeks; the remaining antigens did not lose any mACE2-Fc or CR3022 IgG binding over the course of the study. (C) UV/vis spectroscopy showed minimal absorbance in the near-UV, suggesting a lack of aggregation/particulates after a four week-incubation at three temperatures, with the exception of S-2P trimer, which gained significant absorbance around 320 nm at ambient temperature. RBD-12GS-I53-50 nanoparticle samples at 22-27°C at several earlier time points exhibited similar peaks near 320 nm (see Supplementary Item 2). (D) nsEM of RBD-12GS-I53-50 nanoparticle (top) and and S-2P trimer (bottom) after a four-week incubation at three temperatures. Intact monodisperse nanoparticles were observed at all temperatures, with no observed degradation or aggregation. The S-2P trimer remained well folded in the <−70 and 22-27°C samples, but was unfolded in samples incubated at 2-8°C. Scale bars: RBD-12GS-I53-50, 100 nm; S-2P, 50 nm. (E) DLS of the RBD-12GS-I53-50 nanoparticle after a four-week incubation at three temperatures. No aggregation was observed at any temperature.

**Figure S5.**
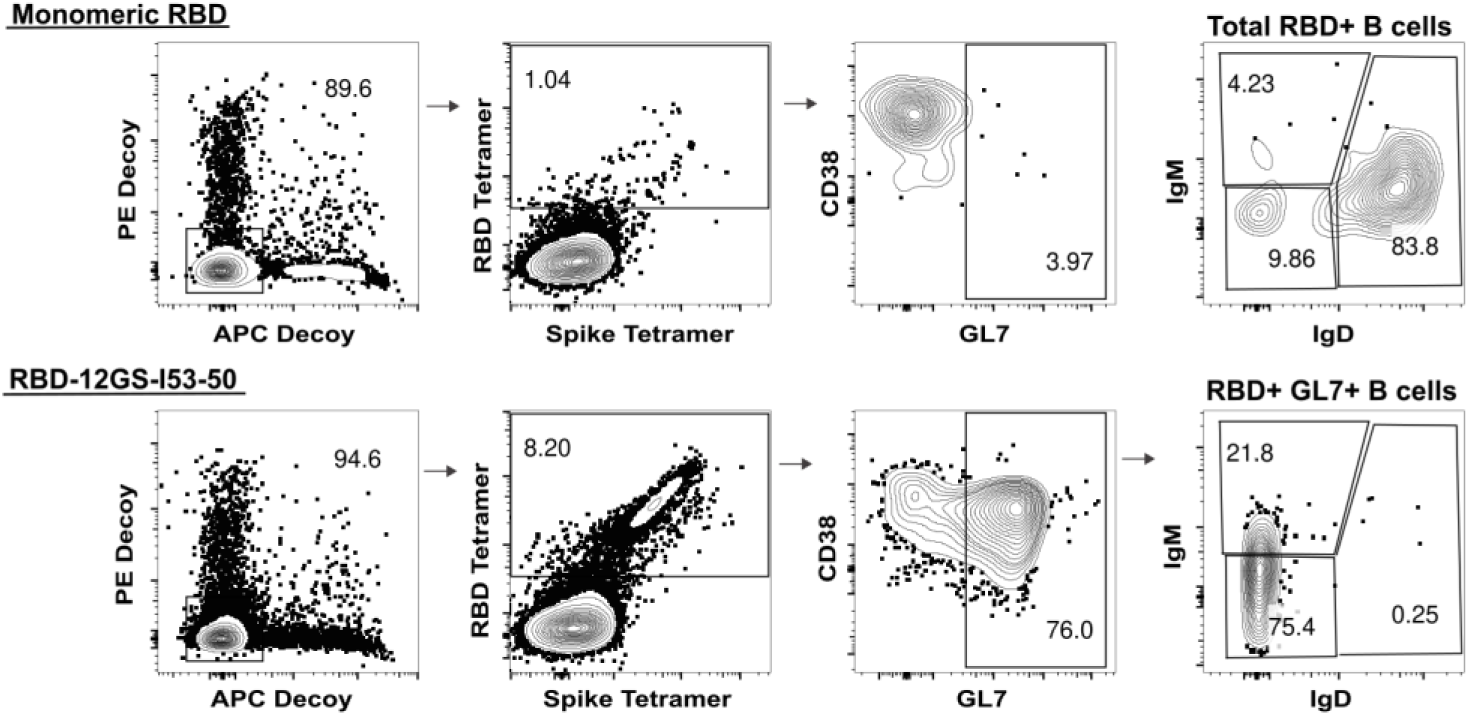
B Cell Gating Strategy, Related to Figure 6. Representative gating strategy for evaluating RBD-specific cells, germinal center (GC) precursors and B cells (CD38^+/−^GL7^+^), and B cell isotypes.

## REFERENCES

Alsoussi, W.B., Turner, J.S., Case, J.B., Zhao, H., Schmitz, A.J., Zhou, J.Q., Chen, R.E., Lei, T., Rizk, A.A., McIntire, K.M., et al.(2020). A Potently Neutralizing Antibody Protects Mice against SARS-CoV-2 Infection. J Immunol.

Anthony, S.J., Gilardi, K., Menachery, V.D., Goldstein, T., Ssebide, B., Mbabazi, R., Navarrete-Macias, I., Liang, E., Wells, H., Hicks, A., et al. (2017). Further Evidence for Bats as the Evolutionary Source of Middle East Respiratory Syndrome Coronavirus. MBio 8.

Anywaine, Z., Whitworth, H., Kaleebu, P., Praygod, G., Shukarev, G., Manno, D., Kapiga, S., Grosskurth, H., Kalluvya, S., Bockstal, V., et al. (2019). Safety and Immunogenicity of a 2-Dose Heterologous Vaccination Regimen With Ad26.ZEBOV and MVA-BN-Filo Ebola Vaccines: 12-Month Data From a Phase 1 Randomized Clinical Trial in Uganda and Tanzania. J Infect Dis 220, 46–56.

Bale, J.B., Gonen, S., Liu, Y., Sheffler, W., Ellis, D., Thomas, C., Cascio, D., Yeates, T.O., Gonen, T., King, N.P., et al. (2016). Accurate design of megadalton-scale two-component icosahedral protein complexes. Science 353, 389–394.

Barnes, C.O., West, A.P., Huey-Tubman, K.E., Hoffmann, M.A.G., Sharaf, N.G., Hoffman, P.R., Koranda, N., Gristick, H.B., Gaebler, C., Muecksch, F., et al. (2020). Structures of Human Antibodies Bound to SARS-CoV-2 Spike Reveal Common Epitopes and Recurrent Features of Antibodies. Cell.

Boyoglu-Barnum, S., Ellis, D., Gillespie, R.A., Hutchinson, G.B., Park, Y.-J., Moin, S.M., Acton, O., Ravichandran, R., Murphy, M., Pettie, D., et al. (2020). Elicitation of broadly protective immunity to influenza by multivalent hemagglutinin nanoparticle vaccines. bioRxiv, 2020.2005.2030.125179.

Brouwer, P.J.M., Antanasijevic, A., Berndsen, Z., Yasmeen, A., Fiala, B., Bijl, T.P.L., Bontjer, I., Bale, J.B., Sheffler, W., Allen, J.D., et al. (2019). Enhancing and shaping the immunogenicity of native-like HIV-1 envelope trimers with a two-component protein nanoparticle. Nat Commun 10, 4272.

Brouwer, P.J.M., Caniels, T.G., van der Straten, K., Snitselaar, J.L., Aldon, Y., Bangaru, S., Torres, J.L., Okba, N.M.A., Claireaux, M., Kerster, G., et al. (2020). Potent neutralizing antibodies from COVID-19 patients define multiple targets of vulnerability. Science.

Bruun, T.U.J., Andersson, A.C., Draper, S.J., and Howarth, M. (2018). Engineering a Rugged Nanoscaffold To Enhance Plug-and-Display Vaccination. ACS Nano 12, 8855–8866.

Corbett, K.S., Edwards, D.K., Leist, S.R., Abiona, O.M., Boyoglu-Barnum, S., Gillespie, R.A., Himansu, S., Schäfer, A., Ziwawo, C.T., DiPiazza, A.T., et al. (2020). SARS-CoV-2 mRNA vaccine design enabled by prototype pathogen preparedness. Nature.

Corti, D., Zhao, J., Pedotti, M., Simonelli, L., Agnihothram, S., Fett, C., Fernandez-Rodriguez, B., Foglierini, M., Agatic, G., Vanzetta, F., et al. (2015). Prophylactic and postexposure efficacy of a potent human monoclonal antibody against MERS coronavirus. Proc Natl Acad Sci U S A 112, 10473–10478.

Dai, L., Zheng, T., Xu, K., Han, Y., Xu, L., Huang, E., An, Y., Cheng, Y., Li, S., Liu, M., et al. (2020). A Universal Design of Betacoronavirus Vaccines against COVID-19, MERS, and SARS. Cell.

Davis, A.K.F., McCormick, K., Gumina, M.E., Petrie, J.G., Martin, E.T., Xue, K.S., Bloom, J.D., Monto, A.S., Bushman, F.D., and Hensley, S.E. (2018). Sera from Individuals with Narrowly Focused Influenza Virus Antibodies Rapidly Select Viral Escape Mutations. J Virol 92.

Dinnon, K.H., Leist, S.R., Schäfer, A., Edwards, C.E., Martinez, D.R., Montgomery, S.A., West, A., Yount, B.L., Hou, Y.J., Adams, L.E., et al. (2020). A mouse-adapted SARS-CoV-2 model for the evaluation of COVID-19 medical countermeasures. bioRxiv, 2020.2005.2006.081497.

Edwards, R.J., Mansouri, K., Stalls, V., Manne, K., Watts, B., Parks, R., Gobeil, S.M.C., Janowska, K., Li, D., Lu, X., et al. (2020). Cold sensitivity of the SARS-CoV-2 spike ectodomain. bioRxiv.

Erasmus, J.H., Khandhar, A.P., O'Connor, M.A., Walls, A.C., Hemann, E.A., Murapa, P., Archer, J., Leventhal, S., Fuller, J.T., Lewis, T.B., et al. (2020). An Alphavirus-derived replicon RNA va*ccine indu*ces SARS-CoV-2 neutralizing antibody and T cell responses in mice and nonhuman primates. Sci Transl Med 12.

Folegatti, P.M., Ewer, K.J., Aley, P.K., Angus, B., Becker, S., Belij-Rammerstorfer, S., Bellamy, D., Bibi, S., Bittaye, M., Clutterbuck, E.A., et al. (2020). Safety and immunogenicity of the ChAdOx1 nCoV-19 vaccine against SARS-CoV-2: a preliminary report of a phase 1/2, single-blind, randomised controlled trial. Lancet.

Gibson, D.G., Young, L., Chuang, R.Y., Venter, J.C., Hutchison, C.A., and Smith, H.O. (2009). Enzymatic assembly of DNA molecules up to several hundred kilobases. Nat Methods 6, 343–345.

Goddard, T.D., Huang, C.C., Meng, E.C., Pettersen, E.F., Couch, G.S., Morris, J.H., and Ferrin, T.E. (2018). UCSF ChimeraX: Meeting modern challenges in visualization and analysis. Protein Sci 27, 14–25.

Graham, B.S. (2020). Rapid COVID-19 vaccine development. Science 368, 945–946.

Guttman, M., Weis, D.D., Engen, J.R., and Lee, K.K. (2013). Analysis of overlapped and noisy hydrogen/deuterium exchange mass spectra. J Am Soc Mass Spectrom 24, 1906–1912.

Henderson, R., Edwards, R.J., Mansouri, K., Janowska, K., Stalls, V., Gobeil, S.M.C., Kopp, M., Li, D., Parks, R., Hsu, A.L., et al. (2020). Controlling the SARS-CoV-2 spike glycoprotein conformation. Nat Struct Mol Biol.

Hoffmann, M., Kleine-Weber, H., Schroeder, S., Krüger, N., Herrler, T., Erichsen, S., Schiergens, T.S., Herrler, G., Wu, N.H., Nitsche, A., et al. (2020). SARS-CoV-2 Cell Entry Depends on ACE2 and TMPRSS2 and Is Blocked by a Clinically Proven Protease Inhibitor. Cell 181, 271–280.e278.

Hou, Y.J., Okuda, K., Edwards, C.E., Martinez, D.R., Asakura, T., Dinnon, K.H., Kato, T., Lee, R.E., Yount, B.L., Mascenik, T.M., et al. (2020). SARS-CoV-2 Reverse Genetics Reveals a Variable Infection Gradient in the Respiratory Tract. Cell 182, 429–446.e414.

Hsia, Y., Bale, J.B., Gonen, S., Shi, D., Sheffler, W., Fong, K.K., Nattermann, U., Xu, C., Huang, P.S., Ravichandran, R., et al. (2016). Design of a hyperstable 60-subunit protein dodecahedron. [corrected]. Nature 535, 136–139.

Hsieh, C.L., Goldsmith, J.A., Schaub, J.M., DiVenere, A.M., Kuo, H.C., Javanmardi, K., Le, K.C., Wrapp, D., Lee, A.G., Liu, Y., et al. (2020). Structure-based design of prefusion-stabilized SARS-CoV-2 spikes. Science.

Huo, J., Zhao, Y., Ren, J., Zhou, D., Duyvesteyn, H.M.E., Ginn, H.M., Carrique, L., Malinauskas, T., Ruza, R.R., Shah, P.N.M., et al. (2020). Neutralisation of SARS-CoV-2 by destruction of the prefusion Spike. Cell Host & Microbe.

Irvine, D.J., and Read, B.J. (2020). Shaping humoral immunity to vaccines through antigen-displaying nanoparticles. Curr Opin Immunol 65, 1–6.

Jackson, L.A., Anderson, E.J., Rouphael, N.G., Roberts, P.C., Makhene, M., Coler, R.N., McCullough, M.P., Chappell, J.D., Denison, M.R., Stevens, L.J., et al. (2020). An mRNA Vaccine against SARS-CoV-2 - Preliminary Report. N Engl J Med.

Kanekiyo, M., Bu, W., Joyce, M.G., Meng, G., Whittle, J.R., Baxa, U., Yamamoto, T., Narpala, S., Todd, J.P., Rao, S.S., et al. (2015). Rational Design of an Epstein-Barr Virus Vaccine Targeting the Receptor-Binding Site. Cell 162, 1090–1100.

Kanekiyo, M., Ellis, D., and King, N.P. (2019a). New Vaccine Design and Delivery Technologies. J Infect Dis 219, S88–S96.

Kanekiyo, M., and Graham, B.S. (2020). Next-Generation Influenza Vaccines. Cold Spring Harb Perspect Med.

Kanekiyo, M., Joyce, M.G., Gillespie, R.A., Gallagher, J.R., Andrews, S.F., Yassine, H.M., Wheatley, A.K., Fisher, B.E., Ambrozak, D.R., Creanga, A., et al. (2019b). Mosaic nanoparticle display of diverse influenza virus hemagglutinins elicits broad B cell responses. Nat Immunol 20, 362–372.

Kanekiyo, M., Wei, C.J., Yassine, H.M., McTamney, P.M., Boyington, J.C., Whittle, J.R., Rao, S.S., Kong, W.P., Wang, L., and Nabel, G.J. (2013). Self-assembling influenza nanoparticle vaccines elicit broadly neutralizing H1N1 antibodies. Nature 499, 102–106.

Keech, C., Albert, G., Reed, P., Neal, S., Plested, J.S., Zhu, M., Cloney-Clark, S., Zhou, H., Patel, N., Frieman, M.B., et al. (2020). First-in-Human Trial of a SARS CoV 2 Recombinant Spike Protein Nanoparticle Vaccine. bioRxiv, 2020.08.05.20168435.

Kim, H.W., Canchola, J.G., Brandt, C.D., Pyles, G., Chanock, R.M., Jensen, K., and Parrott, R.H. (1969). Respiratory syncytial virus disease in infants despite prior administration of antigenic inactivated vaccine. Am J Epidemiol 89, 422–434.

King, N.P., Sheffler, W., Sawaya, M.R., Vollmar, B.S., Sumida, J.P., Andre, I., Gonen, T., Yeates, T.O., and Baker, D. (2012). Computational design of self-assembling protein nanomaterials with atomic level accuracy. Science 336, 1171–1174.

Kirchdoerfer, R.N., Wang, N., Pallesen, J., Wrapp, D., Turner, H.L., Cottrell, C.A., Corbett, K.S., Graham, B.S., McLellan, J.S., and Ward, A.B. (2018). Stabilized coronavirus spikes are resistant to conformational changes induced by receptor recognition or proteolysis. Sci Rep 8, 15701.

Kreimer, A.R., Herrero, R., Sampson, J.N., Porras, C., Lowy, D.R., Schiller, J.T., Schiffman, M., Rodriguez, A.C., Chanock, S., Jimenez, S., et al. (2018). Evidence for single-dose protection by the bivalent HPV vaccine-Review of the Costa Rica HPV vaccine trial and future research studies. Vaccine 36, 4774–4782.

Krenkova, J., Szekrenyes, A., Keresztessy, Z., Foret, F., and Guttman, A. (2013). Oriented immobilization of peptide-N-glycosidase F on a monolithic support for glycosylation analysis. J Chromatogr A 1322, 54–61.

Krishnamurty, A.T., Thouvenel, C.D., Portugal, S., Keitany, G.J., Kim, K.S., Holder, A., Crompton, P.D., Rawlings, D.J., and Pepper, M. (2016). Somatically Hypermutated Plasmodium-Specific IgM(+) Memory B Cells Are Rapid, Plastic, Early Responders upon Malaria Rechallenge. Immunity 45, 402–414.

Kumru, O.S., Joshi, S.B., Smith, D.E., Middaugh, C.R., Prusik, T., and Volkin, D.B. (2014). Vaccine instability in the cold chain: mechanisms, analysis and formulation strategies. Biologicals 42, 237–259.

Lan, J., Ge, J., Yu, J., Shan, S., Zhou, H., Fan, S., Zhang, Q., Shi, X., Wang, Q., Zhang, L., et al. (2020). Structure of the SARS-CoV-2 spike receptor-binding domain bound to the ACE2 receptor. Nature.

Lee, E.C., Liang, Q., Ali, H., Bayliss, L., Beasley, A., Bloomfield-Gerdes, T., Bonoli, L., Brown, R., Campbell, J., Carpenter, A., et al. (2014). Complete humanization of the mouse immunoglobulin loci enables efficient therapeutic antibody discovery. Nat Biotechnol 32, 356–363.

Lee, J.M., Eguia, R., Zost, S.J., Choudhary, S., Wilson, P.C., Bedford, T., Stevens-Ayers, T., Boeckh, M., Hurt, A.C., Lakdawala, S.S., et al. (2019). Mapping person-to-person variation in viral mutations that escape polyclonal serum targeting influenza hemagglutinin. Elife 8.

Letko, M., Marzi, A., and Munster, V. (2020). Functional assessment of cell entry and receptor usage for SARS-CoV-2 and other lineage B betacoronaviruses. Nature Microbiology.

Li, W., Moore, M.J., Vasilieva, N., Sui, J., Wong, S.K., Berne, M.A., Somasundaran, M., Sullivan, J.L., Luzuriaga, K., Greenough, T.C., et al. (2003). Angiotensin-converting enzyme 2 is a functional receptor for the SARS coronavirus. Nature 426, 450–454.

Li, X., Wang, W., Zhao, X., Zai, J., Zhao, Q., Li, Y., and Chaillon, A. (2020). Transmission dynamics and evolutionary history of 2019-nCoV. J Med Virol 92, 501–511.

Lopez-Sagaseta, J., Malito, E., Rappuoli, R., and Bottomley, M.J. (2016). Self-assembling protein nanoparticles in the design of vaccines. Comput Struct Biotechnol J 14, 58–68.

Mandolesi, M., Sheward, D.J., Hanke, L., Ma, J., Pushparaj, P., Vidakovics, L.P., Kim, C., Loré, K., Dopico, X.C., Coquet, J.M., et al. (2020). SARS-CoV-2 protein subunit vaccination elicits potent neutralizing antibody responses. bioRxiv, 2020.2007.2031.228486.

Marcandalli, J., Fiala, B., Ols, S., Perotti, M., de van der Schueren, W., Snijder, J., Hodge, E., Benhaim, M., Ravichandran, R., Carter, L., et al. (2019). Induction of Potent Neutralizing Antibody Responses by a Designed Protein Nanoparticle Vaccine for Respiratory Syncytial Virus. Cell 176, 1420–1431 e1417.

McCallum, M., Walls, A.C., Bowen, J.E., Corti, D., and Veesler, D. (2020). Structure-guided covalent stabilization of coronavirus spike glycoprotein trimers in the closed conformation. Nat Struct Mol Biol.

Menachery, V.D., Yount, B.L., Jr., Debbink, K., Agnihothram, S., Gralinski, L.E., Plante, J.A., Graham, R.L., Scobey, T., Ge, X.Y., Donaldson, E.F., et al. (2015). A SARS-like cluster of circulating bat coronaviruses shows potential for human emergence. Nat Med 21, 1508–1513.

Menachery, V.D., Yount, B.L., Jr., Sims, A.C., Debbink, K., Agnihothram, S.S., Gralinski, L.E., Graham, R.L., Scobey, T., Plante, J.A., Royal, S.R., et al. (2016). SARS-like WIV1-CoV poised for human emergence. Proc Natl Acad Sci U S A 113, 3048–3053.

Millet, J.K., and Whittaker, G.R. (2016). Murine Leukemia Virus (MLV)-based Coronavirus Spike-pseudotyped Particle Production and Infection. Bio Protoc 6.

Mulligan, M.J., Lyke, K.E., Kitchin, N., Absalon, J., Gurtman, A., Lockhart, S.P., Neuzil, K., Raabe, V., Bailey, R., Swanson, K.A., et al. (2020). Phase 1/2 Study to Describe the Safety and Immunogenicity of a COVID-19 RNA Vaccine Candidate (BNT162b1) in Adults 18 to 55 Years of Age: Interim Report. medRxiv, 2020.2006.2030.20142570.

Pallesen, J., Wang, N., Corbett, K.S., Wrapp, D., Kirchdoerfer, R.N., Turner, H.L., Cottrell, C.A., Becker, M.M., Wang, L., Shi, W., et al. (2017). Immunogenicity and structures of a rationally designed prefusion MERS-CoV spike antigen. Proc Natl Acad Sci U S A 114, E7348–E7357.

Pinto, D., Park, Y.J., Beltramello, M., Walls, A.C., Tortorici, M.A., Bianchi, S., Jaconi, S., Culap, K., Zatta, F., De Marco, A., et al. (2020). Cross-neutralization of SARS-CoV-2 by a human monoclonal SARS-CoV antibody. Nature 583, 290–295.

Poh, C.M., Carissimo, G., Wang, B., Amrun, S.N., Lee, C.Y., Chee, R.S., Fong, S.W., Yeo, N.K., Lee, W.H., Torres-Ruesta, A., et al. (2020). Two linear epitopes on the SARS-CoV-2 spike protein that elicit neutralising antibodies in COVID-19 patients. Nat Commun 11, 2806.

Polack, F.P., Teng, M.N., Collins, P.L., Prince, G.A., Exner, M., Regele, H., Lirman, D.D., Rabold, R., Hoffman, S.J., Karp, C.L., et al. (2002). A role for immune complexes in enhanced respiratory syncytial virus disease. J Exp Med 196, 859–865.

Robbiani, D.F., Gaebler, C., Muecksch, F., Lorenzi, J.C.C., Wang, Z., Cho, A., Agudelo, M., Barnes, C.O., Gazumyan, A., Finkin, S., et al. (2020). Convergent antibody responses to SARS-CoV-2 in convalescent individuals. Nature.

Rockx, B., Corti, D., Donaldson, E., Sheahan, T., Stadler, K., Lanzavecchia, A., and Baric, R. (2008). Structural basis for potent cross-neutralizing human monoclonal antibody protection against lethal human and zoonotic severe acute respiratory syndrome coronavirus challenge. J Virol 82, 3220–3235.

Rossen, J.W., de Beer, R., Godeke, G.J., Raamsman, M.J., Horzinek, M.C., Vennema, H., and Rottier, P.J. (1998). The viral spike protein is not involved in the polarized sorting of coronaviruses in epithelial cells. J Virol 72, 497–503.

Sahin, U., Muik, A., Derhovanessian, E., Vogler, I., Kranz, L.M., Vormehr, M., Baum, A., Pascal, K., Quandt, J., Maurus, D., et al. (2020). Concurrent human antibody and T<sub>H</sub>1 type T-cell responses elicited by a COVID-19 RNA vaccine. medRxiv, 2020.2007.2017.20140533.

Seydoux, E., Homad, L.J., MacCamy, A.J., Parks, K.R., Hurlburt, N.K., Jennewein, M.F., Akins, N.R., Stuart, A.B., Wan, Y.-H., Feng, J., et al. (2020). Characterization of neutralizing antibodies from a SARS-CoV-2 infected individual. bioRxiv, 2020.2005.2012.091298.

Shang, J., Ye, G., Shi, K., Wan, Y., Luo, C., Aihara, H., Geng, Q., Auerbach, A., and Li, F. (2020). Structural basis of receptor recognition by SARS-CoV-2. Nature.

Smith, E.C., Sexton, N.R., and Denison, M.R. (2014). Thinking Outside the Triangle: Replication Fidelity of the Largest RNA Viruses. Annu Rev Virol 1, 111–132.

Stettler, K., Beltramello, M., Espinosa, D.A., Graham, V., Cassotta, A., Bianchi, S., Vanzetta, F., Minola, A., Jaconi, S., Mele, F., et al. (2016). Specificity, cross-reactivity, and function of antibodies elicited by Zika virus infection. Science 353, 823–826.

Taylor, J.J., Martinez, R.J., Titcombe, P.J., Barsness, L.O., Thomas, S.R., Zhang, N., Katzman, S.D., Jenkins, M.K., and Mueller, D.L. (2012). Deletion and anergy of polyclonal B cells specific for ubiquitous membrane-bound self-antigen. J Exp Med 209, 2065–2077.

ter Meulen, J., van den Brink, E.N., Poon, L.L., Marissen, W.E., Leung, C.S., Cox, F., Cheung, C.Y., Bakker, A.Q., Bogaards, J.A., van Deventer, E., et al. (2006). Human monoclonal antibody combination against SARS coronavirus: synergy and coverage of escape mutants. PLoS Med 3, e237.

Tortorici, M.A., and Veesler, D. (2019). Structural insights into coronavirus entry. Adv Virus Res 105, 93–116.

Traggiai, E., Becker, S., Subbarao, K., Kolesnikova, L., Uematsu, Y., Gismondo, M.R., Murphy, B.R., Rappuoli, R., and Lanzavecchia, A. (2004). An efficient method to make human monoclonal antibodies from memory B cells: potent neutralization of SARS coronavirus. Nat Med 10, 871–875.

Ueda, G., Antanasijevic, A., Fallas, J.A., Sheffler, W., Copps, J., Ellis, D., Hutchinson, G.B., Moyer, A., Yasmeen, A., Tsybovsky, Y., et al. (2020). Tailored design of protein nanoparticle scaffolds for multivalent presentation of viral glycoprotein antigens. Elife 9.

Verkerke, H.P., Williams, J.A., Guttman, M., Simonich, C.A., Liang, Y., Filipavicius, M., Hu, S.L., Overbaugh, J., and Lee, K.K. (2016). Epitope-Independent Purification of Native-Like Envelope Trimers from Diverse HIV-1 Isolates. J Virol 90, 9471–9482.

Walls, A.C., Park, Y.J., Tortorici, M.A., Wall, A., McGuire, A.T., and Veesler, D. (2020). Structure, Function, and Antigenicity of the SARS-CoV-2 Spike Glycoprotein. Cell 181, 281–292.e286.

Walls, A.C., Tortorici, M.A., Bosch, B.J., Frenz, B., Rottier, P.J.M., DiMaio, F., Rey, F.A., and Veesler, D. (2016a). Cryo-electron microscopy structure of a coronavirus spike glycoprotein trimer. Nature 531, 114–117.

Walls, A.C., Tortorici, M.A., Frenz, B., Snijder, J., Li, W., Rey, F.A., DiMaio, F., Bosch, B.J., and Veesler, D. (2016b). Glycan shield and epitope masking of a coronavirus spike protein observed by cryo-electron microscopy. Nat Struct Mol Biol 23, 899–905.

Walls, A.C., Tortorici, M.A., Snijder, J., Xiong, X., Bosch, B.J., Rey, F.A., and Veesler, D. (2017). Tectonic conformational changes of a coronavirus spike glycoprotein promote membrane fusion. Proc Natl Acad Sci U S A 114, 11157–11162.

Walls, A.C., Xiong, X., Park, Y.J., Tortorici, M.A., Snijder, J., Quispe, J., Cameroni, E., Gopal, R., Dai, M., Lanzavecchia, A., et al. (2019). Unexpected Receptor Functional Mimicry Elucidates Activation of Coronavirus Fusion. Cell 176, 1026–1039.e1015.

Wang, C., Li, W., Drabek, D., Okba, N.M.A., van Haperen, R., Osterhaus, A.D.M.E., van Kuppeveld, F.J.M., Haagmans, B.L., Grosveld, F., and Bosch, B.-J. (2020a). A human monoclonal antibody blocking SARS-CoV-2 infection. bioRxiv, 2020.2003.2011.987958.

Wang, Q., Zhang, Y., Wu, L., Niu, S., Song, C., Zhang, Z., Lu, G., Qiao, C., Hu, Y., Yuen, K.Y., et al. (2020b). Structural and Functional Basis of SARS-CoV-2 Entry by Using Human ACE2. Cell 181, 894–904.e899.

Watanabe, Y., Allen, J.D., Wrapp, D., McLellan, J.S., and Crispin, M. (2020). Site-specific glycan analysis of the SARS-CoV-2 spike. Science.

Weis, D.D., Engen, J.R., and Kass, I.J. (2006). Semi-automated data processing of hydrogen exchange mass spectra using HX-Express. J Am Soc Mass Spectrom 17, 1700–1703.

Woo, P.C., Lau, S.K., Li, K.S., Poon, R.W., Wong, B.H., Tsoi, H.W., Yip, B.C., Huang, Y., Chan, K.H., and Yuen, K.Y. (2006). Molecular diversity of coronaviruses in bats. Virology 351, 180–187.

Wrapp, D., Wang, N., Corbett, K.S., Goldsmith, J.A., Hsieh, C.L., Abiona, O., Graham, B.S., and McLellan, J.S. (2020). Cryo-EM structure of the 2019-nCoV spike in the prefusion conformation. Science 367, 1260–1263.

Wu, Y., Wang, F., Shen, C., Peng, W., Li, D., Zhao, C., Li, Z., Li, S., Bi, Y., Yang, Y., et al. (2020). A noncompeting pair of human neutralizing antibodies block COVID-19 virus binding to its receptor ACE2. Science 368, 1274–1278.

Xiong, X., Qu, K., Ciazynska, K.A., Hosmillo, M., Carter, A.P., Ebrahimi, S., Ke, Z., Scheres, S.H.W., Bergamaschi, L., Grice, G.L., et al. (2020). A thermostable, closed SARS-CoV-2 spike protein trimer. Nat Struct Mol Biol.

Xiong, X., Tortorici, M.A., Snijder, J., Yoshioka, C., Walls, A.C., Li, W., McGuire, A.T., Rey, F.A., Bosch, B.J., and Veesler, D. (2018). Glycan Shield and Fusion Activation of a Deltacoronavirus Spike Glycoprotein Fine-Tuned for Enteric Infections. J Virol 92.

Yan, R., Zhang, Y., Li, Y., Xia, L., Guo, Y., and Zhou, Q. (2020). Structural basis for the recognition of SARS-CoV-2 by full-length human ACE2. Science 367, 1444–1448.

Yang, Y., Liu, C., Du, L., Jiang, S., Shi, Z., Baric, R.S., and Li, F. (2015). Two Mutations Were Critical for Bat-to-Human Transmission of Middle East Respiratory Syndrome Coronavirus. J Virol 89, 9119–9123.

Yu, J., Tostanoski, L.H., Peter, L., Mercado, N.B., McMahan, K., Mahrokhian, S.H., Nkolola, J.P., Liu, J., Li, Z., Chandrashekar, A., et al. (2020). DNA vaccine protection against SARS-CoV-2 in rhesus macaques. Science.

Yuan, M., Wu, N.C., Zhu, X., Lee, C.D., So, R.T.Y., Lv, H., Mok, C.K.P., and Wilson, I.A. (2020). A highly conserved cryptic epitope in the receptor-binding domains of SARS-CoV-2 and SARS-CoV. Science.

Zhang, Z., Zhang, A., and Xiao, G. (2012). Improved protein hydrogen/deuterium exchange mass spectrometry platform with fully automated data processing. Anal Chem 84, 4942–4949.

Zhou, D., Duyvesteyn, H.M.E., Chen, C.P., Huang, C.G., Chen, T.H., Shih, S.R., Lin, Y.C., Cheng, C.Y., Cheng, S.H., Huang, Y.C., et al. (2020a). Structural basis for the neutralization of SARS-CoV-2 by an antibody from a convalescent patient. Nat Struct Mol Biol.

Zhou, H., Chen, X., Hu, T., Li, J., Song, H., Liu, Y., Wang, P., Liu, D., Yang, J., Holmes, E.C., et al. (2020b). A Novel Bat Coronavirus Closely Related to SARS-CoV-2 Contains Natural Insertions at the S1/S2 Cleavage Site of the Spike Protein. Curr Biol 30, 2196–2203.e2193.

Zhou, P., Yang, X.L., Wang, X.G., Hu, B., Zhang, L., Zhang, W., Si, H.R., Zhu, Y., Li, B., Huang, C.L., et al. (2020c). A pneumonia outbreak associated with a new coronavirus of probable bat origin. Nature.

Zhu, F.C., Li, Y.H., Guan, X.H., Hou, L.H., Wang, W.J., Li, J.X., Wu, S.P., Wang, B.S., Wang, Z., Wang, L., et al. (2020a). Safety, tolerability, and immunogenicity of a recombinant adenovirus type-5 vectored COVID-19 vaccine: a dose-escalation, open-label, non-randomised, first-in-human trial. Lancet 395, 1845–1854.

Zhu, N., Zhang, D., Wang, W., Li, X., Yang, B., Song, J., Zhao, X., Huang, B., Shi, W., Lu, R., et al. (2020b). A Novel Coronavirus from Patients with Pneumonia in China,2019. N Engl J Med.

Zost, S.J., Gilchuk, P., Case, J.B., Binshtein, E., Chen, R.E., Nkolola, J.P., Schäfer, A., Reidy, J.X., Trivette, A., Nargi, R.S., et al. (2020). Potently neutralizing and protective human antibodies against SARS-CoV-2. Nature.

