## Supplementary Item 1 for "Elicitation of potent neutralizing antibody responses by designed protein nanoparticle vaccines for SARS-CoV-2"

**Table S1. Amino acid sequences of proteins used in this work, related to Figures 1-6**

>RBD-8GS-I53-50A
MGILPSPGMPALLSLVSLLSVLLMGCVAETGTRFPNITNLCPFGEVFNATRFASVYAWNRKRISNCVADYS
VLYNSASFSTFKCYGVSPTKLNDLCFTNVYADSFVIRGDEVRQIAPGQTGKIADYNYKLPDDFTGCVIAWN
SNNLDSKVGGNYNYLYRLFRKSNLKPFERDISTEIYQAGSTPCNGVEGFNCYFPLQSYGFQPTNGVGYQ
PYRVVVLSFELLHAPATVCGPKKSTGGSGGSGSEKAAKAEEAARKMEELFKKHKIVAVLRANSVEEAIEK
AVAVFAGGVHLIEITFTVPDADTVIKALSVLKEKGAIIGAGTVTSVEQARKAVESGAEFIVSPHLDEEISQFA
KEKGVFYMPGVMTPTELVKAMKLGHTILKLFPGEVVGPQFVKAMKGPFPNVKFVPTGGVNLDNVAEWFK
AGVLAVGVGSALVKGTPDEVREKAKAFVEKIRGATEGGSHHHHHHHH

>RBD-12GS-I53-50A
MGILPSPGMPALLSLVSLLSVLLMGCVAETGTRFPNITNLCPFGEVFNATRFASVYAWNRKRISNCVADYS
VLYNSASFSTFKCYGVSPTKLNDLCFTNVYADSFVIRGDEVRQIAPGQTGKIADYNYKLPDDFTGCVIAWN
SNNLDSKVGGNYNYLYRLFRKSNLKPFERDISTEIYQAGSTPCNGVEGFNCYFPLQSYGFQPTNGVGYQ
PYRVVVLSFELLHAPATVCGPKKSTGSGSGGSGGSGSEKAAKAEEAARKMEELFKKHKIVAVLRANSVE
EAIEKAVAVFAGGVHLIEITFTVPDADTVIKALSVLKEKGAIIGAGTVTSVEQARKAVESGAEFIVSPHLDEEI
SQFAKEKGVFYMPGVMTPTELVKAMKLGHTILKLFPGEVVGPQFVKAMKGPFPNVKFVPTGGVNLDNVA
EWFKAGVLAVGVGSALVKGTPDEVREKAKAFVEKIRGATEGGSHHHHHHHH

>RBD-16GS-I53-50A
MGILPSPGMPALLSLVSLLSVLLMGCVAETGTRFPNITNLCPFGEVFNATRFASVYAWNRKRISNCVADYS
VLYNSASFSTFKCYGVSPTKLNDLCFTNVYADSFVIRGDEVRQIAPGQTGKIADYNYKLPDDFTGCVIAWN
SNNLDSKVGGNYNYLYRLFRKSNLKPFERDISTEIYQAGSTPCNGVEGFNCYFPLQSYGFQPTNGVGYQ
PYRVVVLSFELLHAPATVCGPKKSTGGSGGSGSGGSGGSGSEKAAKAEEAARKMEELFKKHKIVAVLRA
NSVEEAIEKAVAVFAGGVHLIEITFTVPDADTVIKALSVLKEKGAIIGAGTVTSVEQARKAVESGAEFIVSPH
LDEEISQFAKEKGVFYMPGVMTPTELVKAMKLGHTILKLFPGEVVGPQFVKAMKGPFPNVKFVPTGGVNL
DNVAEWFKAGVLAVGVGSALVKGTPDEVREKAKAFVEKIRGATEGGSHHHHHHHH

>Monomeric SARS-CoV-2 RBD
MGILPSPGMPALLSLVSLLSVLLMGCVAETGTRFPNITNLCPFGEVFNATRFASVYAWNRKRISNCVADYS
VLYNSASFSTFKCYGVSPTKLNDLCFTNVYADSFVIRGDEVRQIAPGQTGKIADYNYKLPDDFTGCVIAWN
SNNLDSKVGGNYNYLYRLFRKSNLKPFERDISTEIYQAGSTPCNGVEGFNCYFPLQSYGFQPTNGVGYQ
PYRVVVLSFELLHAPATVCGPKKSTHHHHHHHH

>SARS-CoV-2 S-2P Trimer
MGILPSPGMPALLSLVSLLSVLLMGCVAETGTQCVNLTTRTQLPPAYTNSFTRGVYYPDKVFRSSVLHST
QDLFLPFFSNVTWFHAIHVSGTNGTKRFDNPVLPFNDGVYFASTEKSNIIRGWIFGTTLDSKTQSLLIVNNA
TNVVIKVCEFQFCNDPFLGVYYHKNNKSWMESEFRVYSSANNCTFEYVSQPFLMDLEGKQGNFKNLREF
VFKNIDGYFKIYSKHTPINLVRDLPQGFSALEPLVDLPIGINITRFQTLLALHRSYLTPGDSSSGWTAGAAAY
YVGYLQPRTFLLKYNENGTITDAVDCALDPLSETKCTLKSFTVEKGIYQTSNFRVQPTESIVRFPNITNLCP
FGEVFNATRFASVYAWNRKRISNCVADYSVLYNSASFSTFKCYGVSPTKLNDLCFTNVYADSFVIRGDEV
RQIAPGQTGKIADYNYKLPDDFTGCVIAWNSNNLDSKVGGNYNYLYRLFRKSNLKPFERDISTEIYQAGST
PCNGVEGFNCYFPLQSYGFQPTNGVGYQPYRVVVLSFELLHAPATVCGPKKSTNLVKNKCVNFNFNGLT
GTGVLTESNKKFLPFQQFGRDIADTTDAVRDPQTLEILDITPCSFGGVSVITPGTNTSNQVAVLYQDVNCT
EVPVAIHADQLTPTWRVYSTGSNVFQTRAGCLIGAEHVNNSYECDIPIGAGICASYQTQTNSPSGAGSVA
SQSIIAYTMSLGAENSVAYSNNSIAIPTNFTISVTTEILPVSMTKTSVDCTMYICGDSTECSNLLLQYGSFCT
QLNRALTGIAVEQDKNTQEVFAQVKQIYKTPPIKDFGGFNFSQILPDPSKPSKRSFIEDLLFNKVTLADAGFI
KQYGDCLGDIAARDLICAQKFNGLTVLPPLLTDEMIAQYTSALLAGTITSGWTFGAGAALQIPFAMQMAYR
FNGIGVTQNVLYENQKLIANQFNSAIGKIQDSLSSTASALGKLQDVVNQNAQALNTLVKQLSSNFGAISSVL
NDILSRLDPPEAEVQIDRLITGRLQSLQTYVTQQLIRAAEIRASANLAATKMSECVLGQSKRVDFCGKGYH
LMSFPQSAPHGVVFLHVTYVPAQEKNFTTAPAICHDGKAHFPREGVFVSNGTHWFVTQRNFYEPQIITTD

NTFVSGNCDVVIGIVNNTVYDPLQPELDSFKEELDKYFKNHTSPDVDLGDISGINASVVNIQKEIDRLNEVA
KNLNESLIDLQELGKYEQYIKGSGRENLYFQGGGGSGYIPEAPRDGQAYVRKDGEWVLLSTFLGHHHHH
HHH

>hACE2(R518G)-Fc
MARAWIFFLLCLAGRALASTIEEQAKTFLDKFNHEAEDLFYQSSLASWNYNTNITEENVQNMNNAGDKWS
AFLKEQSTLAQMYPLQEIQNLTVKLQLQALQQNGSSVLSEDKSKRLNTILNTMSTIYSTGKVCNPDNPQE
CLLLEPGLNEIMANSLDYNERLWAWESWRSEVGKQLRPLYEEYVVLKNEMARANHYEDYGDYWRGDY
EVNGVDGYDYSRGQLIEDVEHTFEEIKPLYEHLHAYVRAKLMNAYPSYISPIGCLPAHLLGDMWGRFWTN
LYSLTVPFGQKPNIDVTDAMVDQAWDAQRIFKEAEKFFVSVGLPNMTQGFWENSMLTDPGNVQKAVCH
PTAWDLGKGDFRILMCTKVTMDDFLTAHHEMGHIQYDMAYAAQPFLLRNGANEGFHEAVGEIMSLSAAT
PKHLKSIGLLSPDFQEDNETEINFLLKQALTIVGTLPFTYMLEKWRWMVFKGEIPKDQWMKKWWEMKREI
VGVVEPVPHDETYCDPASLFHVSNDYSFIRYYTGTLYQFQFQEALCQAAKHEGPLHKCDISNSTEAGQKL
FNMLRLGKSEPWTLALENVVGAKNMNVRPLLNYFEPLFTWLKDQNKNSFVGWSTDWSPYADPLVPRGS
GGGGDPEPKSCDKTHTCPPCPAPELLGGPSVFLFPPKPKDTLMISRTPEVTCVVVDVSHEDPEVKFNWY
VDGVEVHNAKTKPREEQYNSTYRVVSVLTVLHQDWLNGKEYKCKVSNKALPAPIEKTISKAKGQPREPQ
VYTLPPSRDELTKNQVSLTCLVKGFYPSDIAVEWESNGQPENNYKTTPPVLDSDGSFFLYSKLTVDKSRW
QQGNVFSCSVMHEALHNHYTQKSLSLSPGK

>mACE2-Fc
MPMGSLQPLATLYLLGMLVASVLAQSTIEEQAKTFLDKFNHEAEDLFYQSSLASWNYNTNITEENVQNMN
NAGEKWSAFLKEQSTLAQMYPLQEIQNLTVKLQLQALQQNGSSVLSEDKSKRLNTILNTMSTIYSTGKVC
NPNNPQECLLLDPGLNEIMEKSLDYNERLWAWEGWRSEVGKQLRPLYEEYVVLKNEMARANHYKDYGD
YWRGNYEVNGVDGYDYNRDQLIEDVERTFEEIKPLYEHLHAYVRAKLMNAYPSYISPTGCLPAHLLGDM
WGRFWTNLYSLTVPFGQKPNIDVTDAMVNQAWNAQRIFKEAEKFFVSVGLPNMTQGFWENSMLTDPGN
VQKVVCHPTAWDLGKGDFRIIMCTKVTMDDFLTAHHEMGHIQYDMAYAAQPFLLRNGANEGFHEAVGEI
MSLSAATPKHLKSIGLLSPDFQEDNETEINFLLKQALTIVGTLPFTYMLEKWRWMVFKGEIPKDQWMKKW
WEMKREIVGVVEPVPHDETYCDPASLFHVSNDYSFIRYYTRTLYQFQFQEALCQAAKHEGPLHKCDISNS
TEAGQKLLNMLKLGKSEPWTLALENVVGAKNMNVRPLLNYFEPLFTWLKDQNKNSFVGWSTDWSPYAD
QSIKVRISLKSALGDKAYEWNDNEMYLFRSSVAYAMRTYFLEIKHQTILFGEEDVRVADLKPRISFNFYVTA
PKNVSDIIPRTEVEEAIRISRSRINDAFRLNDNSLEFLGIQTTLAPPYQSPVTDPLVPRGSGGGGDPEPKSC
DKTHTCPPCPAPELLGGPSVFLFPPKPKDTLMISRTPEVTCVVVDVSHEDPEVKFNWYVDGVEVHNAKT
KPREEQYNSTYRVVSVLTVLHQDWLNGKEYKCKVSNKALPAPIEKTISKAKGQPREPQVYTLPPSRDELT
KNQVSLTCLVKGFYPSDIAVEWESNGQPENNYKTTPPVLDSDGSFFLYSKLTVDKSRWQQGNVFSCSV
MHEALHNHYTQKSLSLSPGK
