## Supplementary Item 2 for "Elicitation of potent neutralizing antibody responses by designed protein nanoparticle vaccines for SARS-CoV-2"

**Walls AC & Fiala B, et al.**

### Supplemental Stability Data

### SDS-PAGE for Monomeric RBD

D1 D3 D7 D14 D21 D28

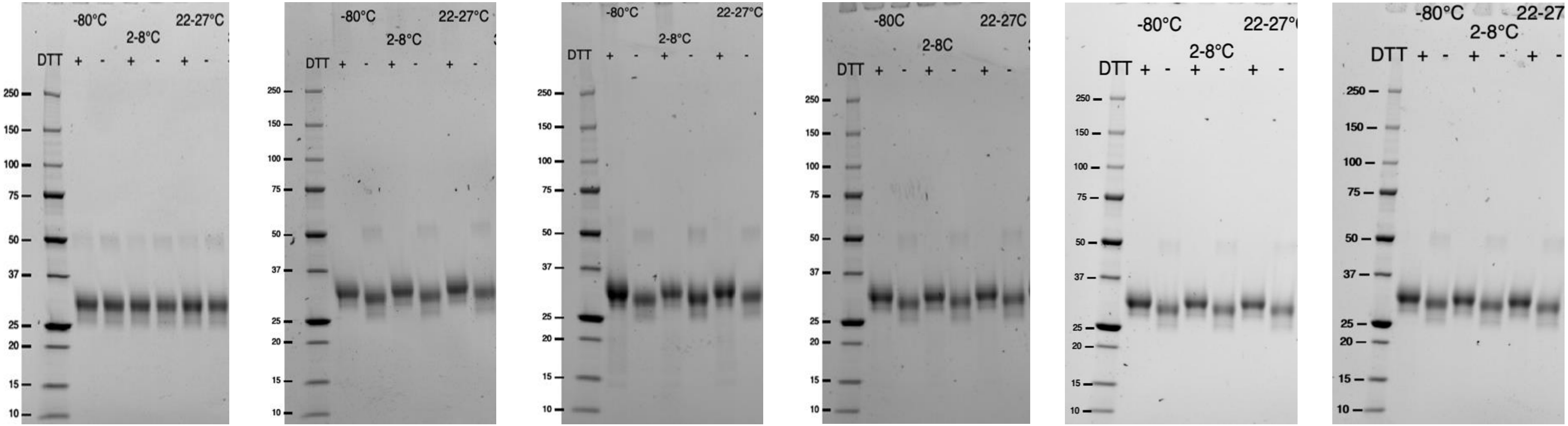

### mACE2-Fc Binding Relative to <-70°C Reference for Monomeric RBD

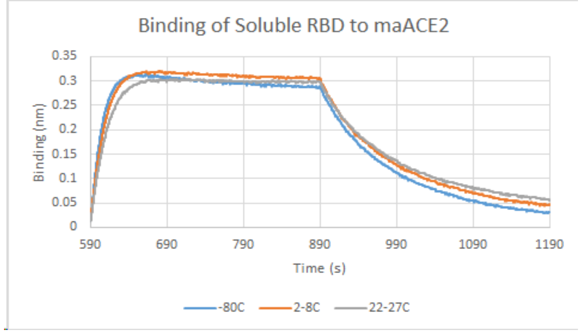


D1

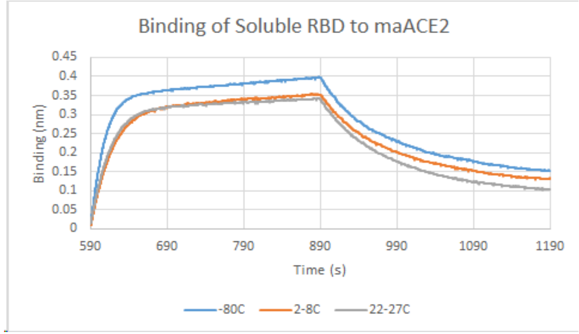


D3

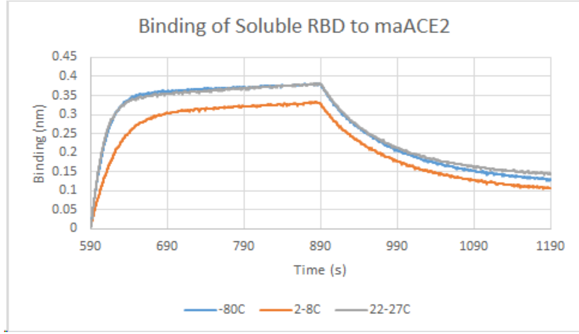


D7

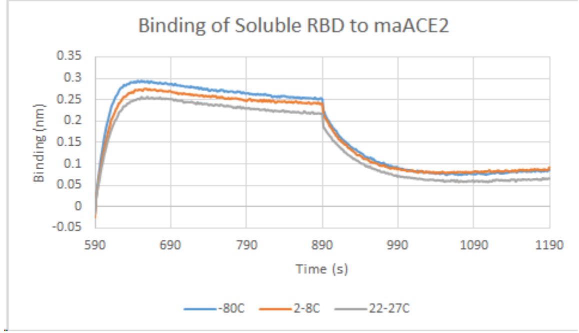


D14

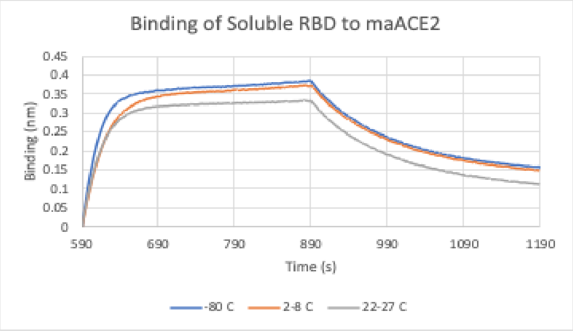


D21

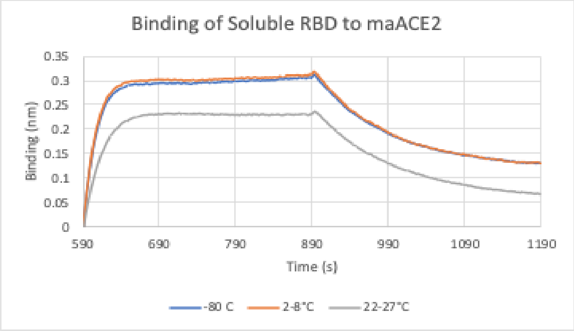


D28

### CR3022 IgG Binding Relative to <-70°C Reference for Monomeric RBD

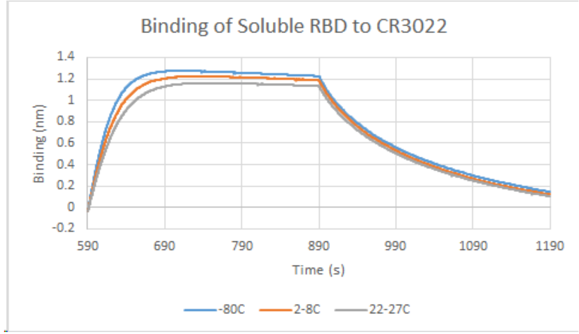


D1

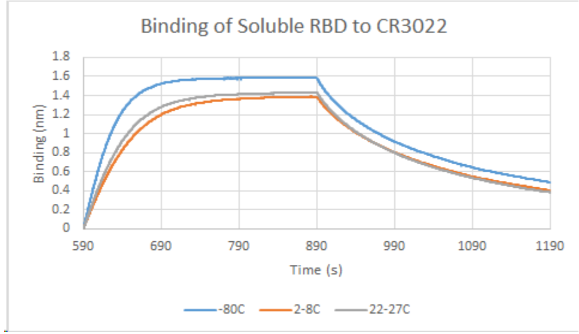


D3

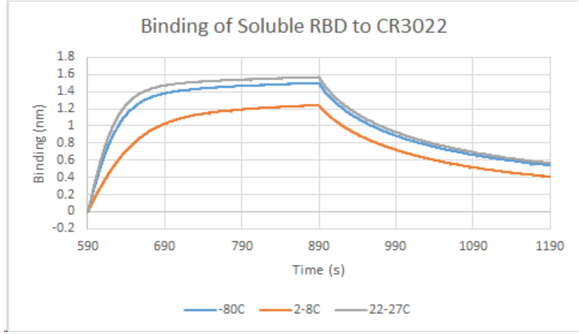


D7

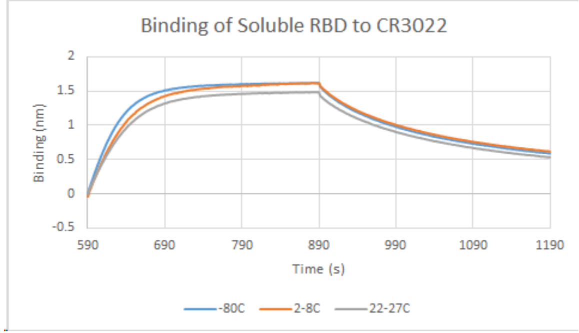


D14

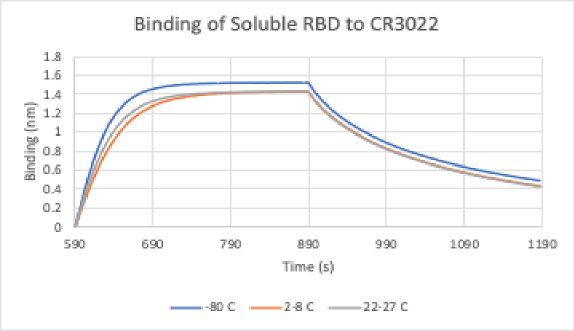


D21

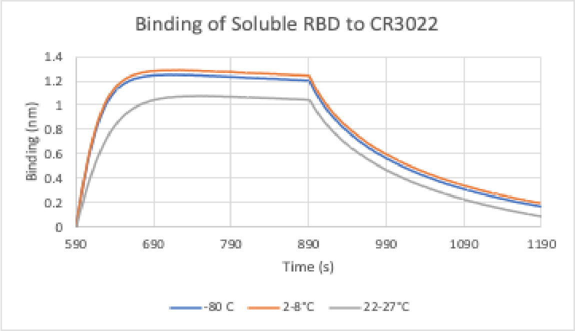


D28

### Absorbance at 320/280 nm for Monomeric RBD

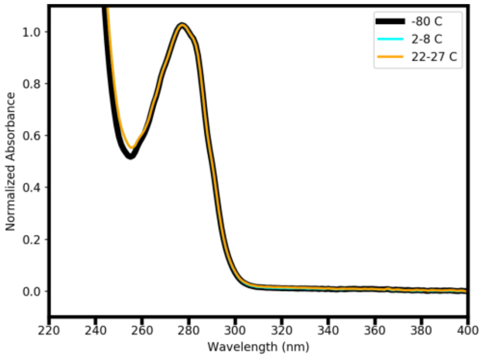


D1

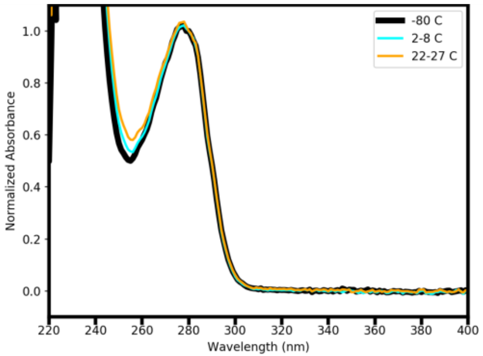


D3

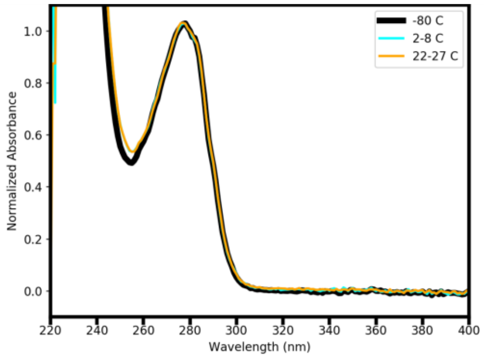


D7

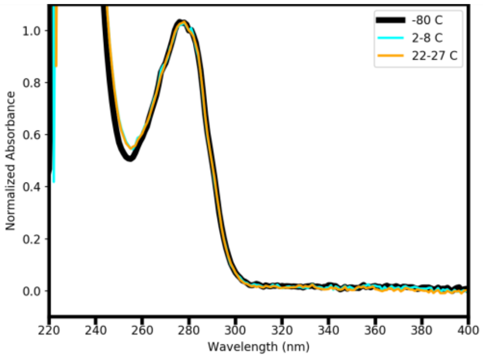


D14

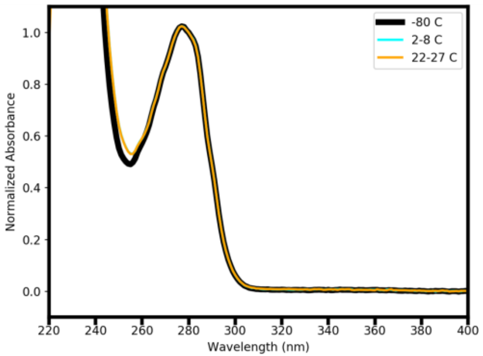


D21

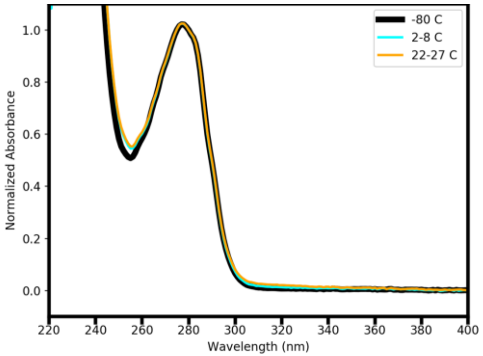


D28

### nsEM for S-2P Trimer, -80°C

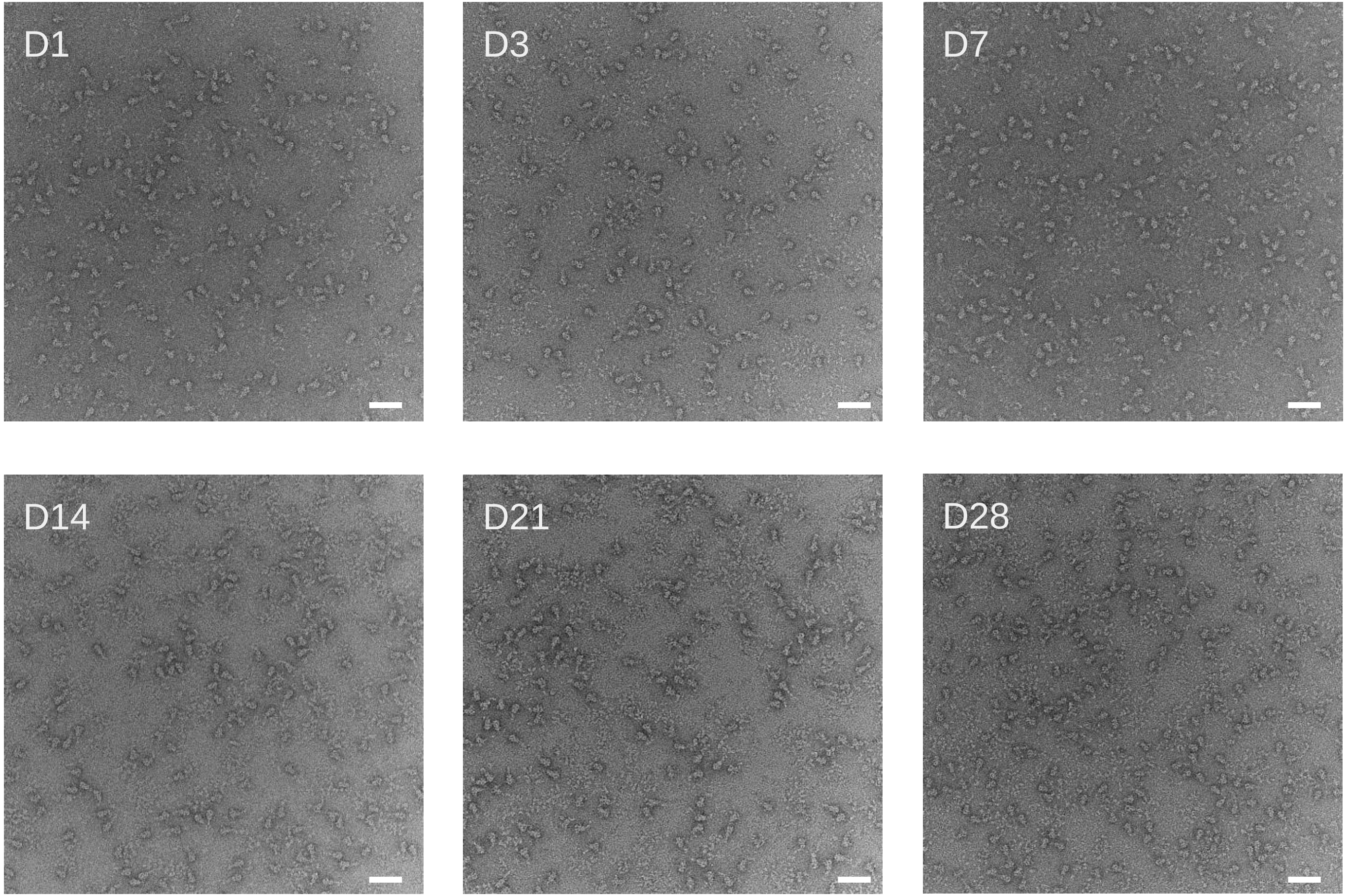


Scale bars: 50 nm

### nsEM for S-2P Trimer, 2-8°C

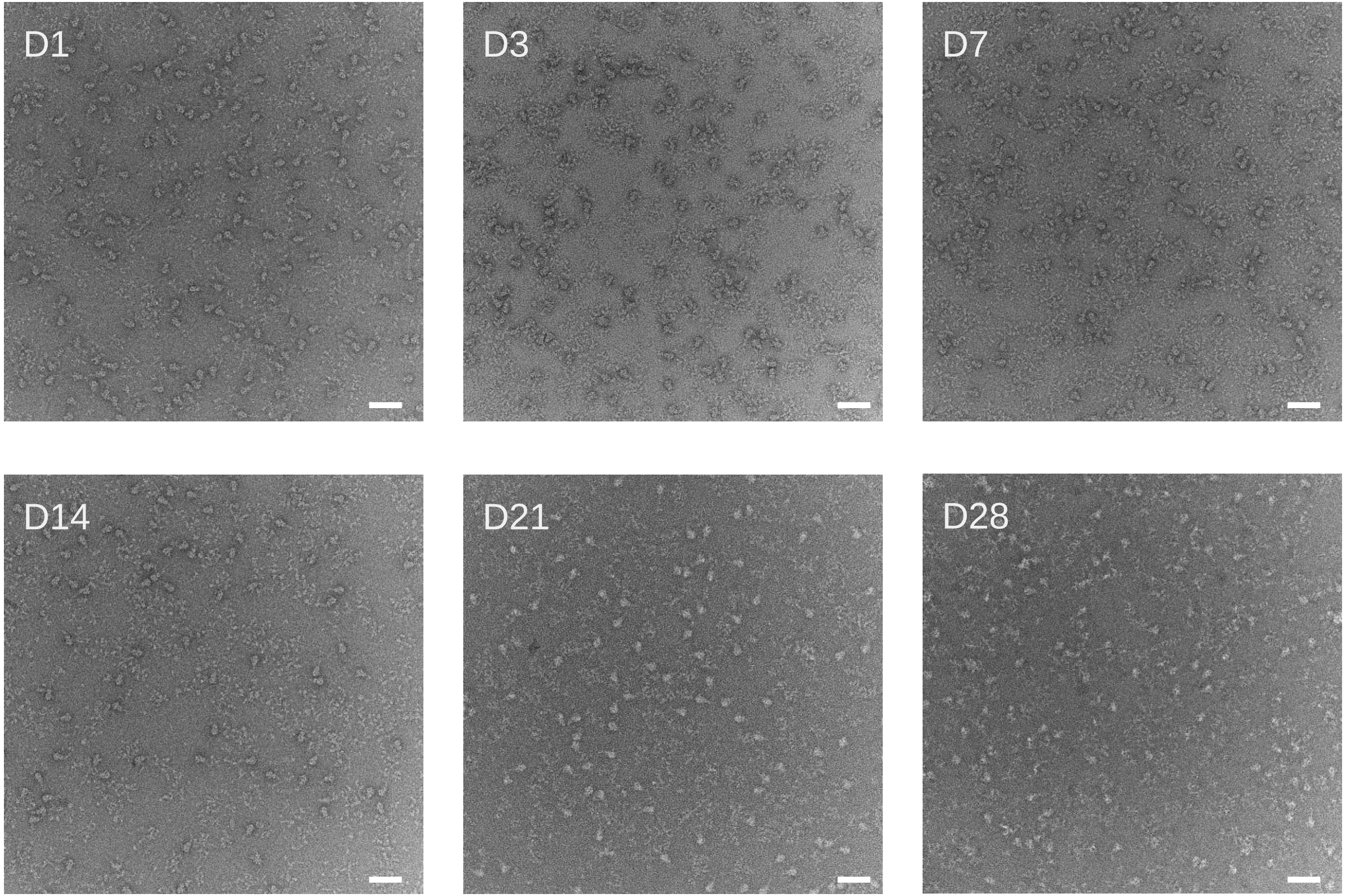


Scale bars: 50 nm

### nsEM for S-2P Trimer, 22-27°C

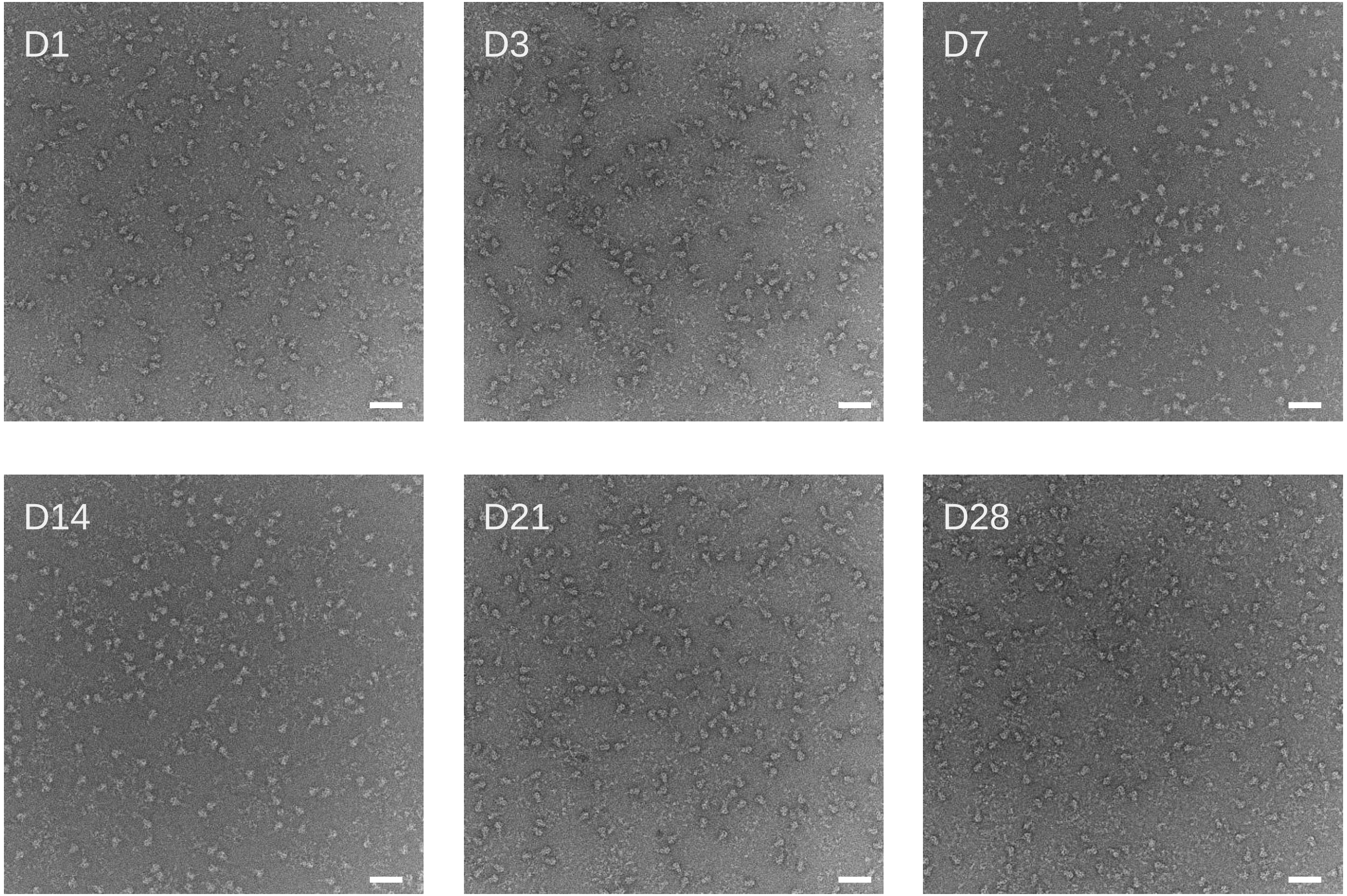


Scale bars: 50 nm

### SDS-PAGE for RBD-8GS-I53-50A

D1 D3 D7 D14 D21 D28

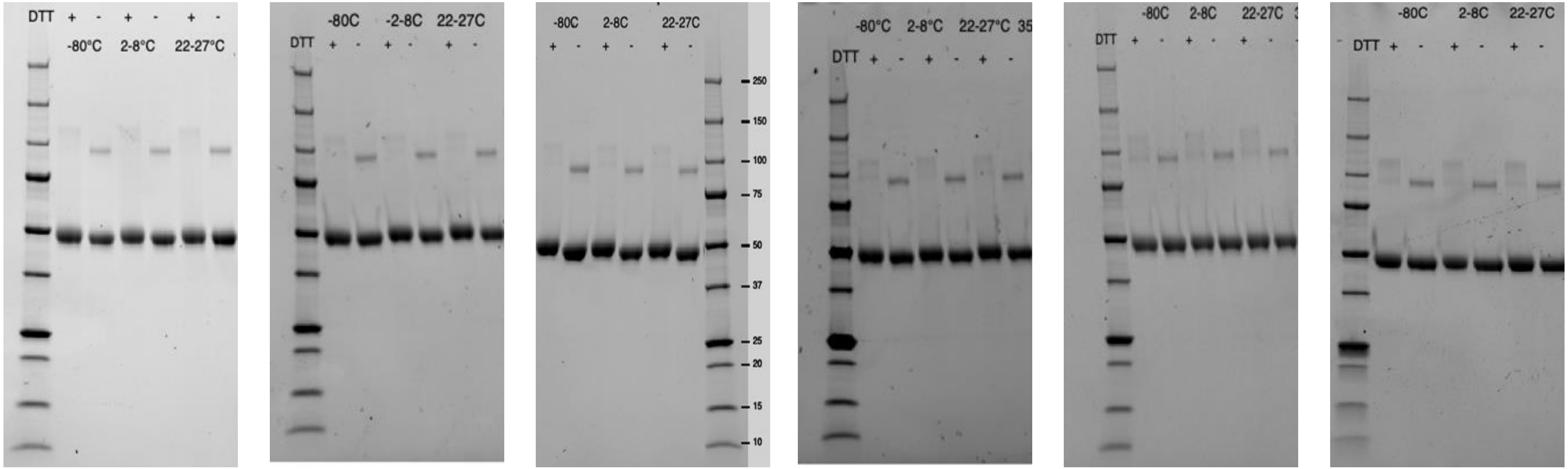

### Absorbance at 320/280 for RBD-8GS-I53-50A

Not measured

D1

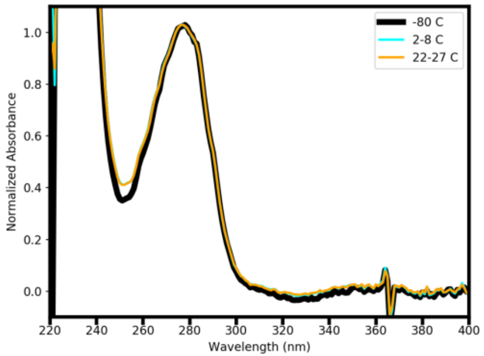


D3

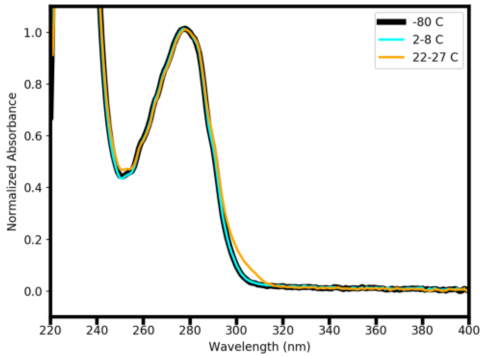


D7

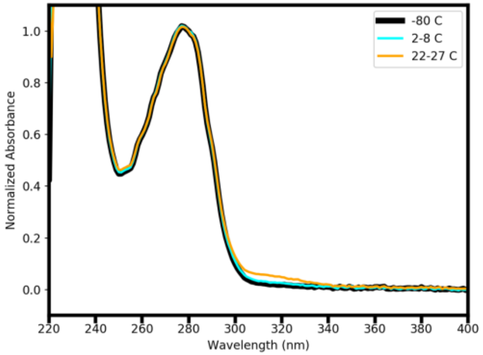


D14

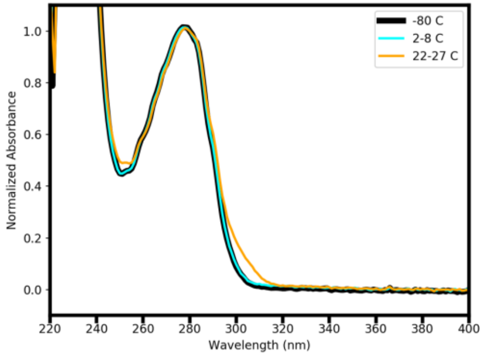


D21

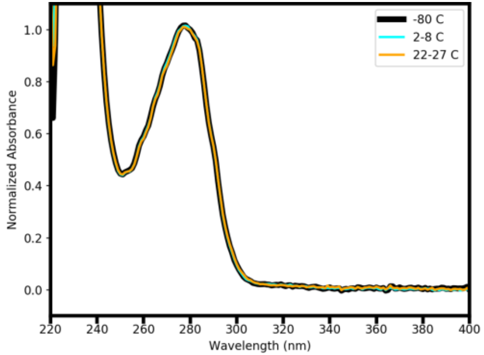


D28

### mACE2-Fc Binding Relative to <-70°C Reference for RBD-8GS-I53-50A

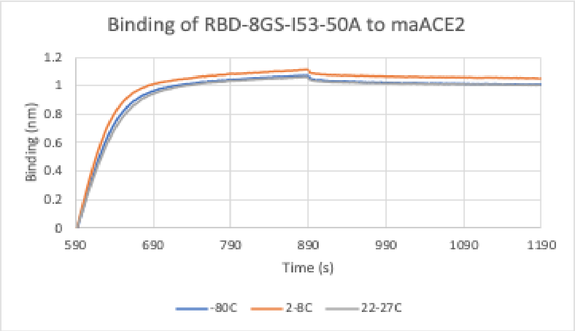


D1

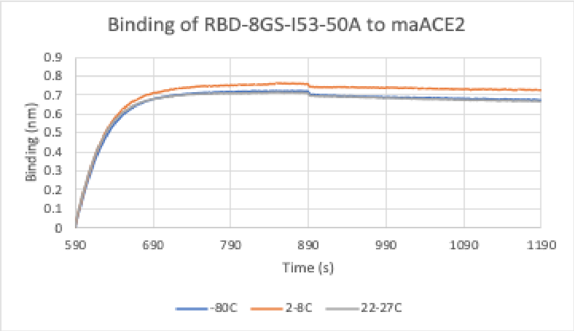


D3

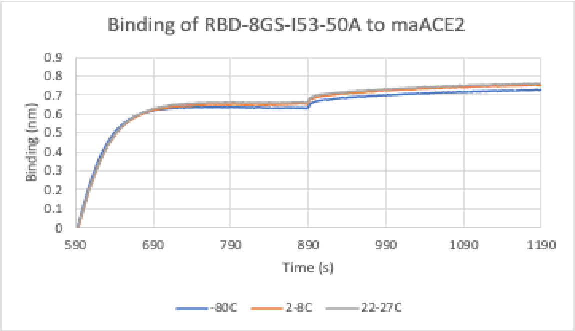


D7

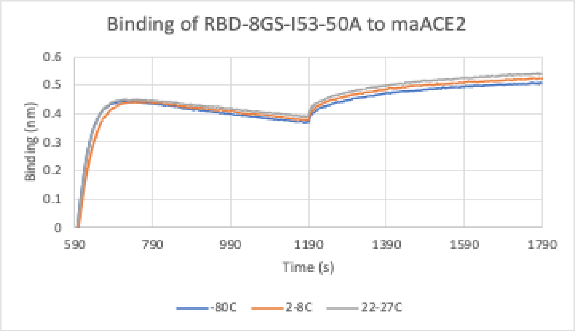


D14

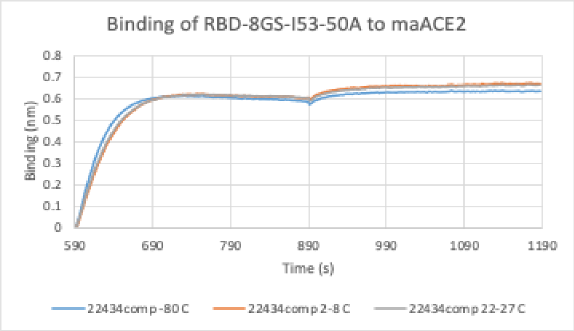


D21

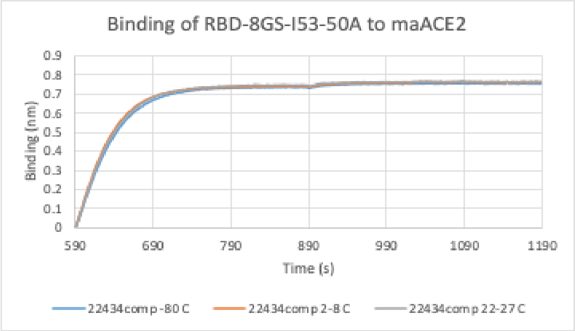


D28

### CR3022 IgG Binding Relative to <-70°C Reference for RBD-8GS-I53-50A

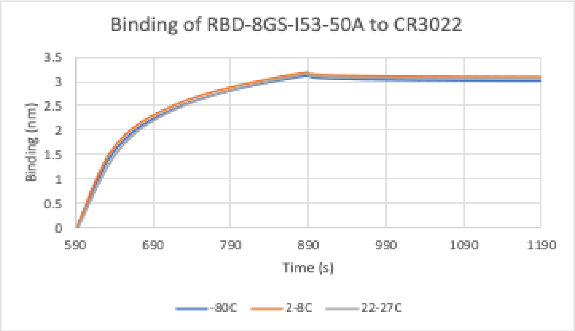


D1

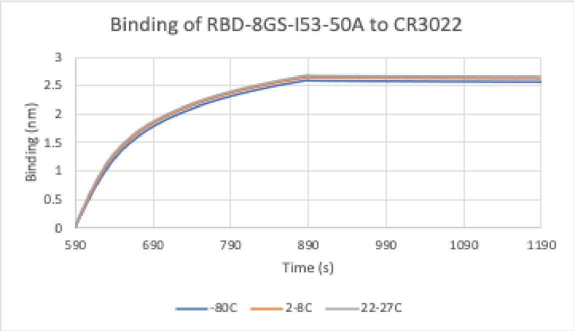


D3

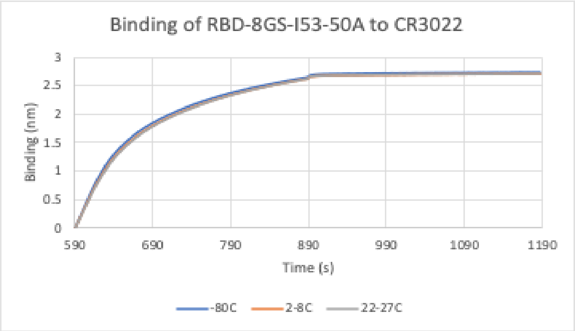


D7

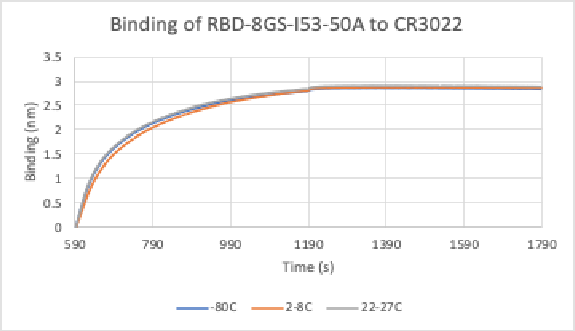


D14

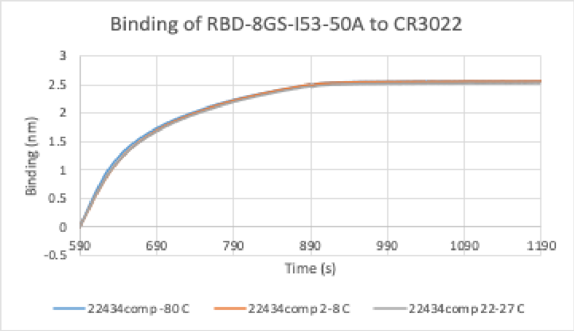


D21

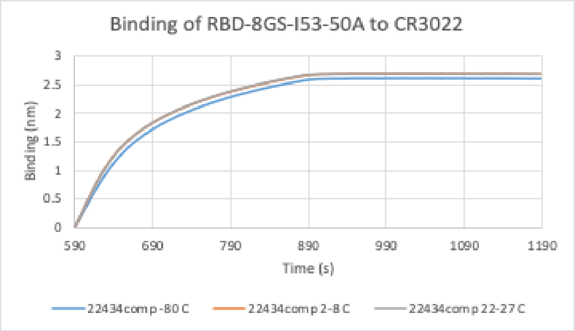


D28

### SDS-PAGE for RBD-12GS-I53-50A

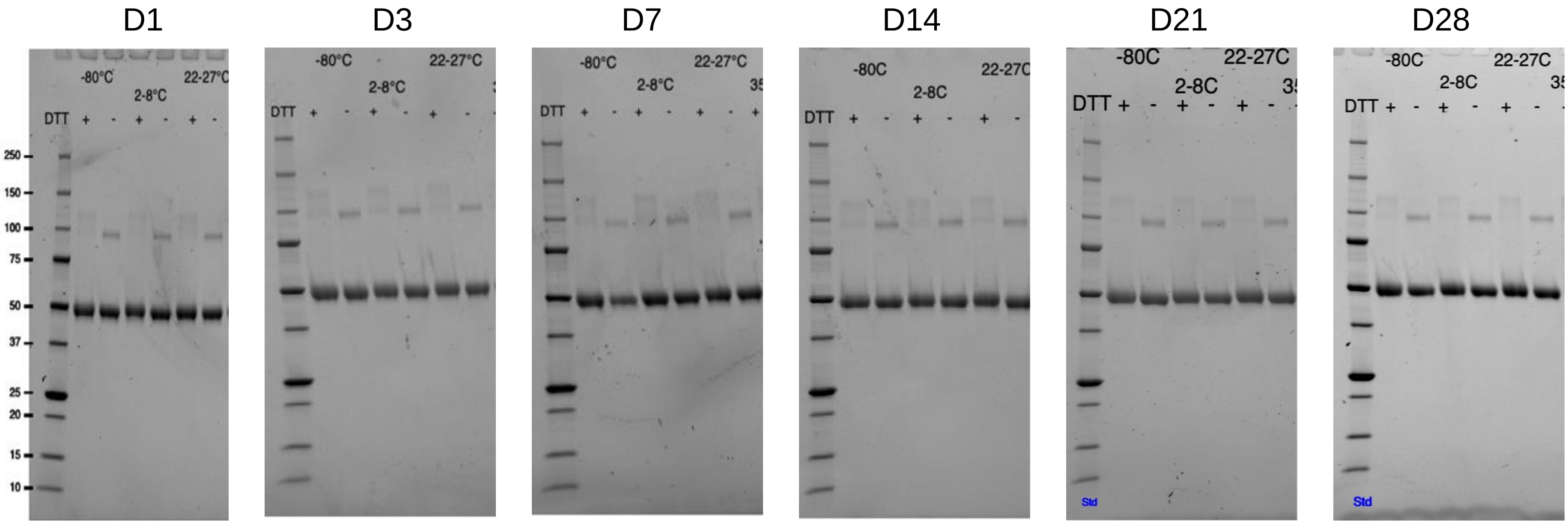

### Absorbance at 320/280 nm for RBD-12GS-I53-50A

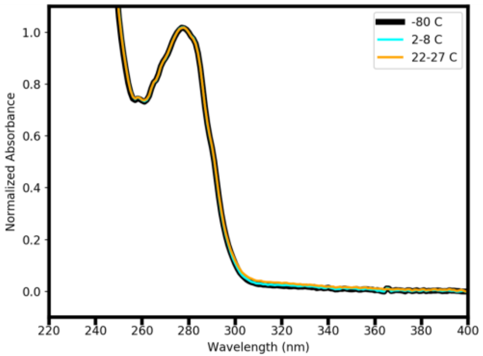


D1

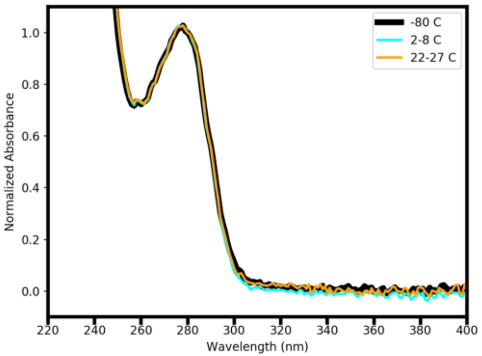


D3

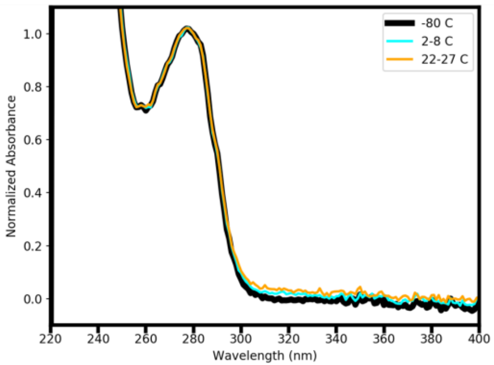


D7

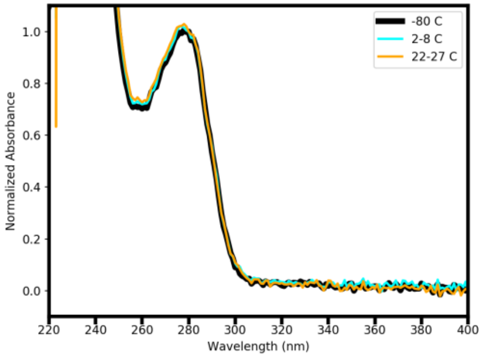


D14

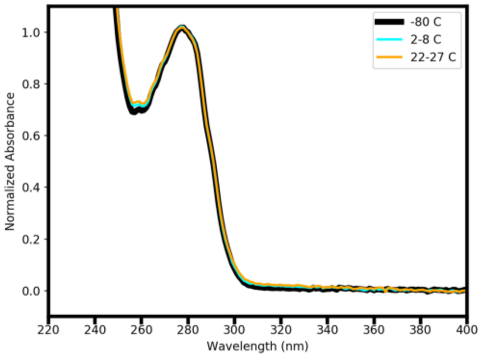


D21

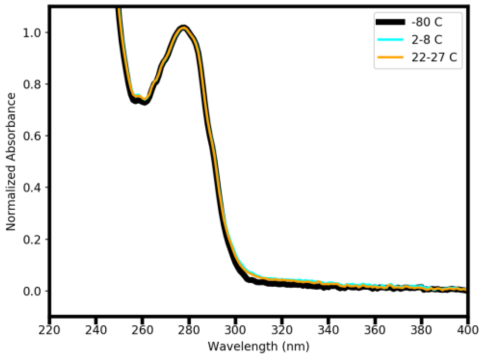


D28

### mACE2-Fc Binding Relative to <-70°C Reference for RBD-12GS-I53-50A

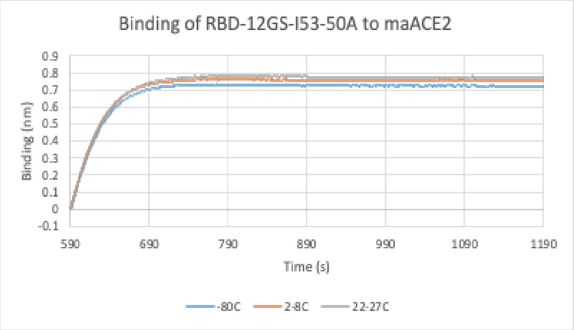


D1

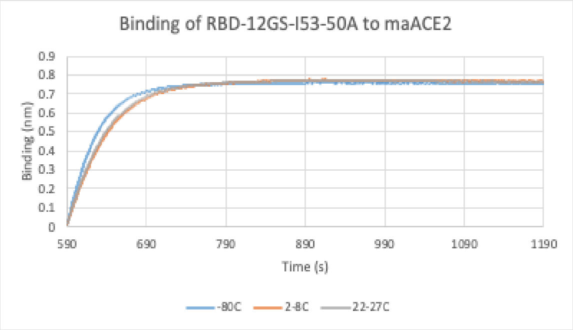


D3

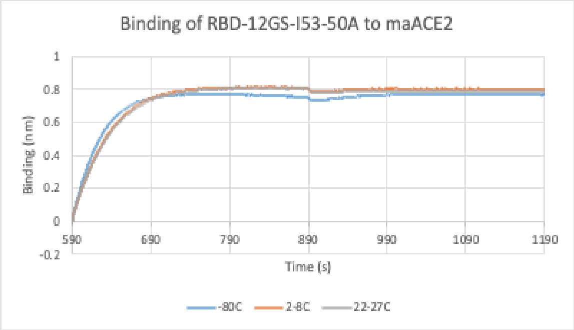


D7

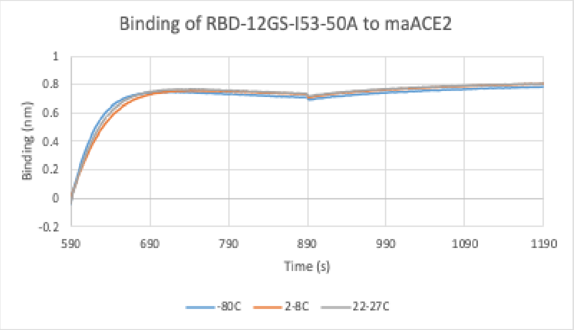


D14

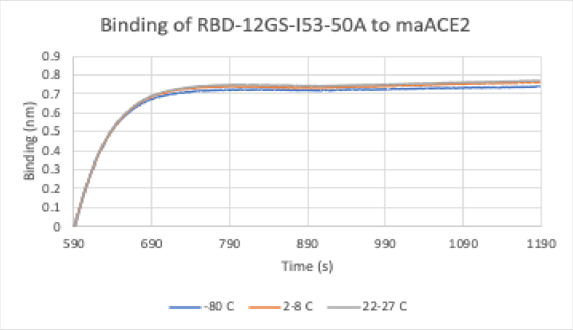


D21

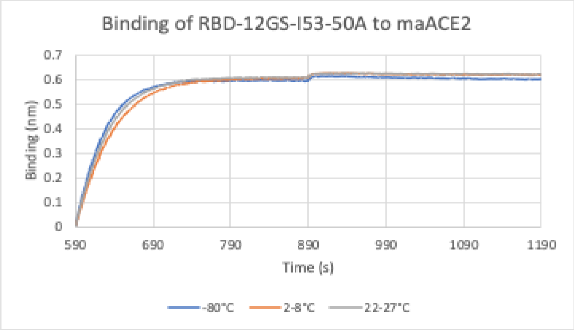


D28

### CR3022 IgG Binding Relative to <-70°C Reference for RBD-12GS-I53-50A

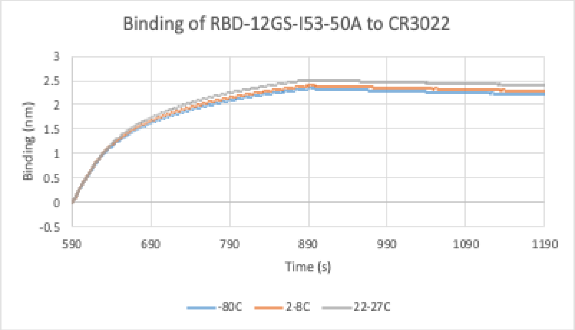


D1

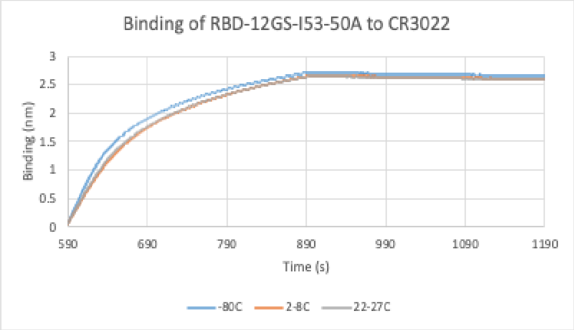


D3

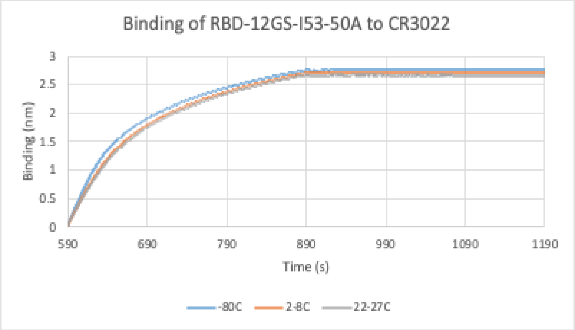


D7

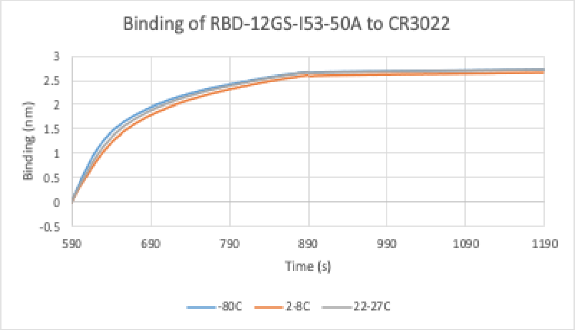


D14

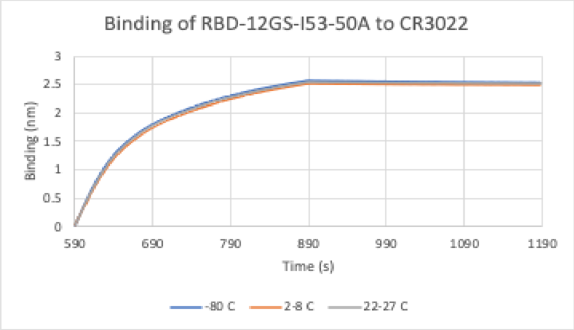


D21

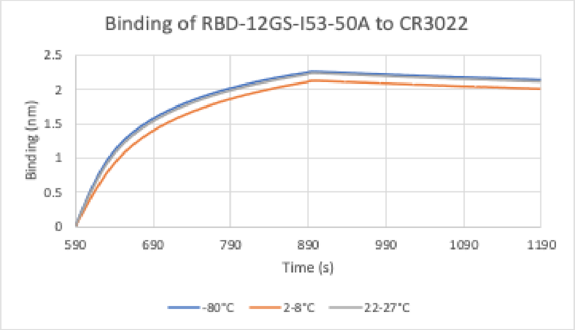


D28

### SDS-PAGE for RBD-16GS-I53-50A

D1

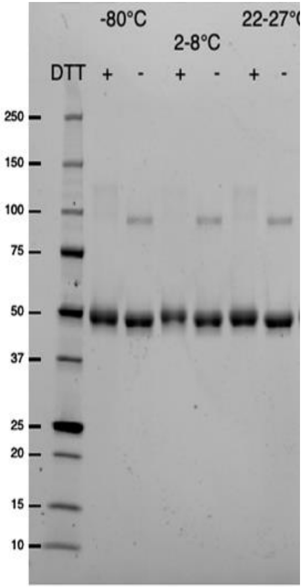


D3

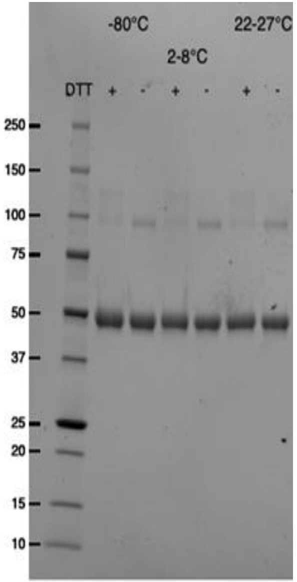


D7

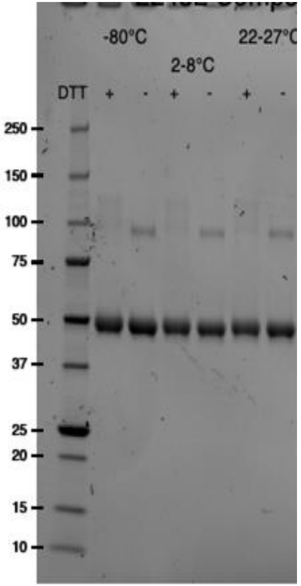


D14

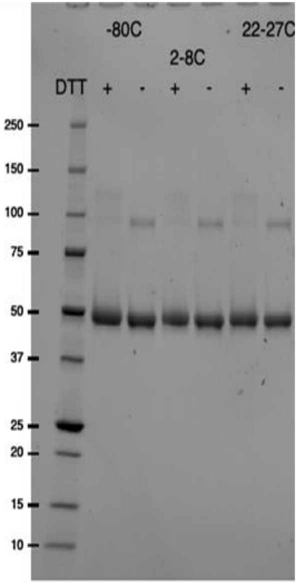


D21

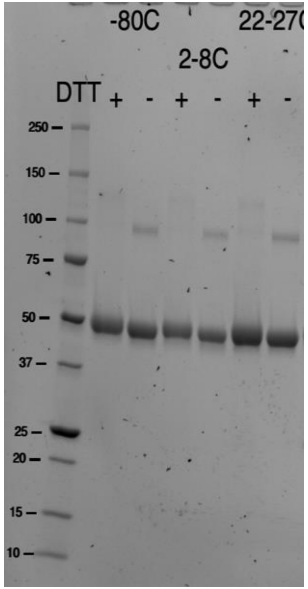


D28

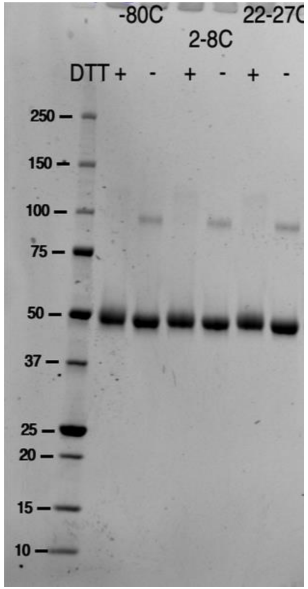

### Absorbance at 320/280 for RBD-16GS-I53-50A

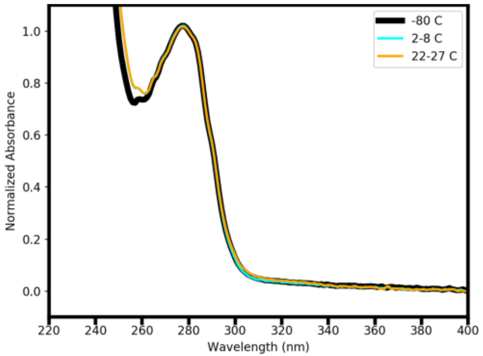


D1

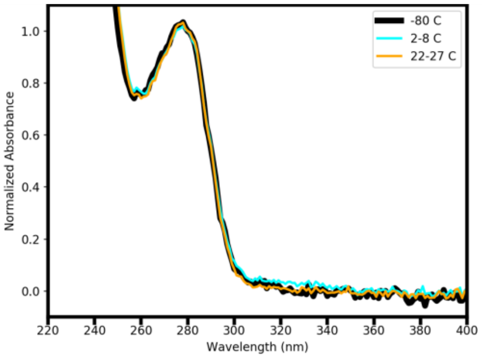


D3

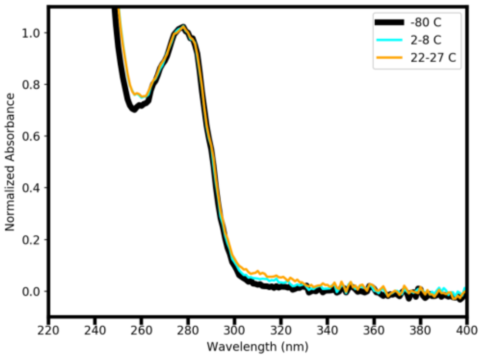


D7

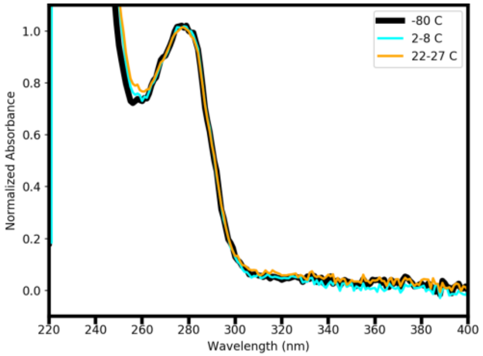


D14

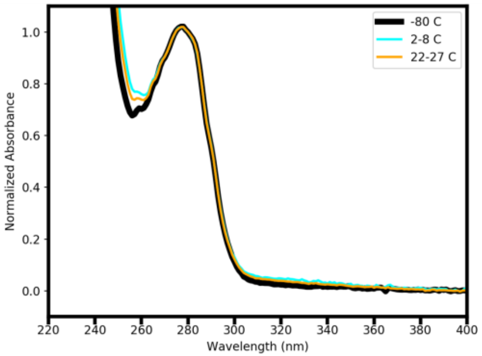


D21

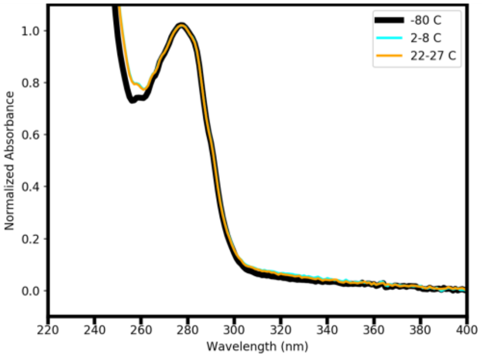


D28

### mACE2-Fc Binding Relative to <-70°C Reference for RBD-16GS-I53-50A

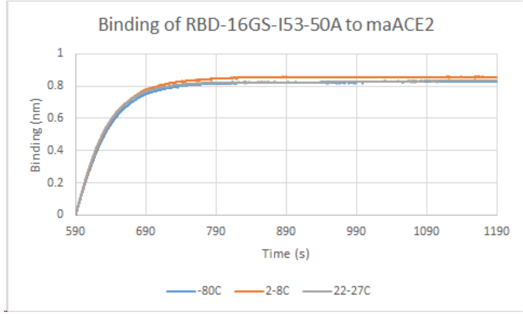


D1

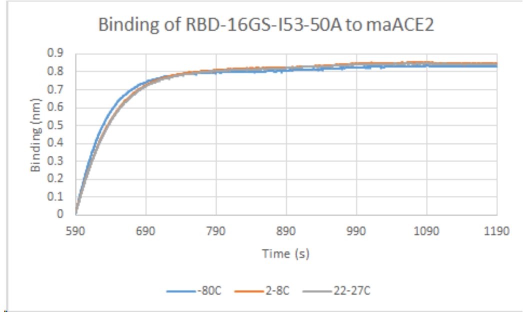


D3

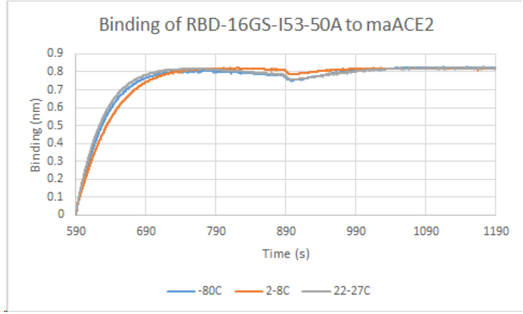


D7

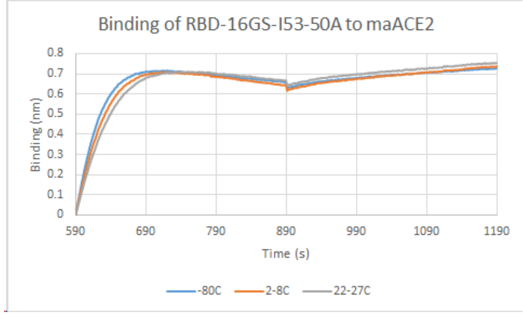


D14

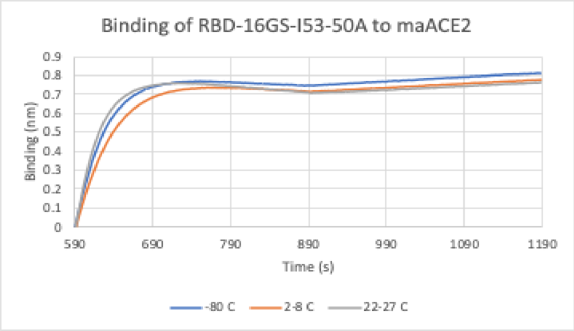


D21

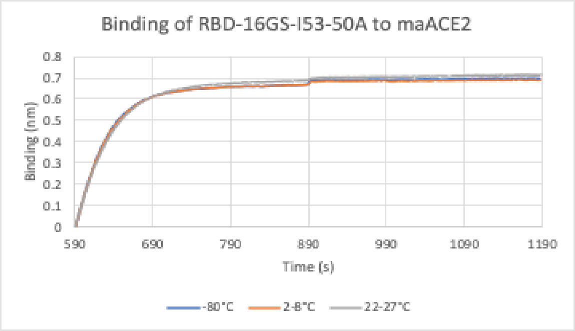


D28

### CR3022 IgG Binding Relative to <-70°C Reference for RBD-16GS-I53-50A

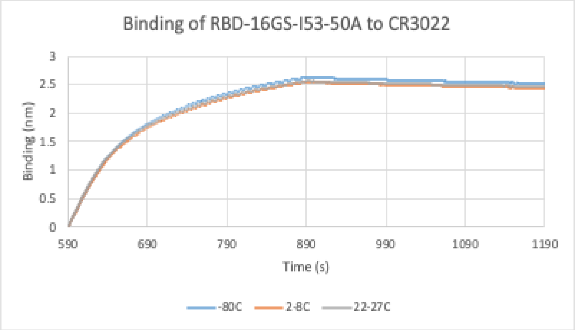


D1

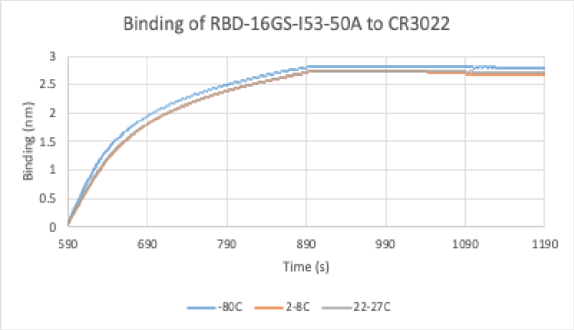


D3

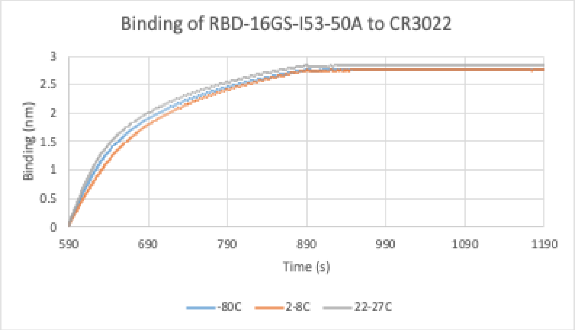


D7

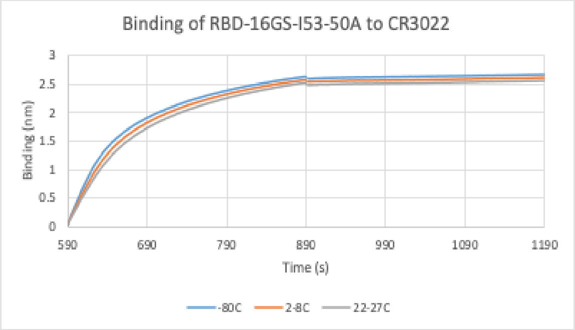


D14

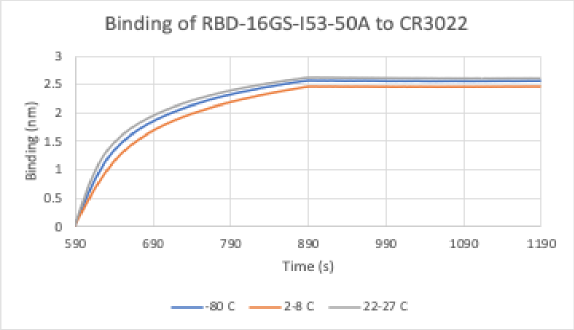


D21

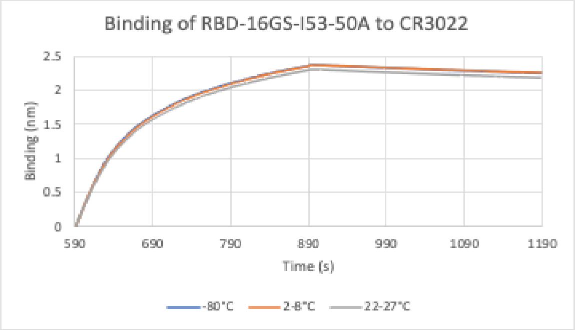


D28

### Absorbance at 320/280 nm for RBD-12GS-I53-50

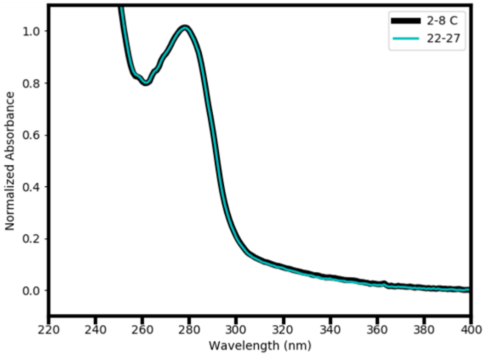

D1

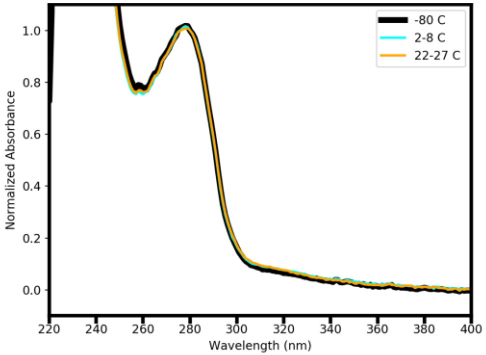

D3

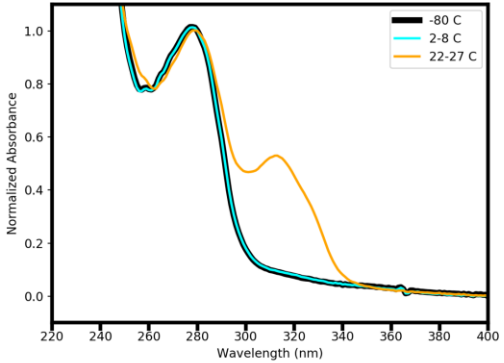

D7

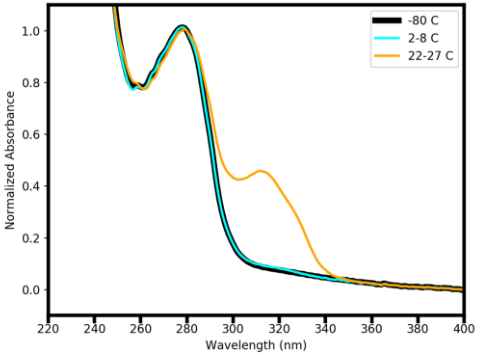

D14

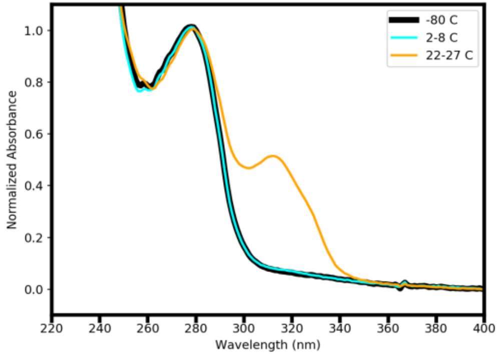

D21

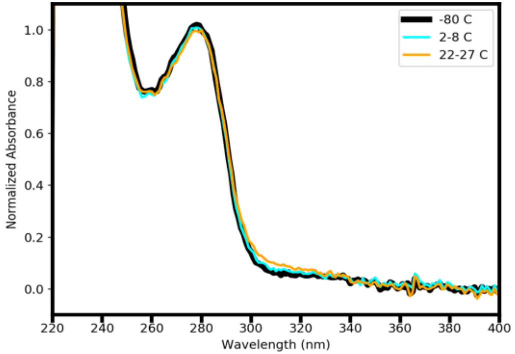

D28

### nsEM for RBD-12GS-I53-50

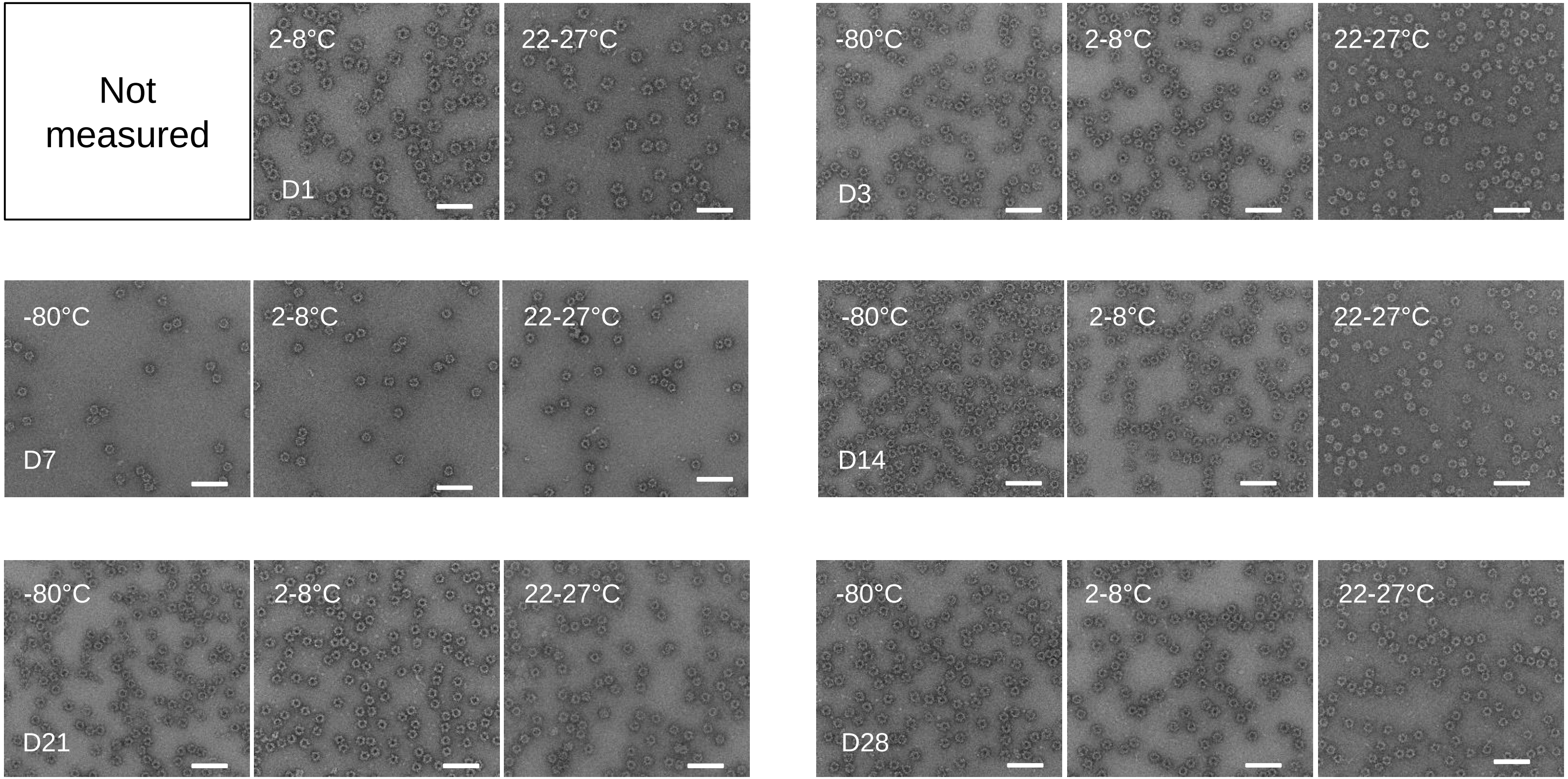


Scale bars: 100 nm
