## Supplementary Item 3 for "Elicitation of potent neutralizing antibody responses by designed protein nanoparticle vaccines for SARS-CoV-2"

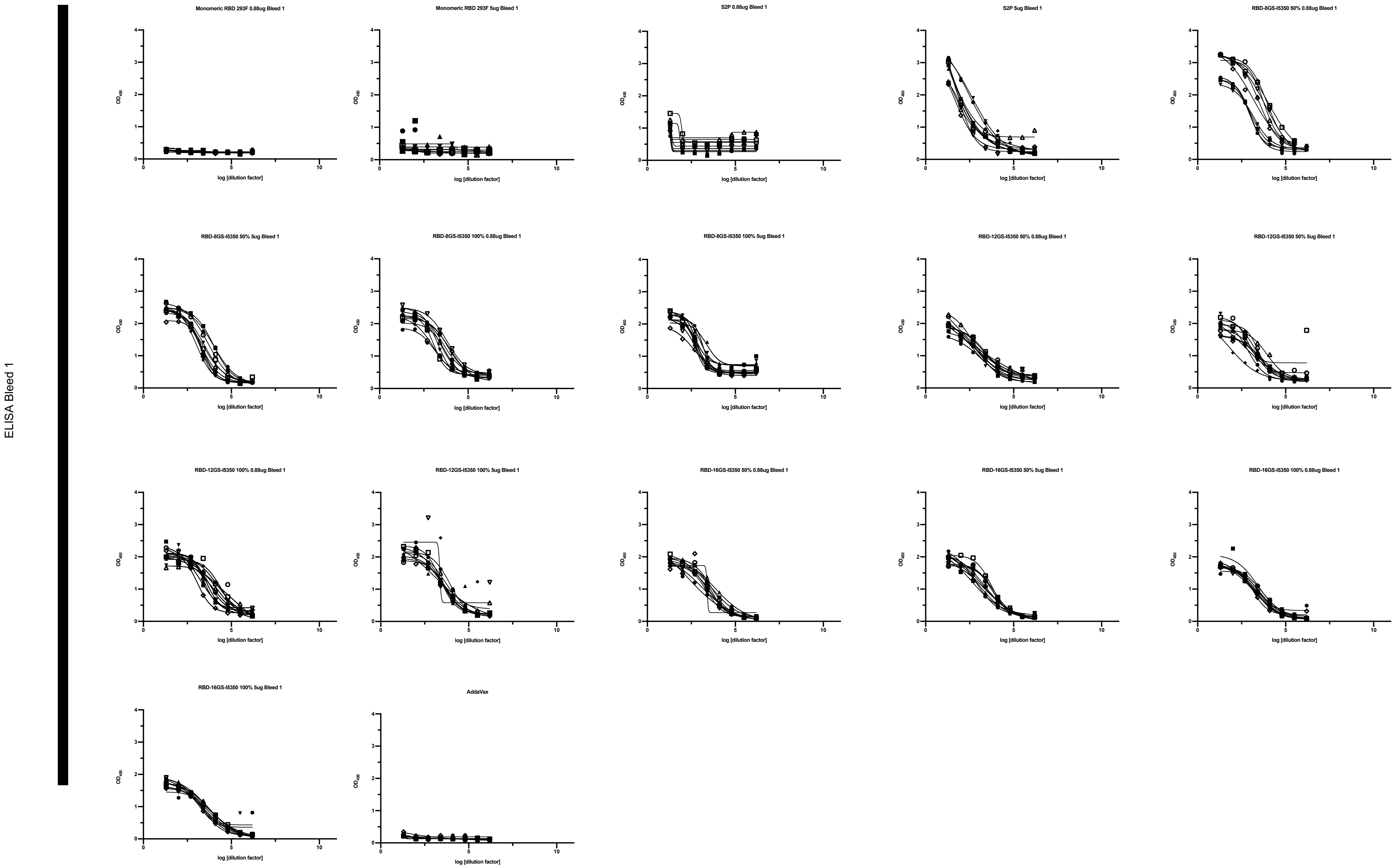
ELISA Bleed 1
Monomeric RBD 293F 0.88ug Bleed 1
Monomeric RBD 293F 5ug Bleed 1
S2P 0.88ug Bleed 1
S2P 5ug Bleed 1
RBD-8GS-I5350 50% 0.88ug Bleed 1
RBD-8GS-I5350 50% 5ug Bleed 1
RBD-8GS-I5350 100% 0.88ug Bleed 1
RBD-8GS-I5350 100% 5ug Bleed 1
RBD-12GS-I5350 50% 0.88ug Bleed 1
RBD-12GS-I5350 50% 5ug Bleed 1
RBD-12GS-I5350 100% 0.88ug Bleed 1
RBD-12GS-I5350 100% 5ug Bleed 1
RBD-16GS-I5350 50% 0.88ug Bleed 1
RBD-16GS-I5350 50% 5ug Bleed 1
RBD-16GS-I5350 100% 0.88ug Bleed 1
RBD-16GS-I5350 100% 5ug Bleed 1
AddaVax
$OD_{450}$
log [dilution factor]

ELISA Bleed 2

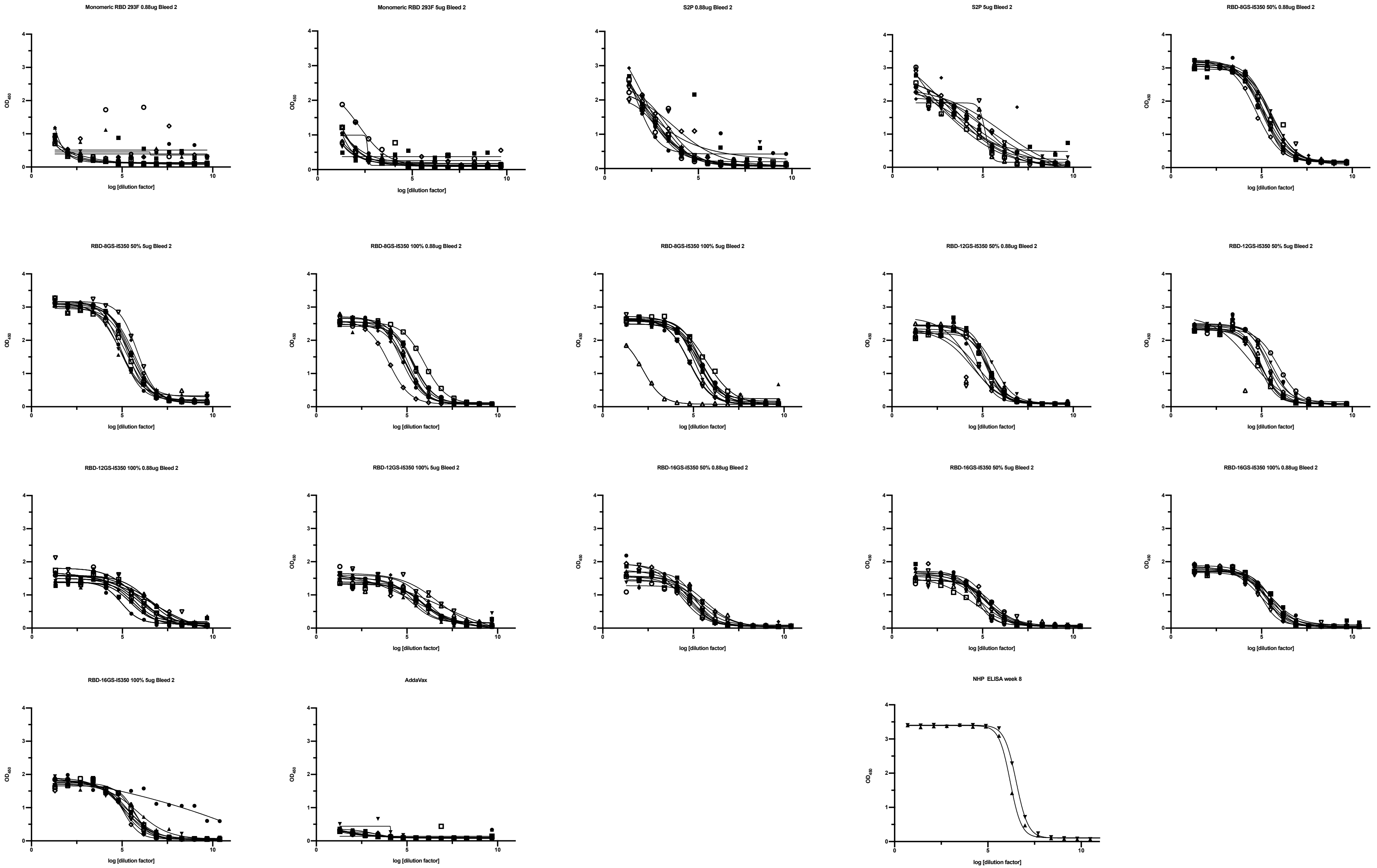

pseudoneutralization Bleed 1

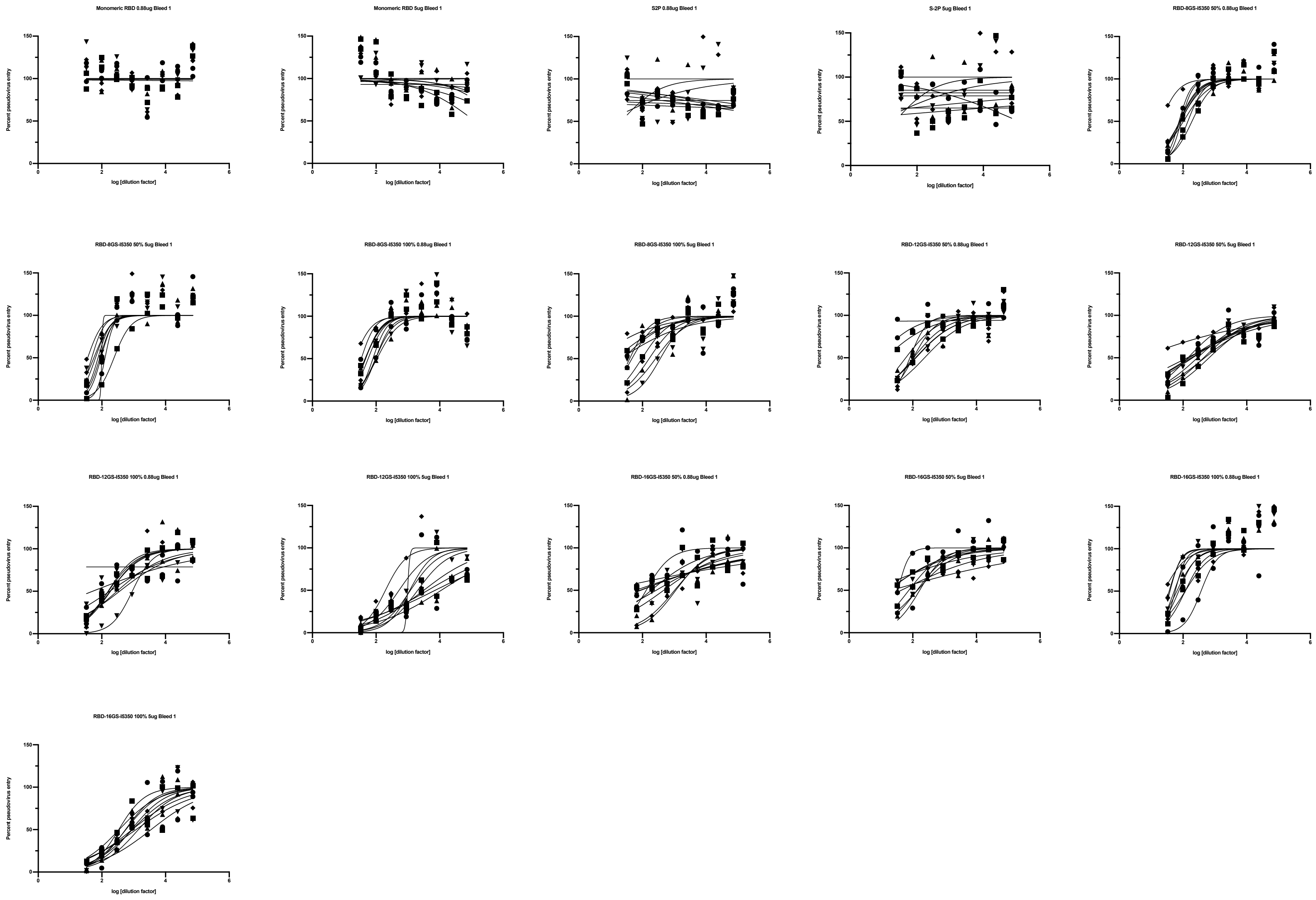

pseudoneutralization Bleed 2

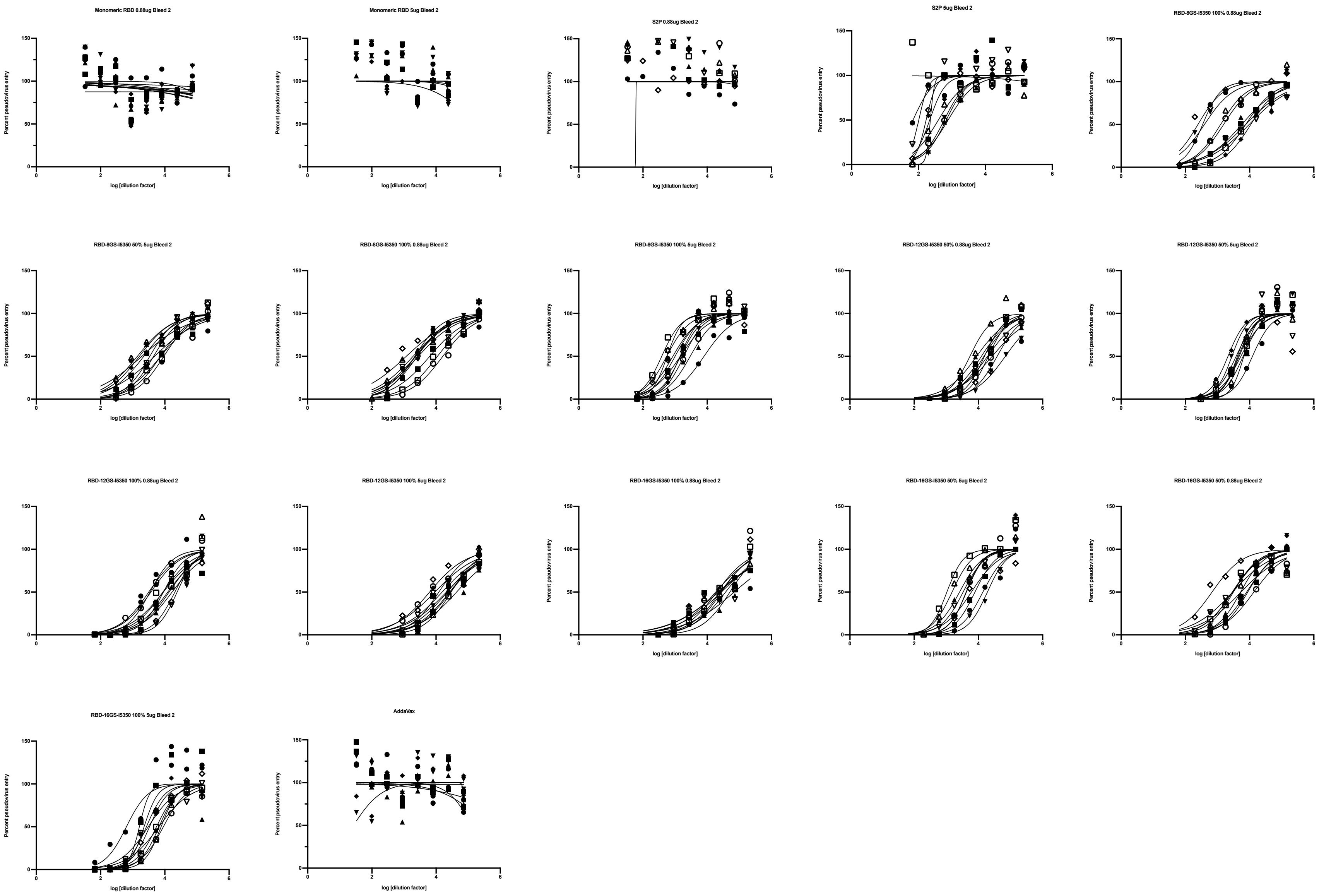

### Live virus neutralization Bleed 1

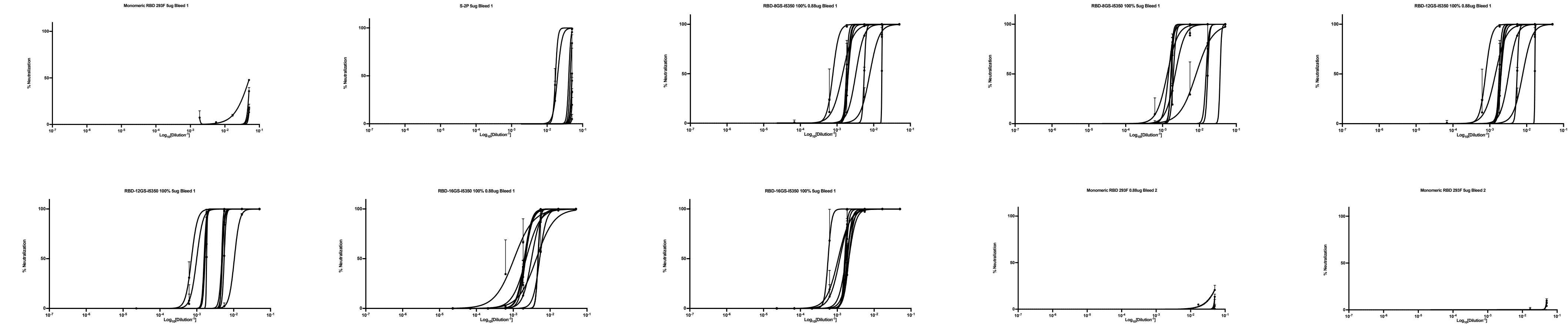

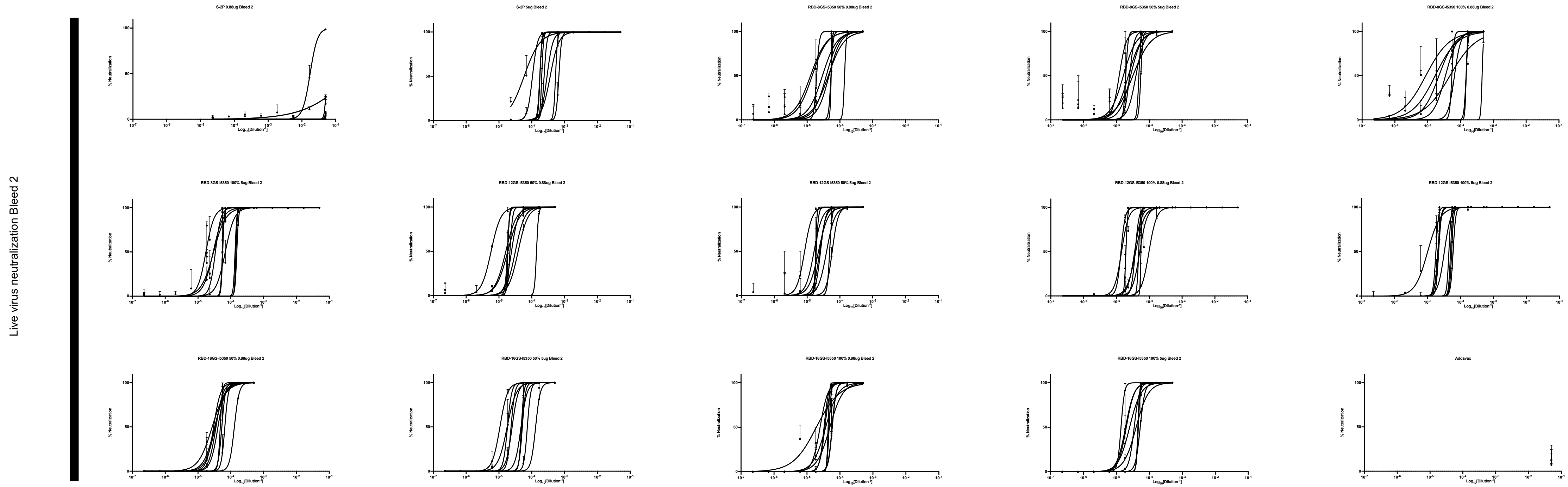

Live virus neutralization Bleed 2
S-2P 0.88ug Bleed 2
S-2P 5ug Bleed 2
RBD-8GS-I5350 50% 0.88ug Bleed 2
RBD-8GS-I5350 50% 5ug Bleed 2
RBD-8GS-I5350 100% 0.88ug Bleed 2
RBD-8GS-I5350 100% 5ug Bleed 2
RBD-12GS-I5350 50% 0.88ug Bleed 2
RBD-12GS-I5350 50% 5ug Bleed 2
RBD-12GS-I5350 100% 0.88ug Bleed 2
RBD-12GS-I5350 100% 5ug Bleed 2
RBD-16GS-I5350 50% 0.88ug Bleed 2
RBD-16GS-I5350 50% 5ug Bleed 2
RBD-16GS-I5350 100% 0.88ug Bleed 2
RBD-16GS-I5350 100% 5ug Bleed 2
Addavax
% Neutralization
Log10[Dilution-1]

KyMAb ELISA

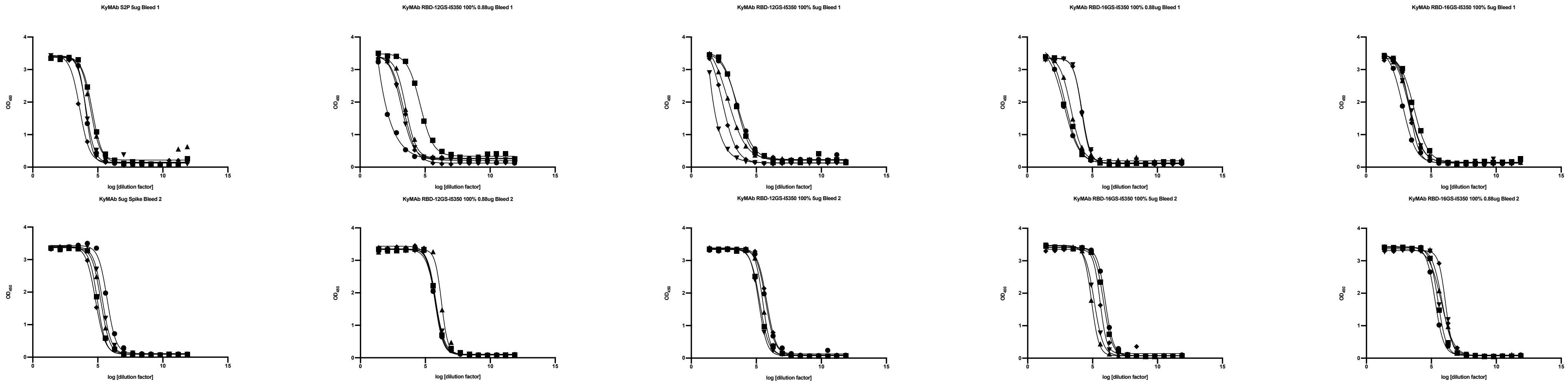

KyMAb S2P 5ug Bleed 1
KyMAb RBD-12GS-I5350 100% 0.88ug Bleed 1
KyMAb RBD-12GS-I5350 100% 5ug Bleed 1
KyMAb RBD-16GS-I5350 100% 0.88ug Bleed 1
KyMAb RBD-16GS-I5350 100% 5ug Bleed 1
KyMAb 5ug Spike Bleed 2
KyMAb RBD-12GS-I5350 100% 0.88ug Bleed 2
KyMAb RBD-12GS-I5350 100% 5ug Bleed 2
KyMAb RBD-16GS-I5350 100% 5ug Bleed 2
KyMAb RBD-16GS-I5350 100% 0.88ug Bleed 2
$OD_{450}$
log [dilution factor]

KyMAb Pseudoneutralization

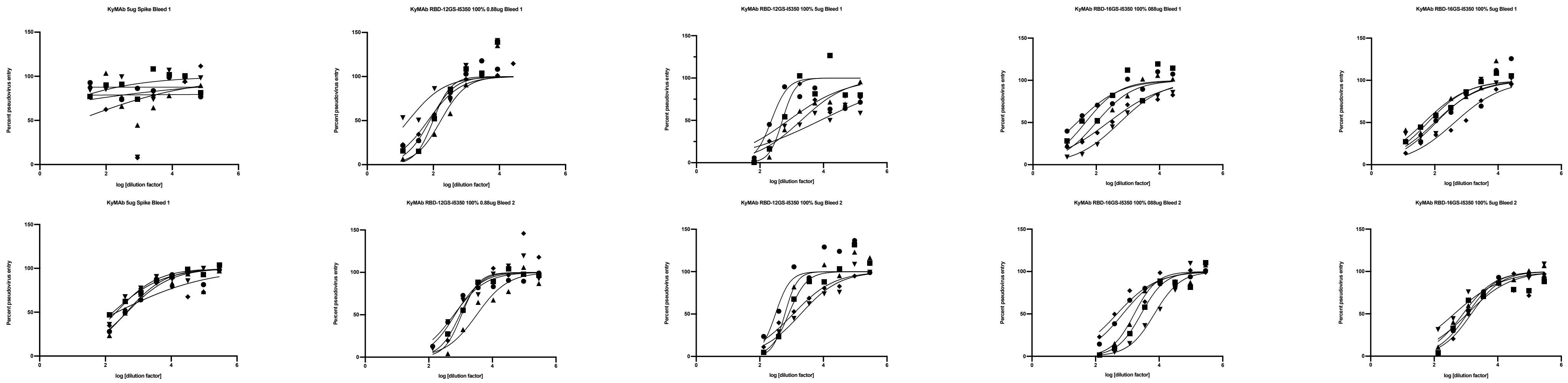
